# Evolution of the SARS-CoV-2 proteome in three dimensions (3D) during the first six months of the COVID-19 pandemic

**DOI:** 10.1101/2020.12.01.406637

**Authors:** Joseph H. Lubin, Christine Zardecki, Elliott M. Dolan, Changpeng Lu, Zhuofan Shen, Shuchismita Dutta, John D. Westbrook, Brian P. Hudson, David S. Goodsell, Jonathan K. Williams, Maria Voigt, Vidur Sarma, Lingjun Xie, Thejasvi Venkatachalam, Steven Arnold, Luz Helena Alfaro Alvarado, Kevin Catalfano, Aaliyah Khan, Erika McCarthy, Sophia Staggers, Brea Tinsley, Alan Trudeau, Jitendra Singh, Lindsey Whitmore, Helen Zheng, Matthew Benedek, Jenna Currier, Mark Dresel, Ashish Duvvuru, Britney Dyszel, Emily Fingar, Elizabeth M. Hennen, Michael Kirsch, Ali A. Khan, Charlotte Labrie-Cleary, Stephanie Laporte, Evan Lenkeit, Kailey Martin, Marilyn Orellana, Melanie Ortiz-Alvarez de la Campa, Isaac Paredes, Baleigh Wheeler, Allison Rupert, Andrew Sam, Katherine See, Santiago Soto Zapata, Paul A. Craig, Bonnie L. Hall, Jennifer Jiang, Julia R. Koeppe, Stephen A. Mills, Michael J. Pikaart, Rebecca Roberts, Yana Bromberg, J. Steen Hoyer, Siobain Duffy, Jay Tischfield, Francesc X. Ruiz, Eddy Arnold, Jean Baum, Jesse Sandberg, Grace Brannigan, Sagar D. Khare, Stephen K. Burley

**Affiliations:** Institute for Quantitative Biomedicine, Rutgers, The State University of New Jersey, Piscataway, NJ USA; Department of Chemistry and Chemical Biology, Rutgers, The State University of New Jersey, Piscataway, NJ USA; Research Collaboratory for Structural Bioinformatics Protein Data Bank, Rutgers, The State University of New Jersey, Piscataway, NJ USA; The Scripps Research Institute, La Jolla, CA USA; Rutgers Cancer Institute of New Jersey, Robert Wood Johnson Medical School, Rutgers, The State University of New Jersey, New Brunswick, NJ USA; Grinnell College, Grinnell, IA USA; University of Notre Dame, Notre Dame, IN USA; University of Maryland Baltimore County Baltimore, MD USA; Stevens Institute of Technology, Hoboken, NJ USA; Frostburg State University, Frostburg, MD USA; Youngstown State University, Youngstown, OH USA; University of Central Florida, Orlando, FL USA; New York City College of Technology, Brooklyn, NY USA; Howard University, Washington, DC USA; Watchung Hills Regional High School, Warren, NJ USA; Xavier University, Cincinnati, OH USA; Hope College, Holland, MI USA; Ursinus College, Collegeville, PA USA; SUNY Oswego, Oswego, NY USA; Roger Williams University, Bristol, RI USA; Brandeis University, Waltham, MA USA; University of Puerto Rico-Rio Piedras, San Juan, Puerto Rico; John Jay College, New York, NY USA; Grand View University, Des Moines, IA USA; Rochester Institute of Technology, Rochester, NY USA; Department of Biochemistry and Microbiology, Rutgers, The State University of New Jersey, New Brunswick, NJ USA; Department of Ecology, Evolution and Natural Resources, School of Environmental and Biological Sciences, Rutgers, The State University of New Jersey, New Brunswick, NJ USA; Department of Genetics, Rutgers, The State University of New Jersey, and Human Genetics Institute of New Jersey, Piscataway, NJ; Center for Advanced Biotechnology and Medicine, Rutgers, The State University of New Jersey, Piscataway, NJ USA; Center for Computational and Integrative Biology, Rutgers, The State University of New Jersey, Camden, NJ USA; Department of Physics, Rutgers, The State University of New Jersey, Camden, NJ USA; Research Collaboratory for Structural Bioinformatics Protein Data Bank, San Diego Supercomputer Center, University of California, San Diego, La Jolla, CA USA; Skaggs School of Pharmacy and Pharmaceutical Sciences, University of California, San Diego, La Jolla, CA USA

**Author notes:** Corresponding Authors:* Khare, S.D. and Burley, S.K.

## Abstract

Three-dimensional structures of SARS-CoV-2 and other coronaviral proteins archived in the Protein Data Bank were used to analyze viral proteome evolution during the first six months of the COVID-19 pandemic. Analyses of spatial locations, chemical properties, and structural and energetic impacts of the observed amino acid changes in >48,000 viral proteome sequences showed how each one of the 29 viral study proteins have undergone amino acid changes. Structural models computed for every unique sequence variant revealed that most substitutions map to protein surfaces and boundary layers with a minority affecting hydrophobic cores. Conservative changes were observed more frequently in cores *versus* boundary layers/surfaces. Active sites and protein-protein interfaces showed modest numbers of substitutions. Energetics calculations showed that the impact of substitutions on the thermodynamic stability of the proteome follows a universal bi-Gaussian distribution. Detailed results are presented for six drug discovery targets and four structural proteins comprising the virion, highlighting substitutions with the potential to impact protein structure, enzyme activity, and functional interfaces. Characterizing the evolution of the virus in three dimensions provides testable insights into viral protein function and should aid in structure-based drug discovery efforts as well as the prospective identification of amino acid substitutions with potential for drug resistance.

## Introduction

SARS-CoV-2, the causative agent of the COVID-19 global pandemic, is a member of the coronavirus family of RNA viruses that cause diseases in mammals and birds (Y. Chen, Liu, & Guo, 2020). The viral genome resembles a single-stranded cellular messenger RNA, ~29.9kb in length with a 7-methyl-G 5’ cap, a 3’ poly-A tail, and more than 10 open reading frames or Orfs (Figure 1). Viral proteins are expressed in two ways. Translation of two long polyproteins occurs initially, yielding the machinery required to copy the viral genome. Subsequent expression of multiple sub-genomic mRNAs produces the four structural proteins present in virions (see below) and other proteins designated as Orf3a, Orf6, Orf7a, Orf7b, Orf8, Orf9b, Orf14, and possibly the hypothetical protein Orf10. The non-structural proteins (nsps) are expressed within the shorter polyprotein 1a (pp1a, encompassing nsp1-nsp11) and the longer polyprotein 1ab (pp1ab, encompassing nsp1-nsp16). Both pp1a and pp1ab require two virally-encoded proteases for processing into individual nsp protomers (Figure 1). nsp3 includes a papain-like protease (PLPro) domain, which is responsible for polypeptide chain cleavage at three sites within the N-terminal portions of both polyproteins (dark blue inverted triangles in Figure 1). Ten additional polypeptide chain cleavages are carried out by nsp5 (light blue inverted triangles in Figure 1), also known as the main protease or the 3C-like protease. The structural proteins present in mature virions include the S-protein (surface spike glycoprotein, responsible for viral entry), the N-protein (nucleocapsid protein), the E-protein (a pentameric ion channel), and the M-protein (a second integral membrane protein found in the viral lipid bilayer).

**Figure 1.**
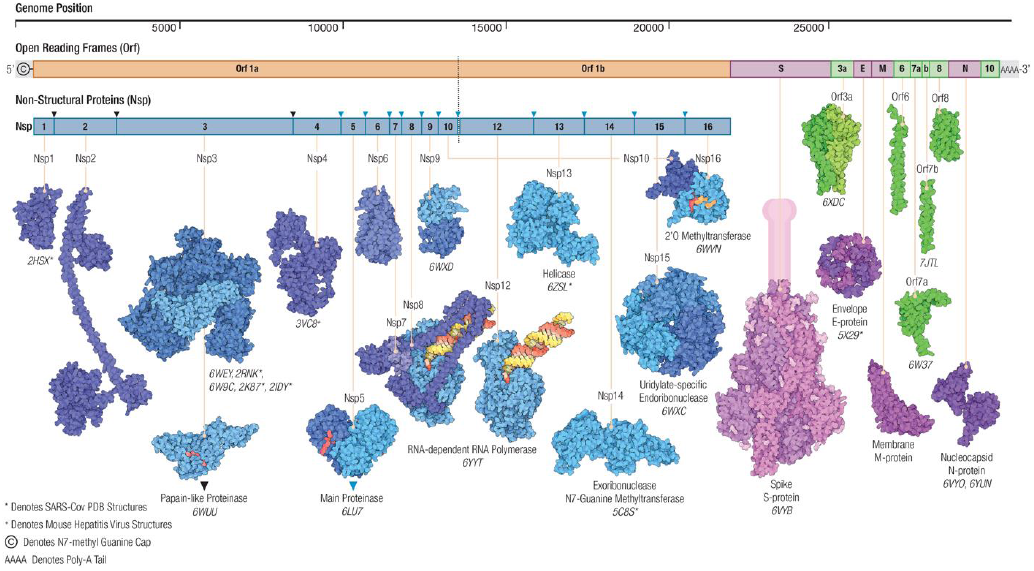
Architecture of the SARS-CoV-2 genome and proteome, including non-structural proteins derived from pp1a and pp1ab (nsps, shades of blue), virion structural proteins (pink/purple), and open reading frame proteins (Orfs, shades of green). Polyprotein cleavage sites are indicated by inverted triangles for Papain-like Proteinase (PLPro, black) and the Main Protease (nsp5, blue). The double-stranded RNA substrate-product complex of the RNA-dependent RNA polymerase (shown as the nsp7-nsp82-nsp12 heterotetramer and separately with only nsp12) is color coded (yellow: product strand, red: template strand). Transmembrane portions of the Spike S-protein are shown in cartoon form (pink).

Coronaviruses have the longest RNA virus genomes of all known single-stranded RNA viruses. Their RNA-dependent RNA polymerases (consisting of nsp7, two copies of nsp8, and nsp12) act together with RNA helicases (nsp13) and proofreading exonucleases (nsp14), to ensure efficient and relatively faithful copying of the lengthy genome (Denison, Graham, Donaldson, Eckerle, & Baric, 2011). Proofreading by nsp14 notwithstanding, coronavirus genome replication is not perfect, and coronaviruses do evolve as they passage serially from one host to the next. Today in the time of COVID-19, genome sequence-based “fingerprinting” of the virus in near real time during the pandemic has provided very detailed accounts of how the virus has moved around the globe since late 2019 as infected individuals, many of them asymptomatic, travelled from continent to continent (Hadfield et al., 2018; Wang, Hozumi, Yin, & Wei, 2020). Viral genome fingerprinting has also enabled detailed analyses of the impact of amino acid changes in particular proteins that modulate infectivity, *etc*. (*e.g*., (Korber et al., 2020)).

Herein, we report a comprehensive study of how the SARS-CoV-2 proteome has evolved in 3D during the first six months of the pandemic between late 2019 and June 25^th^ 2020. We combined viral genome sequence data assembled by GISAID (https://www.gisaid.org), the wealth of experimental 3D structure information for SARS-CoV-2 and other coronavirus proteins available from the open-access Protein Data Bank or PDB (Berman et al., 2000; Protein Data Bank, 1971; wwPDB consortium, 2019), and computed structural models in cases where experimentally-determined structures were not available.

The bulk of this work was initiated by research interns (undergraduates and one high school student) hosted virtually during the summer of 2020 by the Rutgers University Institute for Quantitative Biomedicine (IQB), the Rutgers University RISE Program, and the US-funded RCSB Protein Data Bank headquartered at Rutgers (Burley et al., 2018; Stephen K. Burley et al., 2020; Goodsell et al., 2020). Prior to the online five-week research program, participating students and mentors received one week of online training in 3D molecular visualization and computational bioinformatics in the IQB “Summer of the Coronaverse” Online Boot Camp (S. K. Burley et al., 2020). The methods used in the research study were developed, evaluated, and refined during the online Boot Camp. Supervision of the research phase was provided by IQB graduate students, postdoctoral fellows, and RCSB Protein Data Bank scientific staff, all of whom served as mentors in the Boot Camp. The research interns worked collaboratively in teams, carrying out multiple sequence alignments, constructing phylogenetic trees, computing 3D structural models of viral proteins, visualizing 3D structures, and analyzing the structural, functional, and energetic consequences of SARS-CoV-2 protein amino acid substitutions identified during the first six months of the pandemic. All computed 3D structural models and results of the sequence/energetics analyses are described in the main body of this paper and accompanying Supplementary Materials. The computed 3D structural models and energetics results are made freely available under Creative Commons license CC0 for researchers wishing to perform further computational and experimental studies (see https://iqb.rutgers.edu/covid-19_proteome_evolution).

## Results and Discussion

### Sequence Analyses

Viral genome sequencing and alignments of more than 48,000 individual isolates revealed protein sequence variation between December 2019 and late June 2020. We investigated the spatial locations, chemical properties, and structural and energetic impacts of the observed amino acid changes with reference to the original viral genome/proteome sequence publicly released in January 2020.

Every one of the 29 SARS-CoV-2 study proteins listed in Table 1 underwent changes in amino sequence, generating an average of approximately one unique sequence variant (USV) per study-protein amino acid residue (Lowest: nsp10 at ~0.59 USVs/residue; Highest: Orf3a at ~2.46 USVs/residue). Protein sequence differences were entirely restricted to non-synonymous changes in one or more residues. No insertions or deletions were detected in any of the 29 study proteins. Most USVs reflect a single amino acid change in the protein sequence (~66.8%). Smaller proportions of the USVs showed accumulation of two (~25.4%), three (~6.8%), four (~0.8%), or rarely five or more (~0.2%) amino acid substitutions. Where multiple substitutions were observed in a given study-protein USV, visual inspection of GISAID metadata typically revealed that they accumulated serially, but no systematic effort was made to track sequence changes as a function of sample collection date or geographic location. The modest degree of amino acid sequence variation observed for each of the 29 study proteins analyzed herein is consistent with previous studies of coronavirus evolution, which underscore the importance of the 3’-to-5’ exoribonuclease activity of nsp14 (reviewed in (Denison et al., 2011)). In contrast, RNA viruses that do not possess proofreading enzymes (*eg*., hepatitis C virus) exhibit significantly higher rates of amino acid substitution (Simmonds et al., 2005).

**Table 1.** Summary statistics from analysis of GISAID dataset (downloaded 06/25/2020).

| Study Protein | # Clean Protein Sequences Analyzed | # Clean Protein Sequences Unchanged | # Unique Protein Sequence Variants | Protein Length (residues) | Average # Unique Protein Sequence Variants (USV)/ Residue |
| --- | --- | --- | --- | --- | --- |
| nsp1 | 46414 | 45315 | 212 | 179 | 1.18 |
| nsp2 | 41579 | 28543 | 838 | 638 | 1.31 |
| nsp3a* | 37181 | 35364 | 223 | 206 | 1.08 |
| nsp3b* | 37181 | 36151 | 181 | 206 | 0.88 |
| nsp3c* | 37181 | 35665 | 229 | 332 | 0.69 |
| PLPro* | 37181 | 36133 | 225 | 343 | 0.66 |
| nsp3e* | 37181 | 36114 | 152 | 172 | 0.88 |
| UNK* | 37181 | 34614 | 455 | 686 | 0.66 |
| nsp4 | 45306 | 42803 | 380 | 500 | 0.76 |
| nsp5 | 46797 | 43884 | 217 | 306 | 0.71 |
| nsp6 | 46691 | 39758 | 262 | 290 | 0.90 |
| nsp7 | 48670 | 47876 | 68 | 83 | 0.83 |
| nsp8 | 48335 | 47635 | 144 | 198 | 0.73 |
| nsp9 | 48686 | 48289 | 82 | 113 | 0.73 |
| nsp10 | 46850 | 46507 | 81 | 139 | 0.59 |
| nsp12 | 44203 | 10266 | 730 | 932 | 0.78 |
| nsp13 | 44120 | 39652 | 466 | 595 | 0.79 |
| nsp14 | 31465 | 29600 | 335 | 527 | 0.64 |
| nsp15 | 42022 | 40208 | 326 | 346 | 0.94 |
| nsp16 | 42287 | 41118 | 206 | 298 | 0.69 |
| S-protein | 33290 | 7743 | 1190 | 1273 | 0.93 |
| Orf3a | 45932 | 27554 | 677 | 275 | 2.46 |
| E-protein | 48552 | 48052 | 82 | 75 | 1.09 |
| M-protein | 47326 | 45423 | 181 | 222 | 0.82 |
| Orf6 | 48490 | 47935 | 76 | 61 | 1.25 |
| Orf7a | 41969 | 41146 | 181 | 121 | 1.50 |
| Orf7b | 43211 | 42939 | 56 | 43 | 1.30 |
| Orf8 | 47796 | 42120 | 195 | 121 | 1.61 |
| N-protein | 45635 | 26486 | 889 | 419 | 2.12 |

\*part of nsp3

### Mapping Locations of Observed Sequence Variations in 3D

Experimental structures or computed 3D structural models were assembled for all 29 study proteins and their respective USVs (see Materials and Methods). For each study protein, we identified amino acid substitutions mapping to sites in the polypeptide chain buried in the hydrophobic core, exposed on the macromolecule surface, and present in the “boundary” layer between the core and the surface (Table 2). Not surprisingly, most of the amino acid substitutions occur on the protein surface (~46.2%) or within the boundary layer (~46.4%). Very few occur in the protein core (~7.4%). Characterization of the nature of each substitution (conserved, non-conserved) revealed that non-conservative amino acid changes were common, albeit less so if they occurred in the core (~54.0%) or the boundary layer (~55.0%), rather than on the protein surface (~69.4%). (N.B. A minority of USVs for some study proteins could not be modeled in 3D due to incomplete structural information.)

**Table 2.** Analysis results for 3D spatial locations and energetics of all 29 study-protein USVs. Column label definitions (left to right): Study Protein: study-protein or multiprotein-complex name. USVs: Total—number of USVs for each study protein identified across all GISAID sequences; Modeled—number of USVs for which 3D structural models were computed. 3D Mapping: layer identifications counted across all modeled substitutions. USV substitution count: number of single-substituted, double-substituted, *etc*. USVs. Substitutions: Total—number of unique substitutions identified across all study-protein USVs; Single USV—number of substitutions that occur in only one USV; Max Occurrences—number of USVs in which the most frequent substitution occurred (independent of GISAID count). Conservation: Conserved—number of conserved substitutions; Non-conserved—number of non-conserved substitutions. Energetic Impact: More Stable—number of USVs with ΔΔG^App^≤ 0; Less Stable—number of USVs with ΔΔG^App^ between 0 and two standard deviations above the mean value of <ΔΔG^App^>; Outlier—number of USVs with ΔΔG^App^ greater than two standard deviations above the mean value of <ΔΔG^App^>. <ΔΔG^App^> Stabilizing—average value of ΔΔG^App^ for all values of ΔΔG^App^≤ 0. <ΔΔG^App^> Destabilizing—average value of ΔΔG^App^ for all values of ΔΔG^App^>0 between 0.0 and two standard deviations above<ΔΔG^App^>. Standard deviations were computed using all ΔΔG^App^ values after excluding extreme outliers with ΔΔG^App^ greater than <ΔΔG^App^> plus four standard deviations.

| Study Protein | USVs | | 3D Mapping | | | USV Substitution Count | | | | | | Substitutions | | | Conservation | | Energetic Impact | | | Average $\Delta\Delta G^{\text{App}}$ | |
| --- | --- | --- | --- | --- | --- | --- | --- | --- | --- | --- | --- | --- | --- | --- | --- | --- | --- | --- | --- | --- | --- |
|  | Total | Modeled | Surface | Boundary | Core | 1 | 2 | 3 | 4 | 5 | 6+ | Total | Single USV | Max Occurrences | Conserved | Non-conserved | More Stable | Less Stable | Outlier | Stabilizing | Destabilizing |
| nsp1 | 212 | 212 | 118 | 92 | 15 | 200 | 11 | 1 | 0 | 0 | 0 | 211 | 200 | 5 | 67 | 144 | 44 | 153 | 17 | -1.9 (1.6) | 3.9 (2.9) |
| nsp2 | 838 | 838 | 640 | 578 | 67 | 489 | 260 | 81 | 7 | 1 | 0 | 639 | 456 | 205 | 219 | 420 | 195 | 608 | 38 | -2.1 (2.0) | 4.4 (3.4) |
| nsp3a* | 223 | 99 | 68 | 39 | 0 | 202 | 17 | 4 | 0 | 0 | 0 | 219 | 200 | 5 | 64 | 155 | 31 | 63 | 6 | -2.3 (2.5) | 3.1 (2.6) |
| nsp3b* | 181 | 142 | 68 | 58 | 21 | 170 | 11 | 0 | 0 | 0 | 0 | 174 | 158 | 3 | 66 | 108 | 25 | 107 | 11 | -1.5 (1.1) | 4.0 (3.2) |
| nsp3c* | 229 | 174 | 91 | 82 | 14 | 215 | 12 | 1 | 1 | 0 | 0 | 225 | 210 | 7 | 83 | 142 | 58 | 108 | 10 | -1.9 (1.7) | 2.8 (2.2) |
| PLPro* | 225 | 209 | 117 | 83 | 19 | 215 | 10 | 0 | 0 | 0 | 0 | 221 | 209 | 3 | 71 | 150 | 60 | 138 | 12 | -1.4 (1.3) | 3.2 (2.1) |
| nsp3e* | 152 | 89 | 66 | 26 | 3 | 145 | 5 | 2 | 0 | 0 | 0 | 147 | 140 | 7 | 50 | 97 | 29 | 56 | 5 | -1.8 (1.4) | 2.5 (1.6) |
| UNK* | 455 | 455 | 338 | 151 | 41 | 395 | 55 | 4 | 0 | 0 | 1 | 444 | 383 | 7 | 176 | 268 | 140 | 287 | 30 | -1.7 (1.5) | 3.7 (3.3) |
| nsp4 | 380 | 380 | 248 | 169 | 27 | 327 | 45 | 5 | 3 | 0 | 0 | 362 | 323 | 16 | 158 | 204 | 95 | 269 | 17 | -2.2 (2.1) | 3.1 (2.5) |
| nsp5 | 217 | 217 | 106 | 109 | 32 | 189 | 26 | 2 | 0 | 0 | 0 | 211 | 189 | 8 | 80 | 131 | 43 | 164 | 11 | -1.6 (1.6) | 3.9 (2.9) |
| nsp6 | 262 | 262 | 137 | 165 | 43 | 180 | 81 | 1 | 0 | 0 | 0 | 232 | 192 | 70 | 103 | 129 | 50 | 199 | 14 | -2.4 (1.9) | 4.3 (3.1) |
| nsp9 | 82 | 82 | 59 | 20 | 7 | 79 | 2 | 1 | 0 | 0 | 0 | 85 | 84 | 2 | 25 | 60 | 25 | 52 | 6 | -2.0 (1.5) | 3.1 (2.4) |
| nsp10-nsp16 | 286 | 269 | 128 | 124 | 37 | 266 | 19 | 1 | 0 | 0 | 0 | 282 | 260 | 4 | 105 | 177 | 66 | 191 | 13 | -1.8 (1.6) | 4.0 (2.7) |
| nsp7-nsp8<sub>2</sub>-nsp12 | 934 | 840 | 379 | 857 | 107 | 444 | 427 | 55 | 4 | 1 | 3 | 811 | 655 | 443 | 328 | 483 | 118 | 685 | 43 | -2.2 (2.6) | 7.6 (4.3) |
| nsp13 | 466 | 463 | 307 | 221 | 87 | 363 | 62 | 34 | 6 | 1 | 0 | 417 | 336 | 40 | 165 | 252 | 140 | 301 | 23 | -1.5 (1.6) | 2.4 (1.7) |
| nsp14 | 335 | 335 | 172 | 176 | 32 | 306 | 26 | 1 | 0 | 1 | 1 | 339 | 307 | 6 | 134 | 205 | 68 | 249 | 21 | -1.6 (1.3) | 3.9 (2.8) |
| nsp15 | 326 | 326 | 122 | 198 | 34 | 298 | 28 | 0 | 0 | 0 | 0 | 319 | 294 | 7 | 117 | 202 | 91 | 217 | 21 | -2.2 (2.3) | 3.7 (2.7) |
| Orf3a | 677 | 400 | 239 | 316 | 33 | 303 | 339 | 33 | 2 | 0 | 0 | 428 | 257 | 198 | 145 | 283 | 87 | 295 | 21 | -2.4 (2.2) | 6.4 (4.4) |
| Orf6 | 76 | 76 | 82 | 0 | 0 | 73 | 2 | 0 | 0 | 1 | 0 | 80 | 78 | 2 | 28 | 52 | 16 | 57 | 4 | -0.7 (0.5) | 1.5 (1.0) |
| Orf7a | 181 | 181 | 129 | 53 | 10 | 171 | 9 | 1 | 0 | 0 | 0 | 175 | 160 | 3 | 64 | 111 | 51 | 120 | 11 | -2.7 (3.0) | 3.7 (2.8) |
| Orf7b | 56 | 56 | 60 | 0 | 0 | 52 | 4 | 0 | 0 | 0 | 0 | 57 | 54 | 2 | 21 | 36 | 12 | 41 | 3 | -1.0 (0.8) | 2.4 (1.8) |
| Orf8 | 195 | 169 | 145 | 59 | 8 | 147 | 43 | 5 | 0 | 0 | 0 | 172 | 142 | 31 | 53 | 119 | 36 | 122 | 11 | -1.8 (1.2) | 4.5 (3.1) |
| E-protein | 82 | 56 | 38 | 20 | 1 | 79 | 3 | 0 | 0 | 0 | 0 | 81 | 78 | 3 | 41 | 40 | 18 | 37 | 1 | -2.2 (2.2) | 1.6 (1.4) |
| M-protein | 181 | 181 | 125 | 67 | 15 | 162 | 16 | 2 | 0 | 0 | 1 | 184 | 168 | 5 | 74 | 110 | 38 | 133 | 13 | -1.4 (1.2) | 3.0 (2.3) |
| N-protein | 889 | 262 | 205 | 79 | 9 | 429 | 185 | 237 | 36 | 2 | 0 | 577 | 344 | 272 | 132 | 445 | 74 | 176 | 14 | -1.3 (1.0) | 2.9 (2.3) |
| S-protein | 1190 | 689 | 348 | 824 | 72 | 327 | 675 | 171 | 13 | 3 | 1 | 922 | 652 | 805 | 312 | 610 | 157 | 496 | 38 | -1.8 (1.5) | 3.3 (2.6) |

\*part of nsp3

To further examine the types of amino acid changes in the viral proteome, we generated location-based substitution matrices from the observed USVs for each study protein and for the entire viral proteome (Figure 2). Substitutions to or from all 20 amino acids were observed across all 29 study proteins. Notable non-conservative changes include hydrophobic residues changing to negatively charged residues and *vice versa*, and glycine and proline residues changing to all types of amino acids on the surface, and to a lesser extent within the boundary layer. These trends reflect anticipated constraints imposed by protein structure on the thermodynamic stability due to amino acid substitutions. In the tightly packed environment of the hydrophobic core of a protein, fewer types of amino acid substitutions are likely to be compatible with the 3D structure, and changes that do not impair protein function are likely to be conservative. In contrast, protein boundary layers and surfaces impose far fewer constraints in terms of structural incompatibility and non-conservative substitutions.

**Figure 2.**
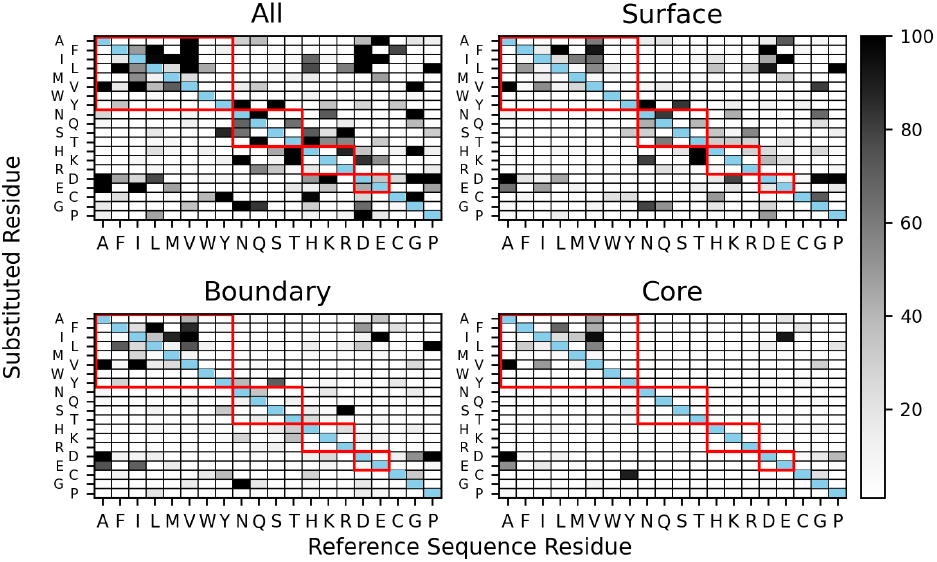
Observed frequencies for all USV substitutions of Reference Sequence Residue (*i.e*., original protein reference sequence amino acid) changing to Substituted Residue for all 29 study proteins considered together. Red boxes enclose conservative substitutions for hydrophobic, uncharged polar, positively charged, and negatively charged amino acids, respectively, in order from upper left to lower right. Cysteine, glycine, and proline are excluded from these groupings. Frequencies range from 0 (white) to 100 (black) for all, surface, and boundary substitutions. Frequencies range from 0 (white) to 50 (black) for core substitutions.

Most of the observed non-conservative changes can be attributed to the architecture of the genetic code and single base changes in the viral RNA genome. For example, Alanine to Aspartic and Glutamic acid changes are achievable *via* single base changes in the second base of their respective codons. However, changes requiring double base changes (*eg*., Proline to Aspartate) were also observed.

### Analyzing Energetic Consequences of Observed Sequence Variations

The energetic impact of observed amino acid substitutions for each unique sequence variant of each study protein was calculated using Rosetta (Table 2, Figure 3). A majority of the amino acid changes were estimated to be moderately destabilizing as judged by changes in the free energy of stabilization (apparent ΔΔG or ΔΔG^App^ =0.0 to +15.0 Rosetta energy units or REU; ~72.0%). A modest number were estimated to be stabilizing (ΔΔG^App^=−0.01 to −15.0 REU; ~23.7%). In the minority of cases, ΔΔG^App^ exceeded +15.0 REU (~4.3%). The distribution of ΔΔG^App^ values was used to identify outliers for each study protein (Table 2).

**Figure 3.**
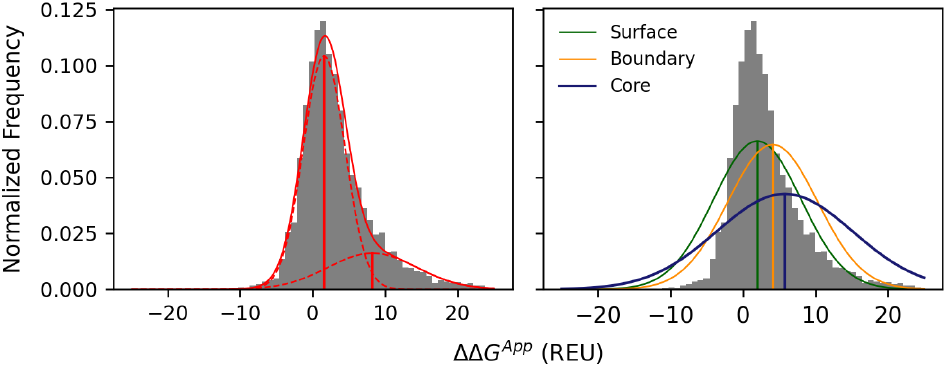
Normalized frequency histogram for ΔΔG^App^ calculated for all USVs aggregated across all 29 study proteins. Left: Overlay with fitted bi-Gaussian curve (solid red line, R^2^=0.95), with fitted individual Gaussian curves (dashed red lines). The means for the individual Gaussian distributions were +1.6 REU (standard deviation or SD: 8.3) and +8.2 REU (SD: 37.6). Right: Overlay of the same normalized frequency histogram with fitted single Gaussian curves fitted to subsets of USVs with Surface (green; mean value: +1.6 REU, SD: 12.2; R^2^=0.86), Boundary (yellow; mean value: +3.7 REU, SD: 24.3; R^2^=0.79), or Core (blue; mean value: +5.1 REU, SD: 42.9; R^2^=0.45) substitutions. USVs with multiple substitutions were included in single Gaussian fitting when all substitutions mapped to the same region of the study protein.

Given that all modeled amino acid substitutions were detected in viruses that likely had infected human hosts when they were isolated, we assume that all modeled USVs correspond to stable, functional proteins. Most globular proteins are marginally stable, with measured free energies of stabilization ΔG~−5 to −15 kcal/mol (Privalov & Gill, 1988), and tolerated amino acid substitutions are expected to have an impact within this range. Therefore, we believe that the small minority of computed large positive ΔΔG^App^ values represent artifacts arising from errors/approximations in our calculations (Table 2). For example, positional restraints on backbone atoms were employed when modeling USVs in Rosetta to prevent substantial departures from the reference protein backbone conformation so more permissive restraints on the polypeptide chain backbone may be required to model computationally the effects of some particularly large amino acid changes. Alternatively, large positive values of ΔΔG^App^ may reflect shortcomings in the Rosetta energy function. Outlier cases provide a benchmark for improvements in Rosetta and other stability calculation approaches. Outliers notwithstanding, ~95% of all computationally modeled USVs yielded reasonable ΔΔG^App^ values.

We next examined the distribution of energetic effects of the observed substitutions for each study protein and aggregated across all 29 viral study proteins. Several previously published experimental and theoretical studies have examined the distributions of thermodynamic stability changes due to point substitutions in individual proteins, and examined the implication of these distributions for molecular evolution (Bloom et al., 2005; Faure & Koonin, 2015; Razban & Shakhnovich, 2020; Tokuriki, Stricher, Schymkowitz, Serrano, & Tawfik, 2007). Our dataset provides an opportunity to re-examine conclusions from these studies which are, with a single exception (Nisthal, Wang, Ary, & Mayo, 2019), based on limited experimental data and/or computational findings. Tokuriki *et al* (2007) used FoldX-based calculations of all single substitutions in 21 different globular proteins and found that despite a diverse range of sizes and folds, the distribution of stability effects largely follows a bi-Gaussian function for each protein. They found that surface residues exhibit a narrow distribution with a modestly destabilizing mean ΔΔG^App^ (<ΔΔG^App^>), whereas core residues exhibit a wider distribution with higher positive <ΔΔG^App^> values (Tokuriki et al., 2007). Such asymmetric distributions were also found for lattice model proteins, and were recently shown to arise from first-principle statistical mechanical considerations and a sufficiently large amino acid alphabet size (Razban & Shakhnovich, 2020). Faure and Koonin (2015) obtained similar distributions across proteomes of five organisms selected from archaea, prokaryota, and eukaryota, suggesting that this distribution of energetic effects is a universal and evolutionarily conserved feature of globular protein folds (Faure & Koonin, 2015).

In contrast with larger and more comprehensive datasets used in previous work (all substitutions at all sites in a protein), approximately one substitution per residue per study protein was sampled in the SARS-CoV-2 dataset downloaded from GISAID. To investigate whether or not the observed stability effects follow a similar distribution, we fit bi-Gaussian models to ΔΔG^App^ histograms for all USVs for all 29 proteins (Figure 3). The bi-Gaussian distribution fits the calculated stability distributions better than a single Gaussian (R^2^=0.95 for a bi-Gaussian and R^2^=0.80 for a single Gaussian). Individual Gaussian peaks correspond closely to the energetic impacts of surface and core substitutions, respectively (Figure 3). This trend was observed for both types of Rosetta-based stability calculations, including those in which a dampened repulsive van der Waals potential was used during the rotamer optimization step. For each calculation type, the mean destabilization calculated for the core substitution distribution is smaller than the mean value associated with the second Gaussian peak observed in the full set of substitutions, possibly due to contributions to the second peak from destabilizing boundary layer substitutions that shift the mean to higher values (and possibly to limitations of the sampling and scoring approach discussed above). Bi-Gaussian fits to ΔΔG^App^ distributions for each of the 29 study proteins considered individually (Supplementary Table Gaussian) show similarly good fits for bi-Gaussian functions for globular study proteins. Robustness with respect to destabilizing effects of amino acid changes both limits and promotes viral evolution. It is, therefore, remarkable that the observed variation in the SARS-CoV-2 proteome over the first six months of the pandemic follows this universal trend, speaking perhaps to the relative rapidity of viral evolution due to large population sizes and imperfect replication machinery.

### Analyses of Study Proteins

The sections that follow provide more detailed results and discussion pertaining to USVs identified for 13 of the 29 SARS-CoV-2 study proteins, including one validated drug target [RNA-dependent RNA polymerase (RdRp, nsp7/nsp82/nsp12 heterotetramer)], five potential small-molecule drug discovery targets [papain-like proteinase (PLPro, part of nsp3), main protease (nsp5), RNA helicase (nsp13), proofreading exoribonuclease (nsp14), and methyltransferase (nsp10/nsp16 heterodimer)], plus the four structural proteins comprising the virion [spike S-protein, nucleocapsid N-protein, pentameric ion channel E-protein, and integral membrane M-protein]. Analysis results obtained for USVs of the remaining study proteins are provided in Supplementary Materials together with additional information regarding all 29 study proteins.

### Non-structural Proteins 7, 8, and 12 (nsp7/nsp8_2_/nsp12)

The RNA-dependent RNA polymerase (RdRp) is a macromolecular machine made up of four protomers, including nsp7, two asymmetrically bound copies of nsp8, and the catalytic subunit nsp12. The resulting heterotetramer is responsible for copying the RNA genome and generating nine subgenomic RNAs (D. Kim et al., 2020). nsp12 consists of three globular domains: an N-terminal nidovirus RdRp-associated nucleotidyltransferase (NiRAN), an interface domain, and a C-terminal RdRp domain. The active site of nsp12 includes residues Thr611 to Met626 (TPHLMGWDYPKCDRAM) comprising Motif A (Y. Gao et al., 2020). nsp12 binds to one turn of double-stranded RNA, and residues D760 and D761 bind to the 3’ end of the RNA and are essential for RNA synthesis (Hillen et al., 2020). The RNA duplex is flanked by α-helical arms formed by N-terminal segments of the two nsp8 protomers, which appear to grip the RNA and prevent its premature dissociation from the RdRp (*i.e*., confer processivity). Positively-charged residues of nsp8 occurring within the RdRp-RNA interface include K36, K37, K39, K40, K46, R51, R57, K58, and K61. Of these, K58 interacts with the RNA duplex emerging from the active site. Any change of this residue in nsp8 yields a replication-incompetent virus (Hillen et al., 2020). Since deposition of PDB ID 6M71 (Y. Gao et al., 2020), a plethora of RdRp structures has become available from the PDB.

Following US Food and Drug Administration (FDA) approval for remdesivir, RdRp can be regarded as being a validated drug target for treatment of SARS-CoV-2-infected individuals. Structures of SARS-CoV-2 RdRp containing incorporated remdesivir (PDB ID 7BV2 (Yin et al., 2020) and PDB ID 7C2K (Q. Wang, J. Wu, et al., 2020)) help explain the drug’s mechanism of action *via* delayed-chain termination (Gordon et al., 2020) and provide a valuable starting point for design of second-generation RdRp inhibitors that are more potent and more selective and possibly orally bioavailable. Residues K545, R553, D623, S682, T687, N691, S759, D760, and D761 in nsp12 interact directly with remdesivir (Yin et al., 2020), while S861 may be involved in a steric clash with the 1’-CN group of remdesivir, possibly perturbing the position of the RNA duplex (Q. Wang, J. Wu, et al., 2020). Knowledge of the structures of remdesivir-RdRp complexes will also provide valuable insights into potential sources of drug resistance.

The experimental structure of the RdRp-duplex RNA complex (PDB ID 6YYT (Hillen et al., 2020)) was used for evolutionary analyses of nsp7, nsp8, and nsp12 (Fig. RdRp A and B). Each protomer is considered in turn below.

#### nsp7

Sequencing of 48,670 viral genomes identified 47,876 unchanged sequences and 68 USVs of nsp7 *versus* the reference sequence, with 66 single and two double substitutions (Tables 1 and 2). Most substitutions occurred in only one USV (~91%). The most frequently observed USV for nsp7 (S25L; non-conservative, surface) was detected 562 times in the GISAID dataset.

#### nsp8

Sequencing of 48,335 viral genomes identified 47,635 unchanged sequences and 144 USVs of nsp8 *versus* the reference protein sequence, with 140 single, two double, one triple, and one quintuple substitutions (Tables 1 and 2). Most substitutions occurred in only one USV (~99%). The most frequently observed USV for nsp8 (M129I; conservative, core) was detected 124 times in the GISAID dataset. No substitutions of the essential RNA-binding residue K58 were observed. Of the remaining eight positively-charged residues that face the RNA duplex, substitutions were observed for five, including K37, K40, R51, R57, and K61 (both R51L and R57L preclude salt bridge formation with RNA). Substitutions of R51 were observed in 3 different USVs, occurring as three distinct substitutions (R51L, R51C, R51H). Another interesting nsp8 USV is the singly-observed quintuple substitution USV occurring within the N-terminal arm (A74S/S76C/A81S/V83L/S85M). This USV may be the result of a sequencing artifact, as none of the five substitutions were observed in any other USV. One other USV exhibits adjacent amino acid changes: M90S/L91F. This pair of residues occurs at the interface with nsp12 for one nsp8 protomer and near a shared interface with nsp7 and nsp12 in the other copy.

#### nsp12

Sequencing of 44,203 viral genomes identified 10,266 unchanged sequences and 730 USVs of nsp12 *versus* the reference sequence, with 249 single, 424 double, 51 triple, 3 quadruple, and 3 multi-point substitutions (Tables 1 and 2). A majority of substitutions occurred in only one USV (~74%). More than 97% (count~32,000) of the ~44,000 GISAID dataset nsp12 sequences differing from the reference sequence carried the same P323L substitution. This substitution constitutes a distinct nsp12 clade that was first detected in the United Kingdom in January 2020 and subsequently in many other countries around the world.

Approximately 61% of the observed amino acid substitutions were non-conservative (364 non-conservative *versus* 228 conservative), with most of the non-conservative changes occurring in the boundary and surface portions of the 3D structure. (N.B.: Only 60 point substitutions map to the protein core.) Two of the multi-point substitutions (A97V/S520I/E522D/D523Y/A529S/L829I and T85S/I201F/V202F/V330E/I333T) were observed only once. In both cases, all substitutions were unique to that particular USV, suggesting that they are both the result of sequencing artifacts.

#### nsp7/nsp8_2_/nsp12 Energetics

The vast majority of the USVs (83%) were estimated to be moderately less stable than the reference sequence (<ΔΔG^App^>~+7.6 REU). In fewer than 4% cases, the estimated change in apparent free energy of stabilization change exceeded + 19.1 REU. A minority of the USVs (~13%) were estimated to be more stable than the reference sequence (<ΔΔG^App^>~−2.2 REU). (N.B.: Hereafter, references will be made to Tables 1 and 2 to avoid repeating the same text summarizing amino acid substitutions and energetics analyses for each of the remaining study proteins.)

#### nsp12 Active Site

Of the residues in active site Motif A (Fig. RdRp C), substitutions were observed in residues H613, L614, M615, W617, Y619, and A625 (Fig. RdRp C). It is remarkable that all of these residues are oriented toward the hydrophobic core of the protein, away from the active site, and should, therefore, not disrupt catalysis. No substitutions were observed for nsp12 residues that interact directly or *via* bridging water molecules with remdesivir (Fig. RdRp C; K545, R553, D623, N691, D760, S759, D760).

#### Protein-Protein Interfaces

The four protomers forming the RdRp heterotetramer bury significant numbers of residues within the various protein-protein interfaces. It is, therefore, difficult to be certain that a distal substitution might not have a steric influence on one or more of these interfaces. Below, we enumerate substitutions with the potential for direct effects on interfacial contacts.

Eleven substitutions involving the following six nsp7 residues could affect binding to nsp12: K7, L14, S15, S26, L40, and L41. Seven of these 11 substitutions were conservative. nsp12 substitutions at the following sites could affect binding to nsp7: T409, P412, F415, Y420, E436, A443, and D445. Y420S would break an observed hydrogen bond with D5 of nsp7. E436G/K would break an observed salt bridge with K43. Many of the nsp7 and nsp12 substitutions occurring within their contact interface were highly destabilizing, with seven giving ΔΔG^App^> + 10 REU.

nsp7 makes minimal contact with one copy of nsp8. Observed nsp7 substitutions at residues S25 (S25L) and S26 (S26A and S26F) involve exchange of serine for a hydrophobic residue. Both substitutions at S26 break an observed hydrogen bond with D163 of nsp8. No nsp8 D163 substitutions were identified.

The contact surface of nsp7 with the second copy of nsp8 is more extensive than with the first. nsp7 substitutions occurring within this inter-subunit interface include residues V6, T9, S15, V16, L20, L28, Q31, F49, E50, M52, S54, L56, S57, V58, L60, S61, V66, I68, and L71 (17/27 substitutions affecting all 19 nsp7 residues were conservative). S54P is a noteworthy amino acid change that inserts a Proline into the middle of an interfacial α-helix. Substitutions of the following nsp8 residues may affect binding to nsp7: residues V83, T84, S85, T89, M90, L91, M94, L95, N100, A102, I107, V115, P116, I119, L122, V131, and A150 (14 of the 21 substitutions involving these 17 sites were conservative).

Because the two nsp8 chains occur in asymmetric environments, a given substitution may alter one interface or the other, or both. Substitutions at 23 sites could affect the nsp8-nsp12 interface for one of the chains (T84, A86, L91, L95, N104, I107, V115, P116, I119, P121, L122, T123, K127, M129, V131, I132, P133, T141, A150, W154, V160, W182, and T187). Substitutions at five sites (T68, K72, R75, S76, and K79) could affect only the nsp8-nsp12 interface with the chain that wraps around nsp7. Substitutions at three sites (V83, M90, and M94) could affect both interfaces. Of these 38 substitutions across 31 sites, 19 were conservative. A P121S substitution in nsp8 could give rise to a backbone hydrogen bond with V398 of nsp12. Two Tryptophan to Cysteine substitutions (W154C and W182C) occurring in nsp8 were extreme outliers with ΔΔG^App^>+30 REU, suggesting that some backbone rearrangement is necessary in response to exchange of the large Tryptophan side chains for smaller Cysteines.

In nsp12, substitutions of 25 residues could affect the interface with the first nsp8 protomer (L270, P323, T324, P328, L329, V330, V338, F340, P378, A379, M380, A382, A383, N386, V398, A399, V405, F407, W509, L514, S518, M519, S520, D523, and V675). Substitutions of 10 residues in nsp12 could affect the interface formed with the second copy of nsp8 (N414, F415, D846, I847, V848, T850, M899, M902, M906, T908). No nsp12 substitutions appear to affect contacts with both copies of nsp8. Of the 50 observed nsp12 substitutions occurring at 35 sites, 26 were conservative. The clade-defining nsp12 P323L substitution occurs at the C-terminus of an α-helix within the smaller interface between nsp12 and the first nsp8 protomer. While the structural consequences of this P→L substitution appear negligible, the computed ΔΔGApp ~8 REU. This apparent discrepancy almost certainly reflects limitations in the Rosetta energetics calculation.

Figure RdRp. (A) Space-filling representation of the experimental structure of the nsp7/nsp82/nsp12 heterotetramer bound to double-stranded RNA (PDB ID 6YYT (Hillen et al., 2020)) viewed into the enzyme active site on the anterior surface of nsp12. (B) Identical view of PDB ID 6YYT with nsp7 and nsp8 removed to reveal interactions of nsp12 with RNA. Protein color coding: nsp12-light blue; nsp8-dark blue; nsp7-blue/grey; RNA color coding: template strand-shades of red; product strand-shades of yellow. (C) Ribbon/atomic stick figure representation of the active site of nsp12 (PDB ID 7BV2 (Yin et al., 2020); mostly grey) occupied by the RNA template:product duplex (backbone shown as tubes, bases shown as sticks, colored in shades of orange) with remdesivir (shown as an atomic stick figure following enzymatic incorporation into the RNA product strand; atom color coding: C-green, N-blue, C-red, S-yellow). The active site Motif A is colored coded magenta (atom color coding for invariant residues: C-magenta, N-blue, O-dark red) and purple (atom color coding for substituted residues: C-purple, N-blue, O-dark red, S-yellow). Residues making direct or water mediated contacts with remdesivir are colored light red (atom color coding: C-light red, N-blue, O-dark red, S-yellow).

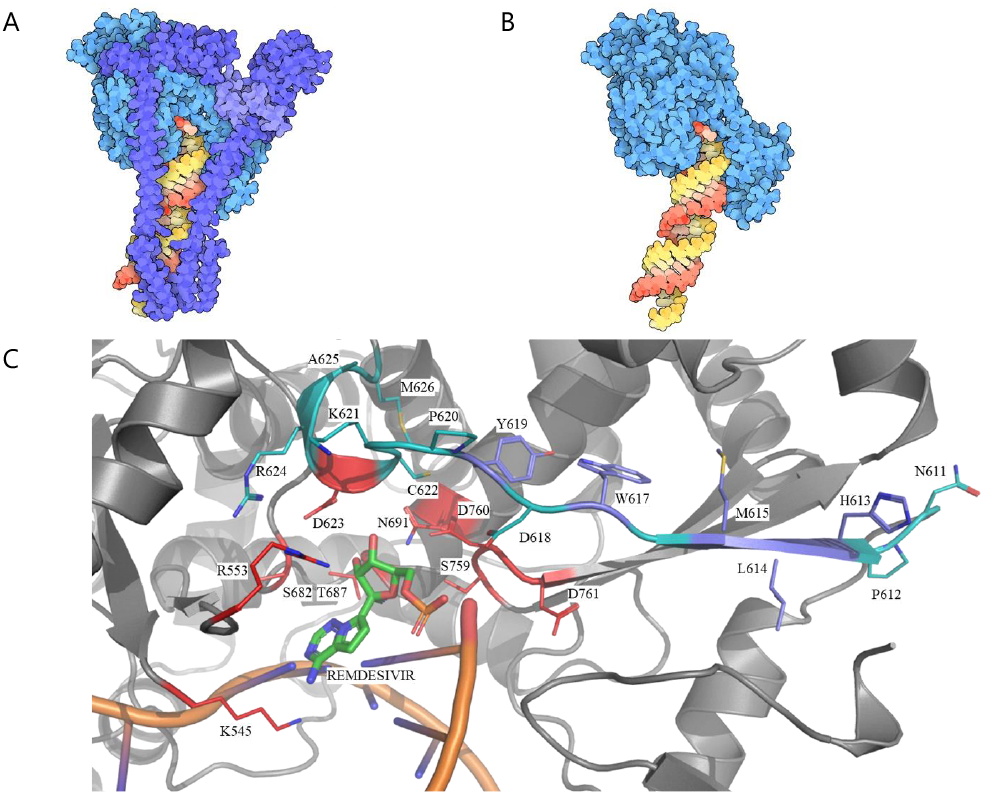

### Non-structural Protein 3 Papain-like Proteinase (PLPro)

The papain-like proteinase (PLPro) is a 343-residue segment occurring within the 1945 residue multi-domain protein nsp3. It is one of two viral proteases responsible for processing of the polyprotein products of translation of the viral genome following infection. This enzyme cleaves the polyproteins pp1a and pp1ab at three sites (black inverted triangles in Figure 1): the nsp1/nsp2 junction and its own N- and C-termini. These three cleavage events liberate nsp1, nsp2, and nsp3. The PLPro portion of nsp3 is also implicated in cleaving post-translational modifications of ubiquitin (Ubl) and ISG15 domains of host proteins as an evasion mechanism against host antiviral immune responses (Clementz et al., 2010).

PLPro is a cytoplasmic cysteine endopeptidase (EC 3.4.22.69) that catalyzes cleavage of the peptide bond C-terminal to LXGG motifs (where X is any amino acid) in the viral polyproteins. This enzyme also recognizes conserved LRGG motifs found within the C-terminal segments of Ubl and ISG15 proteins. According to the MEROPS classification, PLPro belongs to the peptidase clan CA (family C16), containing a Cys-His-Asp catalytic triad (C111–H272-D286). The first structure of SARS-CoV-2 PLPro to be made public (PDB ID 6W9C (Michalska, 2020)) revealed a symmetric homotrimer with each enzyme monomer being highly similar to that of SARS-CoV-1 PLPro (PDB ID 2FE8 (Ratia et al., 2006); root mean square deviation or r.m.s.d.~0.8Å, sequence identity~83%). Since PDB release of this initial SARS-CoV-2 PLPro structure, additional co-crystal structures of PLPro with a variety of ligands have been deposited to the PDB (list updated weekly at http://rcsb.org/covid19). In many of these structures the enzyme is monomeric, indicating that the trimer observed in PDB ID 6W9C is almost certainly a crystal packing artifact. Comparison of the various PLPro monomer structures reveals that the enzyme does not undergo large conformational changes upon binding of inhibitors or (protein) substrates (Fig. PLPro A). We, therefore, used the structure of an inhibited form of the enzyme (PDB ID 6WUU (Rut et al., 2020)) for evolutionary analyses of PLPro (Fig. PLPro A).

Overall substitution trends for PLPro and energetics analysis results are summarized in Tables 1 and 2. P1640L (non-conservative, surface) and T1626I (non-conservative, surface) are the two most common USVs, observed in 48 and 47 GISAID dataset sequences, respectively. No amino acid substitutions were identified in the enzyme active site – the catalytic triad is fully preserved in all observed USVs. However, examination of apo- and inhibitor/substrate-bound structures indicates that several substitutions occur in the ISG15- and ubiquitin-binding regions of PLPro. These substitutions (*eg*., F1632S, D1624G, D1625H, S1633G) mapping to the S2 and S4 α-helices of PLPro (Fig. PLPro B) may alter the binding affinity and specificity of PLPro for interactions with host protein substrates. In cell-based assays, the interactome of SARS-CoV-2 PLPro appears to be significantly different from that of SARS-CoV-1 PLPro. SARS-CoV-2 PLPro prefers ISG15 binding to Ubl whereas SARS-CoV-1 PLPro prefers Ubl binding to ISG15 (Shin et al., 2020). The S2 and S4 regions are interaction hotspots in the interfaces of PLPro with ISG15 and Ubl. Amino acid changes in these regions may change the protein’s interactome. Finally, two observed substitutions affecting active-site proximal proline residues P1810S and P1811S may affect inhibitor binding and represent potential sites of drug resistance mutations (Fig. PLPro C).

Figure PLPro. (A) Space-filling representation of the experimental structure of the PLPro monomer (blue) bound to a covalent inhibitor (Vir250; red/pink) (PDB ID 6WUU (Rut et al., 2020)). (B) Ribbon/atomic stick figure representation of the PLPro-ISG15 interface (ISG15: gray; atom color coding: C-grey, N-blue, O-red; unchanged PLPro residues: cyan, atom color coding: C-cyan, N-blue, O-red; and substituted PLPro residues: purple, atom color coding: C-purple, N-blue, O-red, S-yellow). (C) Ribbon/atomic stick figure representation of PLPro active site (color coding as for Fig. PLPro B) occupied by a non-covalent inhibitor (GRL0617) shown as an atomic stick figure (atom color coding: C-green, N-blue, O-red, H-bonds-dotted yellow lines; PDB ID 7CMD (X. Gao et al., 2020)).

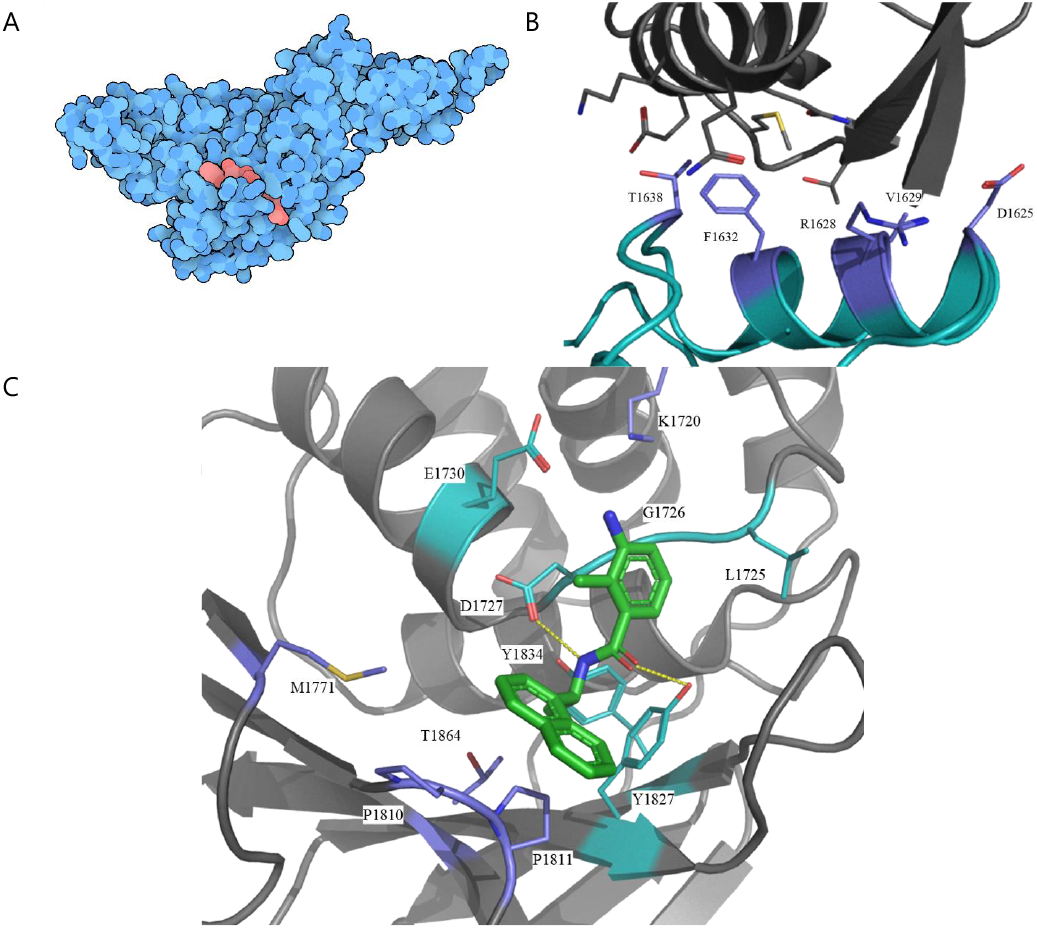

### Non-structural Protein 5 Main Protease (nsp5)

nsp5 is the other viral protease responsible for processing the viral polyproteins (synonyms: main protease, 3CL protease). This enzyme cleaves the longer polyprotein pp1ab at 11 sites (light blue inverted triangles in Figure 1), beginning with liberation of its own N-terminus and concluding with separation of nsp15 from nsp16 near the C-terminus of the polyprotein. nsp5 is a 306-residue cysteine endopeptidase (EC 3.4.22.69) that catalyzes cleavage of sites similar to TSAVLQ/SGFRK (where / denotes the cleavage site). Conserved residues Histidine 41 (H41) and Cysteine 145 (C145) constitute the catalytic dyad (Huang, Wei, Fan, Liu, & Lai, 2004). The first structure of SARS-CoV-2 nsp5 deposited into the PDB (PDB ID 6LU7 (Jin et al., 2020); Figure nsp5) revealed a symmetric homodimeric structure extremely similar to that of its SARS-CoV-1 homolog (r.m.s.d.~0.8Å, sequence identity>95% with PDB ID 1Q2W (Pollack, 2003)). Since PDB release of this initial nsp5 structure, ~200 co-crystal structures of nsp5 with a variety of small chemical fragments and larger ligands have been deposited to the PDB (updated weekly at http://rcsb.org/covid19). Open access to this wealth of structural information spurred the launch of an international COVID-19 Moonshot effort to discover and develop drug-like inhibitors (Chodera, Lee, London, & von Delft, 2020). The inaugural nsp5 structure (PDB ID 6LU7) was used for the evolutionary analyses that follow (Fig. nsp5 A).

Overall substitution trends for nsp5 and energetics analysis results are summarized in Tables 1 and 2. G15S (non-conservative, boundary) is the most common USV, observed in 1082 sequences. The most striking change observed in the GISAID dataset involves H41, the catalytic Histidine (Fig. nsp5 B, shown in red) substitution of which is expected to eliminate catalytic activity. This substitution was detected in the H41P/L50H double substitution. It is possible that loss of H41 has been compensated by the L50H substitution, though the distance between L50 and the active site (L50:C*α*-C145:C*α* ~16Å *versus* H41:C*α*-C145:C*α*~7Å) would require significant backbone rearrangement. Only one viral genome with this USV was detected in the GISAID dataset, which raises the possibility that it represents a sequencing artifact. No other observed USVs included substitutions of residue L50 to Histidine, but other amino acid changes at that site were observed within the GISAID dataset. Experimental characterization of the enzymatic activity of the H41P;L50H double substitution would resolve the issue.

Several amino acids within or adjacent to the substrate binding groove underwent substitutions (Fig. nsp5 B, shown in purple) that may affect substrate binding, including T25, M49, M165, E166, 168, 188, 189, and A191. The most dramatic alteration to the active site occurs in the triple substitution M165L;E166V;A191E. E166 lines the active site cleft, where it is thought to form a hydrogen bond with the pre-scissile residue of the substrate. The same residue also appears to interact with the N-terminus of the homodimeric partner. Each of these substitutions is unique to a single USV, occurring only once in the GISAID dataset. Other substitutions were observed at residues 165 and 191 in other USVs.

A number of residues occurring near the dimerization interface were also substituted, including residues M6, A7, G71, A116, S121, V125, G170, G215, M276, G278, S284, A285, Q299, G302, and T304, any one of which could affect dimerization. In several cases, Glycine residues were substituted for larger hydrogen-bonding residues (even when such substitutions appear to be destabilizing: G71S, G170R, G215R, G278R). While total stability was reduced, dimeric stability was likely increased, consistent with the expected biological importance of strong dimerization. Interesting, all substitutions mapping to the dimer interface occurred in USVs lacking any other substitutions.

Finally, there were four cases in which substitutions to Proline (a helix breaking amino acid) occurred at positions falling within α-helical or β-strand secondary structural elements (K90P, S123P, A206P, S301P). The latter three represent the most extreme energetic outliers of all USVs, and all four were observed only once in the GISAID dataset. S123P occurs at the end of a β-strand at the dimeric interface near the C-terminus of the homodimeric partner. The observed energetic consequences of these substitutions introducing Proline residues suggest the potential for significant structural perturbation.

Figure nsp5. (A) Space-filling representation of the experimental structure of the nsp5 homodimer covalently bound to a substrate analogue inhibitor (PDB ID 6LU7; (Jin et al., 2020)). Color Coding: nsp5 monomers-light and dark blue; substrate analogue PRD_002214 (https://www.rcsb.org/ligand/PRD_002214)-red. (B) Ribbon/atomic stick figure representation of the active site of nsp5 (grey) occupied by PRD_002214 covalently bound to C145 (atom color coding: C-green, N-blue, O-red). Catalytic residues H41 and C145 denoted with red ribbon and atomic stick figure sidechains (atom color coding: C-light red, N-blue, S-yellow). Substituted active site residues denoted with purple ribbon and atomic stick figures (atom color coding: C-purple, N-blue, O-red, S-yellow).

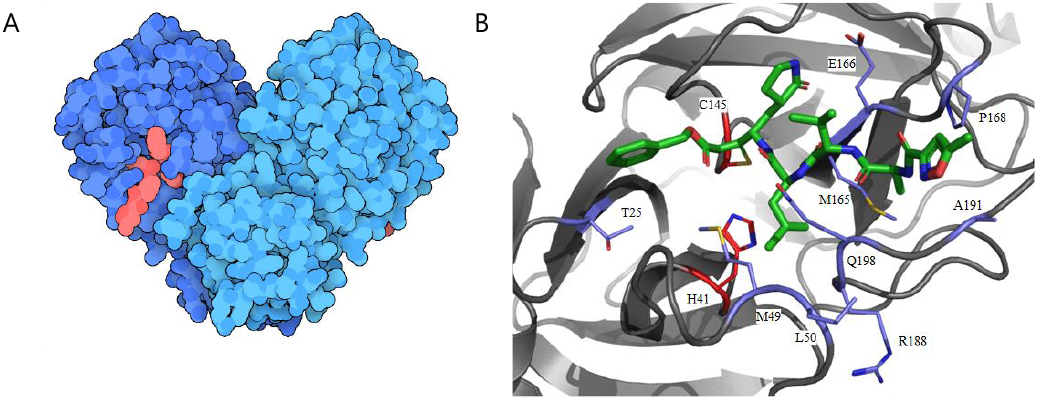

### Non-structural protein 13 (nsp13)

nsp13 plays a central role in viral replication by unwinding RNA secondary structure within the 5’ untranslated region of the genome (Miao, Tidu, Eriani, & Martin, 2020). The enzyme is NTP-dependent and is also known to exhibit 5’-triphosphatase activity. nsp13 is most active in the presence of the RdRp, which suggests that the helicase is required for high-efficiency copying of the viral genome (Adedeji et al., 2012). A recently published 3D electron microscopy (3DEM) structure of the nsp7-nsp82-nsp12/nsp132 heterohexamer provide a structural model for how two copies of the helicase could interoperate with RdRp during RNA synthesis (PDB ID 6XEZ (J. Chen et al., 2020)).

nsp13, a member of helicase superfamily 1, consists of 596 amino acid residues. It adopts a triangular pyramid-like structure consisting of five domains (Zn^++^-binding, stalk, 1B, 1A, and 2A), with each domain directly or indirectly involved in the helicase function. There are three Zn^++^-binding sites located within the N-terminus of the enzyme, involving conserved cysteine and histidine residues (Zn^++^-1: C5, C8, C26, C29; Zn^++^-2: C16, C19, H33, H39; Zn^++^-3: C50, C55, C72, H75). NTPase activity is mediated by six conserved residues situated at the base of the 1A and 2A domains (K288, S289, D374, E375, Q404, R567). The nucleic acid binding channel is formed by domains 1B, 1A, and 2A (Jia et al., 2019). Sequence alignment of SARS-CoV-1 nsp13 with SARS-CoV-2 nsp13 revealed near-perfect identity with a single amino acid difference (I570V). The experimental structure of SARS-CoV-1 nsp13 (PDB ID 6JYT (Jia et al., 2019)) provided the template for Rosetta computation of the SARS-CoV-2 nsp13 homology model used to analyze its evolution in 3D (Fig. nsp13).

Overall substitution trends for nsp13 and energetics analysis results are summarized in Tables 1 and 2. The double substitution P504L;Y541C is the most common nsp13 USV, observed 1,607 times in the GISAID dataset. No substitutions were observed for 11 of the 12 Zn^++^-binding residues. A single substitution was observed for Histidine 33 changing to Glutamine (H33Q), which appears unlikely to abrogate binding of Zn^++^. Potentially important amino acid substitutions involve R337 and R339, two residues known to support helicase activity that are positioned at the entrance of the nucleic acid binding channel. Substitutions were observed in the R337L;A362V and R339L USVs. A SARS-CoV-1 R337A;R339A double substitution showed decreased helicase activity (Jia et al., 2019). It is, therefore, likely that R337L and R339L substitutions in SARS-CoV-2 nsp13 reduced enzyme activity. Another interesting substitution involves the R567, which is important for NTP hydrolysis in SARS-CoV-1 nsp13 (Jia et al., 2019). An R567I substitution occurs in the context of the double substitution USV (V456F;R567I; GISAID dataset count=1) and may reduce SARS-CoV-2 nsp13 helicase activity.

Figure nsp13. (A) Space-filling representation of the computed structural model of nsp13 (green; based on PDB ID 6JYT (Jia et al., 2019)). The RNA helicase active site is located in the upper half of the protein. (B) Space-filling representation of the experimental structure of the nsp132-nsp7/nsp82/nsp12 heterohexamer (PDB ID 6XEZ (J. Chen et al., 2020)), viewed to show the RNA double helix, and (C) viewed looking down the RNA helix axis, showing the two helicase active sites presented to the RNA. (color coding for B and C: nsp13-green, otherwise same color coding as Figure RdRp.)

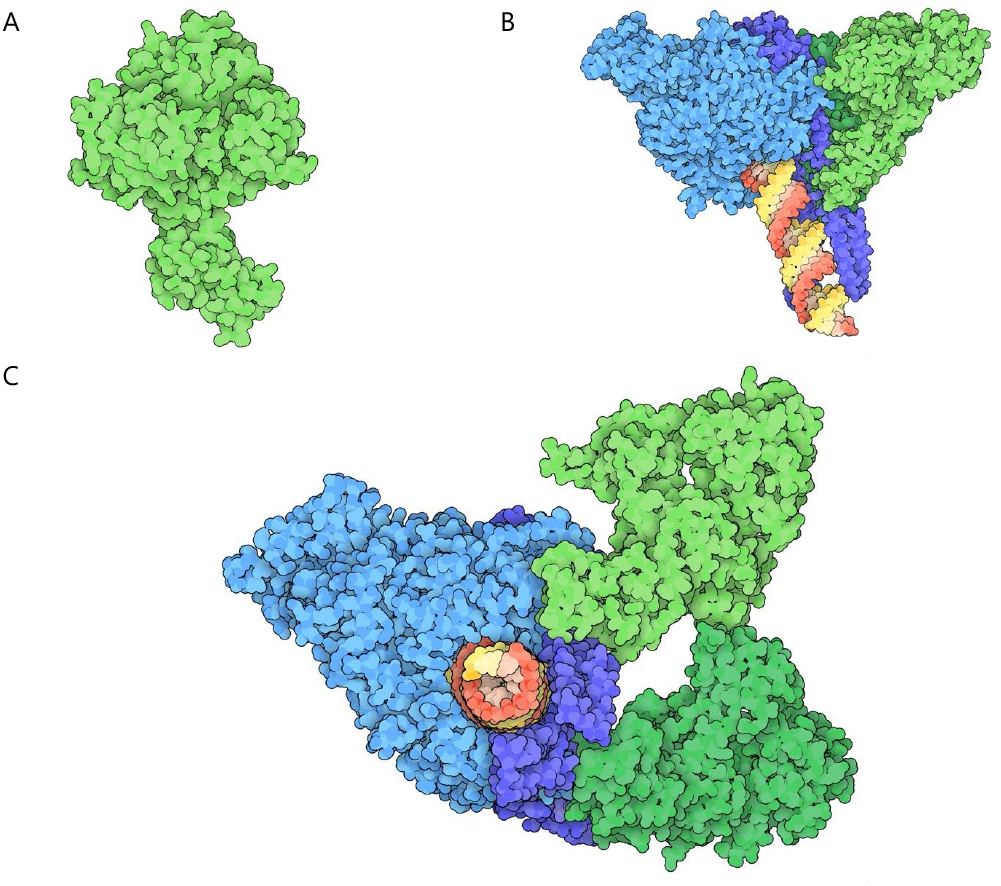

### Non-structural Protein 14 Proofreading Exoribonuclease (nsp14)

nsp14 is a 527-residue protein that acts as both a proofreading exoribonuclease and a methyltransferase to synthesize the N7-methyl-guanine cap 5’ for the mRNA-like genome (Khailany, Safdar, & Ozaslan, 2020; Shannon et al., 2020). It is encoded as part of polyprotein pp1ab and is excised by nsp5. Following excision, it is thought to form a 1:1 complex with non-structural protein 10 (nsp10) to proofread newly formed RNAs synthesized by the RdRp heterotetramer (Eckerle et al., 2010). (N.B.: nsp10 also forms a heterocomplex with nsp16, for which there is an experimental structure available from the PDB (see nsp10/nsp16 section below)). At the time of writing there were no publicly available structures of SARS-CoV-2 nsp14. A computed homology model was used to analyze the evolution of nsp14, based on the structure of SARS-CoV-1 nsp14 (PDB ID 5C8S (Ma et al., 2015)), with which it shares ~95% sequence identity (Fig. nsp14). Superposition of the methyltransferase catalytic centers of SARS-CoV-2 nsp14 and SARS-CoV-1 nsp14 revealed 100% conservation of active site residues, including both the cap binding residues (N306, C309, R310, W385, N386, N422, and F426) and the S-adenosyl methionine (SAM) binding residues (D352, Q354, F367, Y368, and W385). The active site of the exoribonuclease proofreading domain of nsp14 contains a D-E-D-D-H motif (D90, E92, D243, D273, H268), which is identical to the corresponding motif found in SARS-CoV-1 Nps14 (Ma et al., 2015).

Overall substitution trends for nsp14 and energetics analysis results are summarized in Tables 1 and 2. A320V (conservative, core) was the most common substitution, occurring in six USVs with a total GISAID dataset count of 327. A320V also occurred in four double substitution USVs (A320V/D496N, A320V/K349N, A320V/P355S, A320V/A323S). F233L (conservative, core) was the second most common substitution, occurring in 4 USVs, and observed in 273 independently sequenced genomes. It occurred in both a single substitution USV (F233L) and in three double substitution USVs (F233L/A360V, A23S/F233L, F233L/S461P). Two USVs (sequenced in same geographic location) had surprisingly large numbers of amino acid changes and very large ΔΔG^App^ values. The first had five substitutions (T193K/D352E/D358E/Y361K/E364Q), none of which were observed in single substitution USVs. The other had 14 substitutions (Y64F/N67Y/Y69F/P70L/N71Y/M72L/I74F/E77V/I80F/R81S/H82L/V83F/W86C/I87F) with only P70L being observed in another USV as a single substitution. Given the large number of substitutions, extremely unfavorable apparent stabilization energy changes (ΔΔG^App^~20 REU and ~56REU, respectively), and the fact that they were detected only once, we believe that both of these USVs are the result of sequencing artifacts. No substitutions were observed within the active site of the exoribonuclease proofreading domain. The methyltransferase domain displayed a high level of conservation with only three of 12 active site residues substituted. Two guanine cap binding residues (N306 and F426) were found substituted, with N306S (conservative, surface) observed as a single amino acid change and F426L observed once in the double-substitution USV F426L;S448Y. One SAM binding residue was substituted: Q354H (non-conservative, boundary) was observed in five independently sequenced viral genomes.

While we did not generate structural models of the nsp14/nsp10 heterodimer, the structure of SARS-CoV-1 nsp14/nsp10 heterodimer (PDB ID 5C8S (Ma et al., 2015)) allowed us to predict which SARS-CoV-2 amino acid changes may affect nsp10/nsp14 heterodimer formation. Sixteen nsp14 sites of substitution (T5, P24, H26, L27, K47, M62, N67, Y69, V101, N129, T131, K196, V199, I201, P203, and F217, giving a total of 21 distinct substitutions) and eight nsp10 sites of substitution (T12, A18, A20, Y30, A32, I81, K93, and K95, giving a total of 13 distinct substitutions) were mapped to the putative nsp14/nsp10 interface, of which 18 were conservative and 16 were non-conservative. The most prevalent substitutions were T12 (surface, T12I and T12N), A32 (surface, A32S and A32V), H26Y (surface), and P203 (surface, P203L and P203S).

Figure nsp14. (A) Space-filling representation of the computed structural model of the nsp10/nsp14 heterodimer bound to GpppA and S-adenosyl homocysteine (based on PDB ID 5C8S (Ma et al., 2015)). (B) Rotated 90° about the vertical. Color coding: nsp14-light blue; nsp10-dark blue; GpppA-yellow/orange; Exoribonuclease active site Mg^++^ divalent cation: magenta.

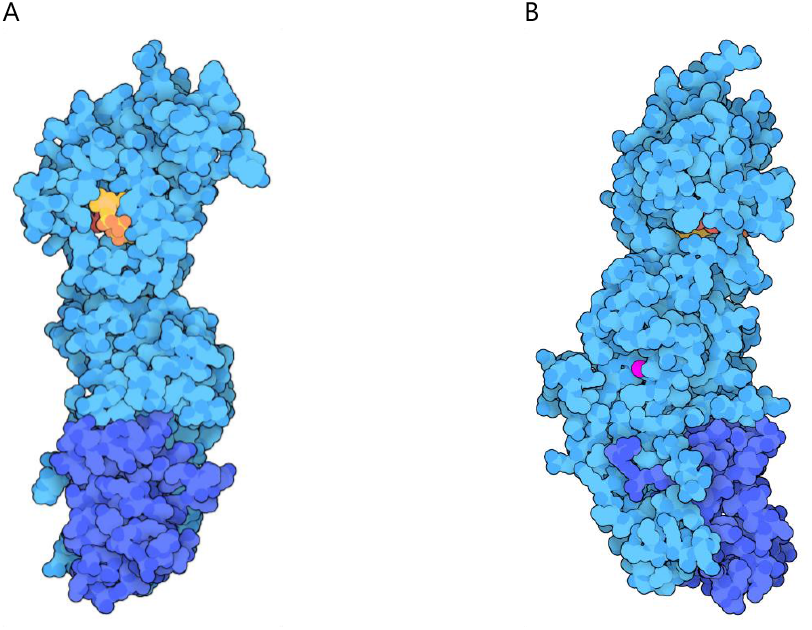

### Non-structural Proteins 10 and 16 Methyltransferase (nsp10/nsp16)

Non-structural proteins nsp10 and nsp16 are both found within pp1ab, from which they are excised by nsp5. Together, nsp10 and nsp16 form a stable heterodimer that functions as a methyltransferase, acting on the 2’ OH of the ribose of the first nucleotide of the viral genome (*i.e*., 5’(m7Gp)(ppAm)[pN]n, where Am denotes 2’-O-ribose methyl-adenosine). This process renders the viral cap structure indistinguishable from that of eukaryotic cap-1, thereby disguising the viral genome so that it resembles cellular RNAs typically found in multicellular organisms and protecting the viral genome from cellular 5’ exonucleases. Enzyme activity of nsp16 depends on SAM as a cofactor, which donates the methyl group from the methionine group for transfer to the ribose of the capped viral RNA (Y. Chen et al., 2011). (N.B.: Capping of the viral RNA is carried out by the N7-guanine methyltransferase domain of nsp14 (Ma et al., 2015)). The structure of the SARS-CoV-2 nsp10/nsp16 heterodimer (PDB ID 6WVN (Minasov et al., 2020)) revealed a heterodimer extremely similar to that of its SARS-CoV-1 homolog (sequence Identities ~93% (for nsp10) and ~98% (for nsp16); r.m.s.d.~1.1Å for PDB ID 6WVN *versus* PDB ID 2XYQ (Decroly et al., 2011)).

The SAM binding site includes residues N43, G71, G73, G81, D99 (3 interactions), D114, C115, D130, and M131 (Rosas-Lemus et al., 2020). The N7-methyl-GpppA binding site consists of residues K24, C25, L27, Y30 (2 interactions), K46, Y132, K137 (2 interactions), K170, T172, E173, H174, S201 (2 interactions), and S202 (4 interactions). Efficient catalytic activity of nsp16 depends on heterodimerization with nsp10, which possesses two zinc-binding motifs (PDB ID 6ZCT (Rogstam et al., 2020)). The two Zn^++^-binding sites of nsp10 are composed of residues C74, C77, H83, and C90; and C117, C120, C128, and C130, respectively.

Polar interactions within the nsp10/nsp16 interface include nsp10:L45-nsp16:Q87; nsp10:G94-nsp16:R86; nsp10:K93-nsp16:S105; nsp10:K43-nsp16:K138; nsp10:Y96-nsp16:A83; and nsp10:A71/G94-nsp16:D106. There is also a salt bridge between H80 and D102 in the SARS-CoV-1 nsp10/nsp16 heterodimer (Y. Chen et al., 2011). At the time of analysis, there was one PDB structure of SARS-CoV-2 nsp10 alone (PDB ID 6ZCT (Rogstam et al., 2020)). A dozen co-crystal structures of the SARS-CoV-2 nsp10/nsp16 heterodimer are available from the PDB, together with nearly 20 structures of nsp10/nsp16 from SARS-CoV-1 and MERS CoV. In the case of SARS-CoV-1, nsp10 also forms a heterodimer with nsp14 (*eg*., PDB ID 5C8S (Ma et al., 2015)). Evolutionary analyses of the nsp10/nsp16 heterodimer that follow were carried out using PDB ID 6WVN (Fig. nsp10/nsp16).

Overall substitution trends for nsp10 and nsp16 and energetics analysis results are summarized in Tables 1 and 2. Several observed substitutions are noteworthy. Two USVs involving SAM binding residues in nsp16 include D99N (non-conservative; core) and D114G (non-conservative; surface), both of which may alter binding affinity to the SAM moiety due to loss of the negative charge upon substitution. Indeed, modeling indicates reduced stability (ΔΔG^App^~7REU in the case of D114G). M131I (conservative; boundary) may also affect SAM binding. By perturbing SAM binding, these substitutions may influence the ability of the enzyme to methylate the first ribose of the viral cap, although these predictions await experimental testing. USVs involving 7-methyl-GpppA binding residues in nsp16 include K24N (non-conservative; surface), D75Y (non-conservative; surface), and S202F (non-conservative; boundary). All of these substitutions had destabilizing effects, with ΔΔG^App^ >7 REU for S202F. D75Y appears to form a new hydrogen bond with the 7-methyl-GpppA, which would slightly shift its position in the binding pocket (Fig. nsp10/nsp16). Only one nsp10 USV affected the Zn^++^-binding residue C130 (C130S;D131H), which would be unlikely to abrogate cation binding.

A number of sites near the protein-protein interface were also substituted, any one of which may affect heterodimer stability, including nsp10 residues K43, T47, T58, F68, and K93; and nsp16 residues P37, G39, M41, V44, T48, G77, V78, P80, R86, T91, D108, T110, M247, and P251. Nine of the interfacial substitutions were conservative and mildly destabilizing, although nsp16 M247I had a more pronounced effect with ΔΔG^App^ > 10 REU. Of the 16 non-conservative interfacial substitutions V78G was most common, appearing in 42 GISAID sequences and three USVs, in two cases occurring concurrently with amino acid changes for P80 (boundary) (P80A and P80L), suggesting that greater flexibility in this region of the protein may be tolerated. Four substitutions were identified that could introduce new hydrogen bonds spanning the heterodimer interface (P37S, G39S, M41T, and G77R), although each of these substitutions appears mildly destabilizing as judged by the results of ΔΔG^App^ calculations with Rosetta.

Figure nsp10/nsp16. Space-filling representation of the experimental structure of the nsp10(dark blue)/nsp16(light blue) heterodimer bound to N7-methyl-GpppA (orange) and SAM (red) (PDB ID 6WVN (Minasov et al., 2020)).

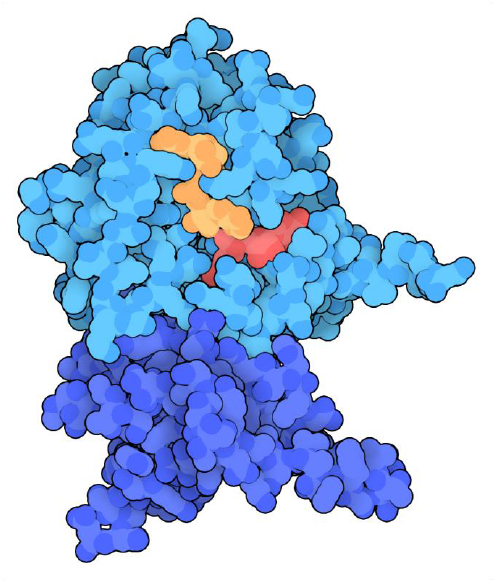

### Structural Spike Surface Glycoprotein (S-protein)

The SARS-CoV-2 spike protein (S-protein) is a membrane-anchored homotrimeric class I fusion protein, that is 1273 residues in length and contains 22 N-linked glycosylation sites (Watanabe, Allen, Wrapp, McLellan, & Crispin, 2020) per monomer (Fig. S-protein A). The S-protein supports viral entry *via* host cell attachment and virion-host membrane fusion. Attachment to a host cell is mediated through the interaction of the S-protein receptor-binding domain (RBD, located in domain S1) with the angiotensin-converting enzyme 2 (ACE2) receptor (Fig. S-protein B). Fusion of the virion to the host cell membrane occurs after cleavage of the S-protein between the S1 and S2 domains, with an additional cleavage (S2’) occurring near the fusion peptide (FP) domain, which is responsible for anchoring to the host cell membrane.

The first experimental structures of the S-protein deposited to the PDB include the pre-fusion state of the S-protein in two conformations–one with all three RBDs in a closed conformation (PDB ID 6VXX (Walls et al., 2020)) and one with RBD protruding upwards (PDB ID 6VSB (Wrapp et al., 2020)). A subsequently deposited PDB structure (PDB ID 6X2B (Henderson et al., 2020)) revealed two upwards protruding RBDs; however, only a single RBD is necessary for ACE2 binding. It is not yet known if protrusion of the RBD from the S-protein trimer is necessary for binding to ACE2 or, as a recent meta-analysis of cryo-EM data suggests (Melero et al., 2020) that interconversion of the RBD between closed and open states represents an intrinsic property of the S-protein. Structures of the S-protein RBD were determined by X-ray crystallography early in the pandemic, both bound to full-length ACE2 receptor (PDB ID 6M17 (Yan et al., 2020)) and bound to relevant ACE2 binding domains (PDB ID 6M0J (Lan et al., 2020); PDB ID 6LZG (Q. Wang, Y. Zhang, et al., 2020)).

Overall substitution trends for the S-protein and energetics analysis results are summarized in Tables 1 and 2. The most commonly observed amino acid change from the reference sequence was D614G, a non-conservative substitution occurring in the SD2 boundary region of the S1 domain (Fig. S-protein C). This substitution appears 21,014 times as a single point substitution and 3,523 times in double or multi-point substitution contexts, accounting for ~68% (805/1190) of all USVs and ~74% (24,537/33,290) of all sequenced genomes downloaded from GISAID. While this substitution is estimated to be slightly destabilizing *versus* the reference sequence (~+0.6 REU), it seems to have emerged early in the pandemic and G614 is now the dominant form of the S-protein worldwide (Korber et al., 2020). The question of if and why G614 is preferred *versus* D614 continues to be debated. It has been hypothesized that this substitution confers increased infectivity, possibly by reducing the pre-emptive shedding of the S1 domain and increasing the total amount of S-protein incorporated into virions (Zhang et al., 2020). A recent cryo-EM-based structural characterization of an engineered D614G S-protein revealed a significantly increased population of conformations in which RBDs are in the open state (PDB ID 6XS6 (Yurkovetskiy et al., 2020)). Interestingly, the measured binding affinity of the G614 spike for ACE2 was slightly lower compared to the D614 variant. The increased population of open conformations in G614 was correlated with loss of inter-protomer contacts in the trimeric spike between D614 from the S1 domain and T859 from the S2 domain this contact was postulated to be a “latch” that favors the closed state (Fig. S-protein C).

Definitive elucidation of the effects of D614G and other substitutions on S-protein stability would require measuring impacts on the stability of all states (pre-fusion, post-fusion, open, closed). Moreover, amino acid changes may impact the structure and stability of complexes with binding partners (ACE2 and other possible co-receptors) and proteases responsible for S-protein cleavage. In this work, we limited our analysis of substitutions to two S-protein PDB structures available in June 2020: a pre-fusion all-closed RBD conformation (PDB ID 6VXX (Walls et al., 2020)), and the RBD-ACE2 complex (PDB ID 6M17 (Yan et al., 2020)). Our methodology could be extended to other structures that continue to be determined at a fast clip, including antibody-bound or inhibitor-bound structures.

### Receptor Binding Domain Substitutions

The most prevalent RBD substitution is the double substitution D614G;T478I (count=57), which is located in an RBD substitution that includes D614G;S477N (count=43), G476S (count=9), and V483A (count=28). These three residues are located in a loop region at the edge of the RBD-ACE2 binding interface. The substitutions were calculated to be destabilizing. Further experimental work will be required to understand how these substitutions might affect the binding of the RBD to ACE2.

### Cleavage-site Substitutions

It was recognized early on in the pandemic that the S-protein possesses a potential furin cleavage site (residue 681-PRRAR/SV-residue687). Furin cleavage is thought to represent another mechanism for transition into a fusion-compatible state (Johnson et al., 2020), thereby contributing to virulence. However, the virus was still found to be infectious upon deletion of the furin cleavage site, indicating that it may not be required for viral entry (Walls et al., 2020) but may affect replication kinetics (Johnson et al., 2020). In that context, it is remarkable that several substitutions are observed within the putative furin cleavage site (P681L/S/H, R682Q/W, R683P/Q, A684T/S/V, S686G). Others have reported that amino acid changes occurred in the furin cleavage site (Xing, Li, Gao, & Dong, 2020). Furin cleavage requires a polybasic motif, but the enzyme is not very stringent, suggesting that these altered sites may still be proteolytically cleaved (Shiryaev et al., 2013).

Prior to virus entry, the S-protein undergoes a second cleavage at the S2’ site (residue 811-KPSKR/SFI-residue 818), which exposes the fusion peptide. This component in the S2 domain fusion machinery attaches to the host cell membrane to initiate membrane fusion. The identity of the enzyme(s) responsible for the cleavage at this site is not known, although given the cleavage site sequence it is thought that it is a furin-like enzyme (Coutard et al., 2020; Hoffmann et al., 2020). We identified several substitutions within the S2’ cleavage domain, including P812L/S/T, S813I/G, F817L, I818S/V. Further experimental study of these substitutions and the replication properties of these altered viruses may provide insight into the role played by furin cleavage in SARS-CoV-2 infection and virulence.

### Fusion Machinery Substitutions

Following cleavage at the S2’ site, the S-protein fuses the viral membrane with the host cell endosomal membrane. S2’ cleavage exposes the fusion peptide (loosely defined as residues 816-855), which then inserts into the host cell membrane. SARS-CoV-1 and SARS-CoV-2 fusion peptide sequences are very similar (~93% sequence homology) (Tang, Bidon, Jaimes, Whittaker, & Daniel, 2020). Our analyses, however, identified many USVs in which amino acid changes in this segment occurred during the pandemic (*i.e*., L821I, L822F, K825R, V826L, T827I, L828P, A829T, D830G/A, A831V/S/T, G832C/S, F833S, I834T). The active conformation and mode of insertion of the SARS-CoV-2 fusion peptide have not been experimentally characterized, making the impact of these substitutions impossible to assess. It may be significant that many of the observed amino acid changes in the fusion peptide are conservative.

A partial structure of the post-fusion state of the S-protein was determined early in the pandemic (PDB ID 6LXT (Xia et al., 2020)). During the final stages of membrane fusion, the HR1 and HR2 domains of class I fusion proteins assemble into a 6-helix bundle (Tang et al., 2020). HR2 sequences of SARS-CoV-1 and SARS-CoV-2 are identical. Differences in HR1 sequences between the two viruses suggest that SARS-CoV-2 HR2 makes stronger interactions with HR1 (Xia et al., 2020). Several substitutions occur on the solvent accessible surface of the HR1 domain (*eg*., D936Y, S943P, S939F) and do not seem to participate in stabilizing interactions with HR2. It is, therefore, unclear how these non-conservative amino acid changes might affect the packing or stability of the post-fusion S-protein. Other residues in HR2 undergoing substitutions during the pandemic (*eg*., K1073N, V1176F) or in the transmembrane or cytoplasmic tail domains (*e.g*., G1219C, P1263L) are not present in the post-fusion structure of the 6-helix bundle. Future experimental work to determine the conformation of the FP, HR1, HR2, and TM domains along the entire membrane fusion pathway should help to elucidate substitutions affecting these segments of the S-protein.

### N-terminal Domain Substitutions

The N-terminal domain (NTD) of the S-protein includes the first ~300 residues. Thus far, the function of the NTD has not been experimentally characterized. It is the target of neutralizing antibodies obtained from convalescent serum of individuals previously infected with SARS-CoV-2 (Chi et al., 2020), and the site of many substitutions identified in this work. Interestingly, the S-protein NTD of MERS-CoV utilizes sugar-binding receptors as a secondary means of interaction with host cells. Awasthi and co-workers have proposed that the SARS-CoV-2 S-protein NTD may do the same. Their computational modelling results suggest that that the NTD β4-β5 (69-HVSGTNGTKRF-79) and β14-β15 (243-ALHRSYLTPGDSSSGWTAGA-262) loop regions form a sialoside-binding pocket that would support engagement of host cell sialic acid moieties (Awasthi et al., 2020). Our analyses documented that virtually all of the residues in these loops underwent amino acid changes during the pandemic (β4-β5: H69Y, V70F, S71F/Y, G72R/E/W, T73I, N74K, G75R/V/D, T76I, K77M/N, R78M/K, F79I; β14-β15: A243S/V, H245Y/R, R246I/S/K, S247R/N/I, Y248S, L249S/F, T250N, P251S/H/L, G252S, D253G/Y, S254F, S255F/P, S256P, G257S/R, W258L, A260S/V, G261V/S/D/R, A262S/T). Unfortunately, these loop regions are largely absent from the 3DEM structures used in our analysis (PDB ID 6VXX (Walls et al., 2020); PDB ID 6VSB (Wrapp et al., 2020)), presumably because they are largely unstructured. Notwithstanding the paucity of 3D structural information, many of these substitutions would likely disrupt stabilizing electrostatic interactions between NTD and sialic acid derivatives postulated by Awasthi and coworkers (Awasthi et al., 2020). Experimental work will be required to evaluate SARS-CoV-2 NTD interactions with sialic acid and how amino acid changes in the NTD affects binding to host cells.

Figure S-protein. (A) Space-filling representation of the experimental structure of the S-protein homotrimer with one RBD protruding upwards (PDB ID 6VSB (Wrapp et al., 2020)); Color coding: RBD up monomer-dark pink, RBD down monomers purple, N-linked carbohydrates-light pink). Membrane spanning portions are depicted in cartoon form. (B) Ribbon/atomic stick figure representation of the RBD interacting with ACE2 (PDB ID 6LZG (Q. Wang, Y. Zhang, et al., 2020)). RBD ribbon color: cyan or purple (substituted residues), atom color coding: C-cyan or purple, N-blue, O-red). ACE2 ribbon color: grey; atom color coding: C-grey, N-blue, O-red. (C) Ribbon/atomic stick figure representation of the D614 reference sequence structure (PDB ID 6VSB (Wrapp et al., 2020); D614 ribbon color: cyan; atom color coding: C-cyan, N-blue, O-red) overlayed on the D614G substitution structure (PDB ID 6XS6 (Yurkovetskiy et al., 2020); D614G ribbon color-grey; atom color coding: C-grey, N-blue, O-red). H-bonds denoted with dotted yellow lines.

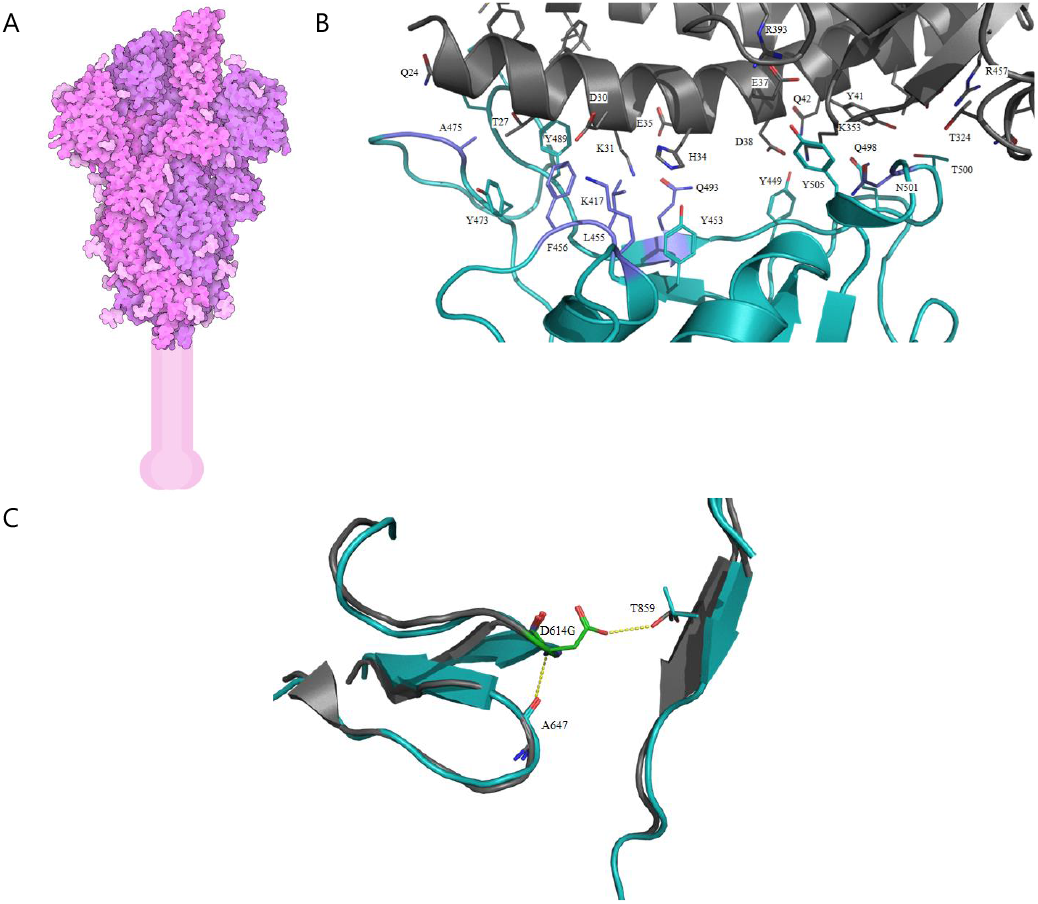

### Structural Nucleocapsid Protein (N-protein)

The nucleocapsid N-protein (422 residues in length) forms a ribonucleoprotein (RNP) complex with viral RNA to protect and stabilize it within the viral envelope. The N-terminal domain (NTD) is responsible for nucleotide binding, while the C-terminal domain (CTD) is responsible for dimerization (Kang et al., 2020). They are connected by a serine/arginine-rich (SR) linker region that is thought to be intrinsically disordered on the basis of amino acid composition. Experimental structures for the N- and C-terminal domains of the SARS-CoV-2 N-protein (PDB ID 6VYO (Chang, 2020); PDB ID 6YUN (Zinzula et al., 2020)) were used for the evolutionary analysis (Fig. N-protein). Residues for which 3D structural information were not available include 1-48, 174-247, and 365-422.

Overall substitution trends for the N-protein and energetics analysis results are summarized in Tables 1 and 2. The most frequently observed USV (R203K/G204R) observed 11,425 times affects two residues within the SR linker region for which there is no 3D structural information. Thereafter, R203K (conservative, atomic coordinates not present in either PDB structure) is the most common substitution, observed 13,130 times, and occurring in 272 other USVs. The R203K/G204R double substitution also appears in a majority of the triple point substitutions (228/237 triples, 35/36 quadruples, 1/2 quintuples). Another interesting USV includes the 5-point substitution, R36Q/R203K/G204R/T135I/K373N). The NTD contains several basic residues (Arginine and Lysine) that are located in the finger subdomain and appear likely to interact with the RNA. Several substitutions in these finger-domain residues were observed in various USVs (*eg*., R92S, R93L, R88L). If and how these may affect RNA-binding remains to be investigated.

Figure N-protein. Space-filling representation of the experimental structures of N-protein domains (PDB IDs 6VYO (Chang, 2020) and 6YUN (Zinzula et al., 2020)). [N.B. The relative orientations of the N-terminal (upper: residues 49-173) and C-terminal (lower: residues 248-364) domains was chosen arbitrarily. No structural information is currently available for residues 1-48, 174-247, and 365-422.]

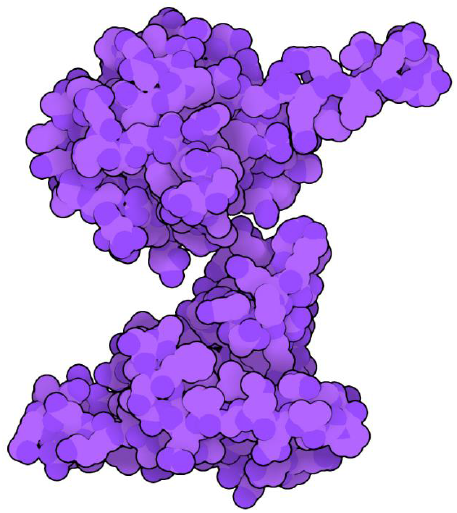

### Structural Protein Ion Channel Envelope Protein (E-protein)

The integral membrane E-protein is the smallest of the SARS-CoV-2 structural proteins (75 residues). It plays important roles in virus-like particle production and maturation. Coronavirus E-proteins are co-translationally inserted into the endoplasmic reticulum (ER) and transported to Golgi complexes (Ruch & Machamer, 2012). Although it is abundantly expressed within the cell, only a modest number of copies are actually incorporated into the viral envelope (estimated number/virion~20 for SARS-CoV-1, (DeDiego et al., 2007)). Instead, most of the protein participates in virion assembly and budding together with the SARS-CoV-2 integral membrane M-protein (also a virion structural protein). Additional functions of the E-protein are thought to include, preventing M-protein aggregation and inducing membrane curvature (Schoeman & Fielding, 2019). Recombinant coronaviruses lacking E-proteins display weakened maturation, reduced viral titers, or yield incompetent progeny, highlighting its role in maintaining virion integrity.

The E-protein consists of a shorter hydrophilic N-terminal segment, a longer hydrophobic transmembrane domain (TMD), and a hydrophilic C-terminal domain. An amphipathic α-helix within the TMD oligomerizes into an homopentameric arrangement perpendicular to the plane of the lipid bilayer forming an ion-conducting viroporin (Schoeman & Fielding, 2019). Residues lining the pore include N15, L19, A22, F26, T30, I33, and L37. The NMR structure of the SARS-CoV-1 E-protein (PDB ID 5X29 (Surya, Li, & Torres, 2018)) served as the template for generating the computed structural model of the SARS-CoV-2 E-protein that was used for analyzing its evolution in 3D (Fig. E-protein). The N-terminal seven residues and the C-terminal ten residues were omitted from the homology model, because they were not reported in the SARS-CoV-1 NMR structure.

Overall substitution trends for the E-protein and energetics analysis results are summarized in Tables 1 and 2. S68F (non-conservative, structural location unknown) is the most common USV, observed 107 times in the GISAID dataset. The most intriguing changes in the protein are L37R and L37H USVs, located near the entrance to the pore (Fig. E-protein). The changes of Leucine to Arginine or Histidine are notable because the canonical transmembrane domain lacks charged residues. The SARS-CoV-1 E-protein is preferentially selective for cations, although it can transport anions (Verdia-Baguena et al., 2012). Substitution of L37 to a positively charged residue may affect ion passage selectivity and/or its ability to transport ions.

SARS-CoV-1 E-protein is N-linked glycosylated at N66 (S. C. Chen, Lo, Ma, & Li, 2009). At the time of writing, there were no published reports pertaining to SARS-CoV-2 E-protein glycosylation. The corresponding residue in SARS-CoV-2 E-protein is N66, which underwent substitution to Histidine in a single USV (N66H) that would abrogate glycosylation. Observed amino acid substitutions involving loss or gain of other potential sites of N-linked and O-linked glycosylation include A41S, C43S, N48S, S50G, P54S, S55F, S68C, S68F, and S68Y.

Figure E-Protein. (A) Space-filling representation of the computed structural model of the E-protein with individual protomers shown with shades of pink and purple. (B) Ribbon representation with each protomer shown using a different color viewed parallel to the membrane (left, membrane shown, N- and C-termini labeled) and down the five-fold axis from the virion surface (right). (C) Pore-lining substitutions L37R and L37H compared to L37 in the reference sequence (residue 37 is shown in a color-coded space-filling representation; C-gray; O-red; N-blue).

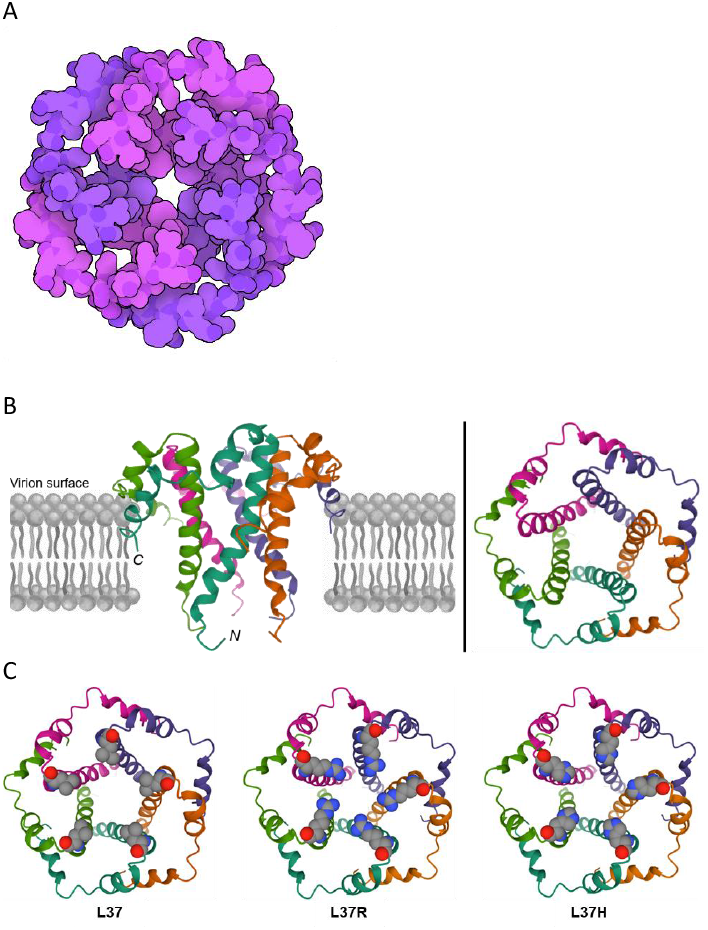

### Structural Integral Membrane Protein (M-protein)

The integral membrane M-protein (222 residues in length) is the most abundant structural protein in the SARS-CoV-1 virion (Siu et al., 2008). It is co-translationally inserted into the ER and transported to Golgi complexes (S. C. Chen et al., 2009), where it is responsible for directing virus assembly and budding *via* interactions with E-, N-, and S-proteins. The SARS-CoV-2 M-protein is predicted to consist of a small glycosylated amino-terminal ectodomain, a triple-membrane spanning domain, and a carboxyl-terminal endodomain that extends 6-8 nm into the viral particle. The C-terminal portion of coronaviral M-proteins bind to the N-protein within the cell membrane of the ER or Golgi complex, stabilizing the nucleocapsid and the core of the virion. M-proteins also interacts with the E-protein to trigger budding, and with the S-protein for incorporation into virions (Tseng, Chang, Wang, Huang, & Wang, 2013). Following assembly, virions are transported to the cell surface and released *via* exocytosis. The M-protein is believed to exist as a dimer in the cell membrane and may adopt two conformations that allow it to bend the membrane and interact with N-protein/RNA RNP (Neuman et al., 2011). Sequence alignment of SARS-CoV-2 M-protein to its SARS-CoV-1 homolog revealed high sequence identity (~90%). The M-protein structural model used for analyzing evolution in 3D was computed by the David Baker Laboratory during a CASP competition (CASP-C1906 Stage 2, Fig. M-Protein).

Overall substitution trends for the M-protein and energetics analysis results are summarized in Tables 1 and 2. T175M (non-conservative, surface) is the most common USV, observed 746 times in the GISAID data set (~39% of the observed variant M-proteins). An N5S substitution affects the sole N-linked glycosylation site in the small ectodomain. Given that M-protein glycosylation is not essential for maintaining virion morphology or growth kinetics (Voss et al., 2009), it is unclear if M-protein function is affected by the N5S substitution.

Figure M-Protein. (A) Space-filling representation of the computed structural model of the M-protein protomer. The glycosylated N-terminus is located at the apex of the structure. (B) Ribbon/atomic stick figure representation (Color coding: ectodomain-blue, transmembrane α-helices-red, endodomain-green). N- and C-termini are labeled, together with residues N5, L124, T175, and R186 (shown in ball and stick representation; atom color coding: C-green, O-red, N-blue).

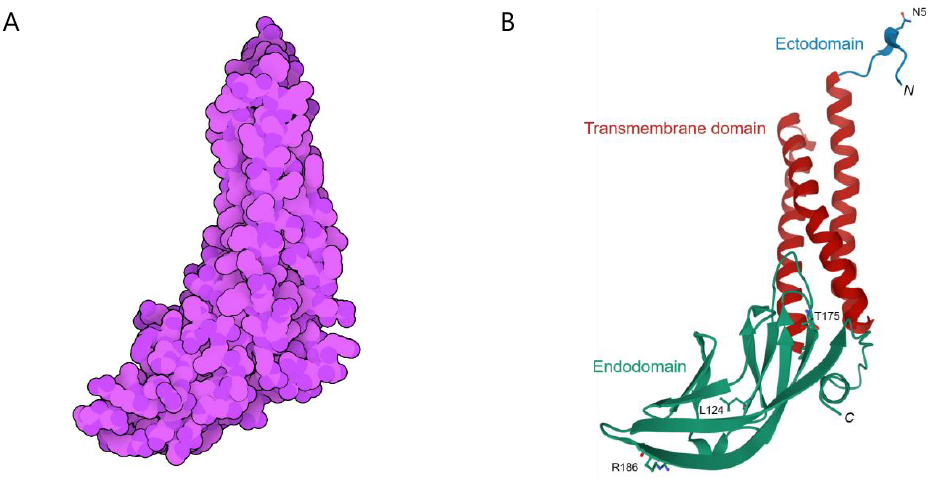

### Implications for the Ongoing Pandemic and Discovery and Development of Effective Countermeasures

Our analyses of SARS-CoV-2 genome sequences archived by GISAID documented that every one of the 29 study proteins underwent amino acid changes *versus* the original reference sequence during the first six months of the pandemic. Most of these substitutions occurred infrequently. Approximately two thirds of the substitutions were non-conservative, and most appear to have arisen from single or double nucleotide changes in the RNA genome. Computational 3D structure modeling of the USVs demonstrated that substitutions primarily occurred in the boundary layers and the surfaces of the viral proteins. The vast majority of the amino acid changes appear to be moderately destabilizing, as judged by the results of energetics (ΔΔG^App^) calculations. Given that most of the viral genomes archived by GISAID were obtained from samples provided by infected individuals, we believe that the viruses and hence the viral proteins were functional and capable of causing disease in humans. Where multiple substitutions were detected in a USV, we believe that most were the product of cumulative changes. At least one of the observed amino acid changes in multi-substitution USVs was almost always detected as a single substitution in another USV derived from a sample collected earlier in the pandemic. There is every reason to believe that the pool of viruses circulating in humans and some mammals (*eg*., *Mustela lutreola* or European mink) around the world today will continue to diverge from the reference sequence. We have made 3D structure models of 7,462 USVs and our analysis results freely available under the most permissive Creative Commons license to facilitate the work of research groups using experimental and computational tools to characterize SARS-CoV-2 protein function and study the structural and functional consequences of the myriad substitutions observed during the first half of 2020.

Some, certainly not all, of the 29 viral proteins analyzed herein represent promising targets for discovery and development of small-molecule anti-viral agents. At the time of writing, one small-molecule drug (remdesivir targeting the RdRp) has received full approval from the US FDA. This compound was originally discovered during the search for an Ebola virus therapeutic. Although it failed to demonstrate efficacy in clinical trials for Ebola victims, the safety profile encouraged the sponsor company (Gilead Sciences Inc.) to successfully repurposed the drug for treatment of SARS-CoV-2 infected individuals. Open access to PDB structures of remdesivir bound to the RdRp sets the stage for structure-guided discovery of second generation nucleoside analogs with superior potency and/or selectivity, more desirable drug-like properties, or better Absorption-Distribution-Metabolism-Excretion profiles (*e.g*., improved oral bioavailability to avoid intravenous administration) (Westbrook, Soskind, Hudson, & Burley, 2020). Open access to our computed 3D structural models of 840 RdRp USVs will provide useful information that may enable drug hunting teams to anticipate potential sources of drug resistance during selection for candidates slated for *in vitro* pre-clinical development studies.

Open access to PDB structures of other essential SARS-CoV-2 enzymes (and their closely related SARS-CoV-1 homologs) have already facilitated initiation of structure-guided drug discovery campaigns for PLPro, nsp5, nsp13, nsp14, and nsp10/nsp16. As for the RdRp, free availability of computed 3D structural models of nearly 1,500 USVs may provide useful information pertaining to potential causes of drug resistance. Knowledge of sequence (and 3D structure) variation during the pandemic could also be used to prioritize these potential drug targets using quantitative assessments of active site conservation. The best drug discovery targets could be those proteins observed to undergo the fewest amino acid changes in their active (or drug-binding) site during the first six months of the pandemic. It is also possible that inhibitors making contacts with residues that are not engaged by substrates will be more susceptible to the emergence of drug resistance.

The S-protein is the target of both monoclonal antibodies (for passive immunization) and vaccines. At the time of writing, a number of monoclonal antibodies had already received Emergency Use Authorization (EUA) from the US FDA (*eg*., bamlanivimab; sponsor company Eli Lilly and Co.). EUAs for one or more vaccines can be reasonably expected before the end of 2020. Open access to a host of PDB structures of the S-protein in various conformational states and in complexes with host cell proteins and Fab fragments of monoclonal antibodies will facilitate the work of research teams focused on discovery and development of second generation monoclonal antibodies and vaccines. Free availability of 689 3D structural models of S-protein USVs may provide insights into potential efficacy failures due to amino acid changes in the S-protein that interfere with viral antigen recognition by antibodies (monoclonal or humoral) or T-cells while preserving ACE2 receptor binding.

### Postscript

Unwillingness in virtually every country to prepare adequately for the possibility that a third coronavirus would jump the species barrier has dealt humanity a devastating blow. Global fatalities attributed to the SARS-CoV-1 outbreak in the early 2000s numbered only 774 (total number of cases=8,098), and there were no deaths in the US. Had the virus rendered asymptomatic individuals infectious, however, we now know that the situation would have been catastrophically worse. The world was fortunate once again, relatively speaking, when MERS-CoV struck nearly a decade after SARS-CoV-1. From 2012 through May 31, 2019, MERS-CoV infected 2,442 individuals and killed 842 worldwide. The virus is currently circulating in dromedary camels in Africa, the Middle East, and southern Asia, with no end to human infections in sight and no effective anti-viral therapies or vaccines available.

Analyzing coronavirus protein evolution in 3D provides a sobering lesson regarding the potential value of proper pandemic preparedness. nsp5 amino acid sequences are highly conserved across all known coronaviruses. For example, SARS-CoV-2 nsp5 is 95% identical in amino acid sequence to that of SARS-CoV-1. nsp5 3D structures are also highly conserved across all known coronaviruses. Indeed, the nsp5 proteins of SARS-CoV-2 and SARS-CoV-1 are extremely similar in 3D structure (α-carbon r.m.s.d.~0.8Å; Fig. SARS-CoV-1/SARS-COV-2). The residues lining the active site are identical and the active sites are structurally similar (non-hydrogen atom r.m.s.d.<0.5Å), precisely because they recognize virtually identical (if not identical) peptide cleavage substrates during viral polyprotein processing. We had every opportunity in the wake of the SARS-CoV-1 medical emergency to discover and develop a drug targeting SARS-CoV-1 nsp5 and other coronavirus main proteases. Structure-guided approaches using PDB ID 1Q2W (Pollack, 2003) and the many structures of SARS-CoV-1 nsp5 subsequently released by the PDB would almost certainly have yielded one (possibly more) potent and selective enzyme inhibitor(s) with good drug-like properties and an acceptable safety profile.

While it may have appeared economically rational to consign SARS-CoV-1 (and the possibility of serious coronavirus outbreaks in humans) to the annals of history, hindsight tells us otherwise. Economists would describe this a “failure of the free market” for emergency medicines. With no clear prospect of income from drugs targeting future epidemics that may never come and no additional incentives provided by public health authorities, biopharmaceutical companies focused on business as usual. The emergence of MERS-CoV and its ongoing persistence in endemic areas should have served as additional warnings that emerging coronavirus pandemics represent a very real and present danger to humanity. A safe and effective drug targeting SARS-CoV-1 nsp5 would almost certainly be working today for SARS-CoV-2! Simply put, an investment of a few hundred million US$ in discovering and developing SARS-CoV-1 nsp5 inhibitors, and eventually drugs in the 2000s, could today have already saved more than a million lives and prevented tens of trillions of US$ in global economic losses (Burley, 2020).

Rising numbers of COVID-19 infections and deaths worldwide as we approach the northern hemisphere winter of 2020-2021 show that we must prepare for the next coronavirus outbreak. The science and the unmet medical needs are clear. It is time for governments, industry, and NGOs to confront the failure of the free market head-on. We scientists have it within our power to present policy makers with viable options that could make all the difference when (it’s no longer a matter of if) another coronavirus jumps the species barrier to human.

Figure SARS-CoV-1/SARS-CoV-2. Ribbon representation overlay of experimental structures of SARS-CoV-2 nsp5 (PDB ID 6LU7; (Jin et al., 2020); Color coding: magenta, purple, gold) and SARS-CoV-1 nsp5 (PDB ID 1Q2W (Pollack, 2003); Color coding: green). Substrate analog inhibitor present in PDB ID 6LU7 is shown as an atomic stick figure (Atom color coding: carbon-white, oxygen-red, nitrogen-blue).

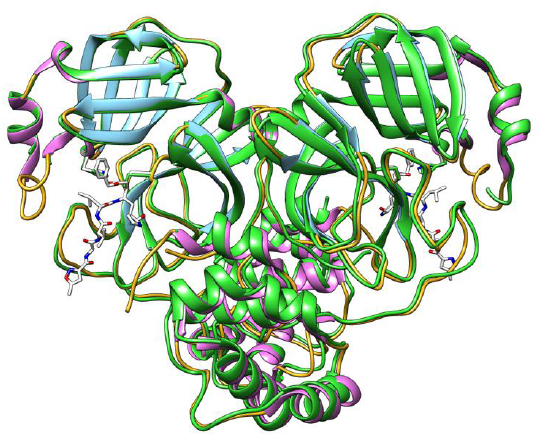

## Materials and Methods

### SARS-CoV-2 Study Protein Sequences

Pre-aligned protein sequences were downloaded in FASTA format from the GISAID website (gisaid.org) (Elbe & Buckland-Merrett, 2017; Shu & McCauley, 2017) on June 25^th^ 2020. Sequence alignments for each of the SARS-CoV-2 proteins (hereafter study proteins) were constructed by removing non-human sequences from the alignment; removing truncated sequences; removing incompletely determined sequences (*i.e*, those with one or more “X” in lieu of an amino acid one-letter code); and eliminating duplicates. Study protein sequences made public by researchers in the People’s Republic of China on January 10^th^ 2020 (GenBank accession code MN908947.3) (Wu et al., 2020) were defined as the “reference sequence” for each individual study protein and all unique sequence variant (USV) or amino acid substituted forms of individual study proteins were compared with their respective reference sequence. We have assumed that none of observed USVs yielded study proteins that either failed to fold or lost necessary biochemical functionality for other reasons, because it is likely given the timing of specimen collection that all of the viral RNAs were isolated from infected individuals and are, therefore, presumed to have been infectious. For sequence identity calculations, GenBank accession code AY278741.1 was used as the source of SARS-CoV-1 protein reference sequences.

### Experimentally-determined Structures of Study Proteins from the PDB Archive

Atomic coordinates for the experimental structures of 13 study proteins were downloaded from the PDB archive *via* the RCSB PDB website (RCSB.org), including nsp3b (part of nsp3) (PDB ID 6WEY (Frick, Virdi, Vuksanovic, Dahal, & Silvaggi, 2020)), Papain-like Proteinase (part of nsp3) (PLPro; PDB ID 6WUU (Rut et al., 2020)), nsp5 (PDB ID 6LU7 (Jin et al., 2020)), nsp7 (PDB IDs 7BV1 and 7BV2 (Yin et al., 2020), nsp8 (PDB IDs 7BV1 and 7BV2 (Yin et al., 2020)), nsp9 (PDB ID 6WXD (Littler, Gully, Colson, & Rossjohn, 2020)), nsp10 (PDB ID 6WVN (Minasov et al., 2020)), nsp12 (PDB IDs 7BV1 and 7BV2 (Yin et al., 2020)), nsp13 (PDB ID 6ZSL, (Newman, 2020)), nsp15 (PDB ID 6VWW (Y. Kim et al., 2020)), nsp16 (PDB ID 6WVN (Minasov et al., 2020)), S-protein (PDB ID 6VXX (Walls et al., 2020); PDB ID 6M17 (Yan et al., 2020)), Orf3a (PDB ID 6XDC (Kern et al., 2020)), and the N-protein (PDB ID 6VYO (Chang, 2020); PDB ID 6YUN (Zinzula et al., 2020)).

### Computed Structural Models of Study Proteins

In three cases, computed structural models for study proteins were downloaded from the Robetta-based predictions from the website for Seattle Structural Genomics Center for Infectious Disease (https://www.ssgcid.org/cttdb/molecularmodel_list/?organism__icontains=COVID-19), including nsp1 (https://robetta.bakerlab.org/domain.php?id=15554), nsp14 (https://robetta.bakerlab.org/results.php?id=15671), and Orf7a (https://robetta.bakerlab.org/results.php?id=15657).

The computed structural model for the SARS-CoV-2 E-protein were generated using the solution state NMR structure of the SARS-CoV-1 E-protein embedded in lyso-myristoyl phosphatidylglycerol micelles (PDB ID 5X29, model 1 (Surya et al., 2018)) as a template, and substituting differing residues using the MUTATE feature of VMD (Humphrey, Dalke, & Schulten, 1996). The structural model was then subjected to 10,000 steps of energy minimization in vacuum using NAMD 2.13 (Phillips et al., 2020) and the CHARMM 36 force field (MacKerell Jr. et al., 1998).

Swiss-Model (Waterhouse et al., 2018) was the source of computed structural models of three study proteins, including nsp3a using 77% sequence identical template SARS-CoV-1 nsp3a (part of nsp3) (PDB ID 2GRI (Serrano et al., 2007)); nsp3c (part of nsp3) using 75% sequence identical template SARS-CoV-1 nsp3c (part of nsp3) (PDB ID 2W2G (Tan et al., 2009)); and nsp3e (part of nsp3) using ~81% identical template SARS-CoV-1 nsp3e (part of nsp3) (PDB ID 2K87 (Serrano et al., 2009)).

Computed structural models for the eight remaining study proteins were obtained from the Rosetta-based Baker group predictions (TS131) CASP website (https://predictioncenter.org/caspcommons/targetlist.cgi; Model 1 was chosen), including nsp2, nsp3a (part of nsp3), UNK (part of nsp3), nsp4, nsp6, the M-protein, Orf6, Orf7b, and Orf8.

nsp11, Orf9b, Orf14, and hypothetical protein Orf10 were excluded from consideration owing to lack of sequence and/or 3D structure data.

### Molecular Visualization and Graphics

The RCSB Protein Data Bank web-native molecular graphics tool (Mol*; (Sehnal, Rose, Koca, Burley, & Velankar, 2018)) was used for visual inspection and comparison of reference and amino-acid-substitute study proteins. Space-filling representation figures were generated using Illustrate (Goodsell, Autin, & Olson, 2019). Ribbon/atomic stick figure representation figures were generated using Mol* and PyMOL (DeLano, 2002).

### Rosetta-based Analyses of Substitution Location(s), Conservation, and Energetics

PyRosetta (Chaudhury, Lyskov, & Gray, 2010) was used to analyze each study protein and its observed USVs. All residue pairs with C_α_-C_α_ distance <5.5Å were considered neighbors, and residue pairs with C_α_-C_α_ distance <11Å were also considered neighbors if their C_α_-Cβ vectors were at an angle <75°. Residue layer identifications were performed on reference (rather than substituted) study-protein structures, based on side chain neighbors within a cone centered on the C_α_-C_β_ vector, which is independent of side chain conformation. The Layer Determination Factor (LDF) is defined as LDF=((cos(θ)+0.5)/1.5)^2^/(1 + exp(d − 9)), where θ is the angle between the C_α_-C_β_ vector of a given residue and that of a neighbor, and d is the C_α_-C_α_ distance between residue and neighbor. LDF is summed over nearby neighbors and if its value is <2, the residue is considered surface. If it is >5.2, the residue is considered core. Otherwise, it is considered boundary.

Amino acid substitution conservation was determined by whether or not a residue change stayed within a residue type group as follows: hydrophobic (A, F, I, L, M, V, W, Y), negatively-charged (D, E), positively-charged (H, K, R), and uncharged hydrophilic (N, Q, S, T) and any substitution to a residue outside the native residue’s group was considered non-conservative. Changes to or from Glycine, Proline, or Cysteine were considered non-conservative. Amino acid substitutions in study proteins were identified by alignment with the reference sequence.

Experimental structures and computed structural models of study proteins were prepared for computational analyses using the Rosetta FastRelax protocol, employing atom positional restraints to limit significant changes to backbone geometry. Homooligomeric proteins were modeled using the symmetric protein modeling framework in Rosetta (DiMaio, Leaver-Fay, Bradley, Baker, & Andre, 2011). Integral membrane proteins were modeled using Rosetta membrane protein modeling framework (Alford et al., 2015).

Structural models for study protein USVs were computed by replacing the reference side chain atomic coordinates in the starting model with those of the substituted amino acid(s) and performing three rounds of Monte Carlo optimization of rotamers for all side chains falling within an 8Å radius of the substitution(s), followed by gradient-based energy minimization of the entire structure, with atom positional restraints to limit significant changes to backbone geometry. Computed structural model optimizations were performed with three different combinations of scoring functions based on previous work (Kellogg, Leaver-Fay, & Baker, 2011), including “hard-hard”, indicating that both side chain optimization and structure minimization were performed with default van der Waals repulsion term in the Rosetta scorefunction, “soft-soft” indicating that for both steps, a different scorefunction was used that has dampened van der Waals repulsion (in this case, the backbone was entirely prevented from moving during minimization), and “soft-hard” indicating that the soft-repulsive score function was used for side chain rotamer optimization, while the hard-repulsive scorefunction was used for energy minimization. The scorefunctions used were REF2015 (Alford et al., 2017) and REF2015_soft (Kellogg et al., 2011) for soluble proteins, and franklin2019 (Alford, Fleming, Fleming, & Gray, 2020) for integral membrane proteins (with a dampened van der Waals repulsion weight in the case of soft repulsion).

Energetic consequences of amino acid substitutions were determined by performing identical side chain optimization and energy minimization on both wild-type and substituted models thrice and subtracting the total energy of the lowest-scoring wild-type model from that of the lowest-scoring substituted model (dividing by the number of symmetric chains where applicable). The “soft-hard” protocol emerged as the preferred method because it generated the lowest number of outliers. Only USVs in which a unique set of substitutions occurred at residue positions that were present in the available study-protein structures were included in the energy analyses (7,462 USVs).

## Supporting information

Supplementary descriptions, figures, tables, and models

## Acknowledgements

We thank all of the many structural biologists who have deposited coronavirus protein structures to the PDB archive since 2002. We also thank Drs. David Baker and Ivan Anischenko for providing computed structural models generated by the David Baker Laboratory, Ms. Virginia Jiang and Dr. Scott Banta for help with Rosetta calculations involving integral membrane proteins, and Dr. Andrew Brooks for advice regarding sequencing artifacts. We gratefully acknowledge contributions from all members of the Research Collaboratory for Structural Bioinformatics PDB and our Worldwide Protein Data Bank partners. RCSB PDB is jointly funded by the National Science Foundation (DBI-1832184), the US Department of Energy (DE-SC0019749), and the National Cancer Institute, National Institute of Allergy and Infectious Diseases, and National Institute of General Medical Sciences (NIGMS) of the National Institutes of Health (NIH) under grant R01 GM133198. The Khare laboratory has been funded by NIH NIGMS (R01 GM132565) and NSF (CBET1923691). J.H.L. was funded by NIH NIGMS T32 Training Grants GM008339 and GM135141. E.A. acknowledges support by a National Institutes of Health MERIT Award (R0137 AI027690) and a Rutgers Center for COVID-19 Response and Pandemic Preparedness Award. J.S. and G.B. are funded by the Busch Biomedical Foundation. The Baum laboratory is funded by NIH R35 GM 136431. The BASIL Consortium is funded by NSF IUSE grants 1709170, 1709355, 1709805, and 1709278. We gratefully acknowledge support for L.H.A.A., A.K., E.M., S.S., B.T., A.T., L.W., and M.O.-A. by the Rutgers University RISE (Research Intensive Summer Experience) Program, an NSF REU for A.T., S.S., L.W., and a New Jersey Space Grant Consortium Award for L.H.A.A. and B.T.

## Author Contributions

S.D.K. and S.K.B. developed the research plan. All authors helped to assemble the data, develop and execute analysis strategies, prepare figures and tables, and write the manuscript.

## Author Information

The authors declare no competing financial interests.

## References

Adedeji, A. O., Marchand, B., Te Velthuis, A. J., Snijder, E. J., Weiss, S., Eoff, R. L., … Sarafianos, S. G. (2012). Mechanism of nucleic acid unwinding by SARS-CoV helicase. PLoS ONE, 7(5), e36521. doi:10.1371/journal.pone.0036521

Alford, R. F., Fleming, P. J., Fleming, K. G., & Gray, J. J. (2020). Protein Structure Prediction and Design in a Biologically Realistic Implicit Membrane. Biophys J, 118(8), 2042–2055. doi:10.1016/j.bpj.2020.03.006

Alford, R. F., Koehler Leman, J., Weitzner, B. D., Duran, A. M., Tilley, D. C., Elazar, A., & Gray, J. J. (2015). An Integrated Framework Advancing Membrane Protein Modeling and Design. PLoS Comput Biol, 11(9), e1004398. doi:10.1371/journal.pcbi.1004398

Alford, R. F., Leaver-Fay, A., Jeliazkov, J. R., O’Meara, M. J., DiMaio, F. P., Park, H., … Gray, J. J. (2017). The Rosetta All-Atom Energy Function for Macromolecular Modeling and Design. J Chem Theory Comput, 13(6), 3031–3048. doi:10.1021/acs.jctc.7b00125

Awasthi, M., Gulati, S., Sarkar, D. P., Tiwari, S., Kateriya, S., Ranjan, P., & Verma, S. K. (2020). The Sialoside-Binding Pocket of SARS-CoV-2 Spike Glycoprotein Structurally Resembles MERS-CoV. Viruses, 12(9). doi:10.3390/v12090909

Berman, H. M., Westbrook, J., Feng, Z., Gilliland, G., Bhat, T. N., Weissig, H., … Bourne, P. E. (2000). The Protein Data Bank. Nucleic Acids Res, 28(1), 235–242. doi:10.1093/nar/28.1.235

Bloom, J. D., Silberg, J. J., Wilke, C. O., Drummond, D. A., Adami, C., & Arnold, F. H. (2005). Thermodynamic prediction of protein neutrality. Proc Natl Acad Sci U S A, 102(3), 606–611. doi:10.1073/pnas.0406744102

Burley, S. K. (2020). How to help the free market fight coronavirus. Nature, 580(7802), 167. doi:10.1038/d41586-020-00888-7

Burley, S. K., Berman, H. M., Christie, C., Duarte, J. M., Feng, Z., Westbrook, J., … Zardecki, C. (2018). RCSB Protein Data Bank: Sustaining a living digital data resource that enables breakthroughs in scientific research and biomedical education. Protein Sci, 27(1), 316–330. doi:10.1002/pro.3331

Burley, S. K., Bhikadiya, C., Bi, C., Bittrich, S., Chen, L., Crichlow, G., … Zhuravleva, M. (2020). RCSB Protein Data Bank: Powerful new tools for exploring 3D structures of biological macromolecules for basic and applied research and education in fundamental biology, biomedicine, biotechnology, bioengineering, and energy sciences. Nucleic Acid Res., gkaa1038. doi:10.1093/nar/gkaa1038

Burley, S. K., Bromberg, Y., Craig, P., Duffy, S., Dutta, S., Hall, B. L., … Zardecki, C. (2020). Virtual Boot Camp: COVID-19 evolution and structural biology. Biochem Mol Biol Educ, 48, 511–513. doi:10.1002/bmb.21428

Chang, C., Michalska, K., Jedrzejczak, R., Maltseva, N., Endres, M., Godzik, A., Kim, Y., Joachimiak, A. (2020). Crystal structure of RNA binding domain of nucleocapsid phosphoprotein from SARS coronavirus 2. doi: 10.2210/pdb6VYO/pdb.

Chaudhury, S., Lyskov, S., & Gray, J. J. (2010). PyRosetta: a script-based interface for implementing molecular modeling algorithms using Rosetta. Bioinformatics, 26(5), 689–691. doi:10.1093/bioinformatics/btq007

Chen, J., Malone, B., Llewellyn, E., Grasso, M., Shelton, P. M. M., Olinares, P. D. B., … Campbell, E. A. (2020). Structural Basis for Helicase-Polymerase Coupling in the SARS-CoV-2 Replication-Transcription Complex. Cell, 182(6), 1560–1573 e1513. doi:10.1016/j.cell.2020.07.033

Chen, S. C., Lo, S. Y., Ma, H. C., & Li, H. C. (2009). Expression and membrane integration of SARS-CoV E protein and its interaction with M protein. Virus Genes, 38(3), 365–371. doi:10.1007/s11262-009-0341-6

Chen, Y., Liu, Q., & Guo, D. (2020). Emerging coronaviruses: Genome structure, replication, and pathogenesis. J Med Virol, 92(4), 418–423. doi:10.1002/jmv.25681

Chen, Y., Su, C., Ke, M., Jin, X., Xu, L., Zhang, Z., … Guo, D. (2011). Biochemical and structural insights into the mechanisms of SARS coronavirus RNA ribose 2’-O-methylation by nsp16/nsp10 protein complex. PLoS Pathog, 7(10), e1002294. doi:10.1371/journal.ppat.1002294

Chi, X., Yan, R., Zhang, J., Zhang, G., Zhang, Y., Hao, M., … Chen, W. (2020). A neutralizing human antibody binds to the N-terminal domain of the Spike protein of SARS-CoV-2. Science, 369(6504), 650–655. doi:10.1126/science.abc6952

Chodera, J., Lee, A. A., London, N., & von Delft, F. (2020). Crowdsourcing drug discovery for pandemics. Nat Chem, 12(7), 581. doi:10.1038/s41557-020-0496-2

Clementz, M. A., Chen, Z., Banach, B. S., Wang, Y., Sun, L., Ratia, K., … Baker, S. C. (2010). Deubiquitinating and interferon antagonism activities of coronavirus papain-like proteases. J Virol, 84(9), 4619–4629. doi:10.1128/JVI.02406-09

Coutard, B., Valle, C., de Lamballerie, X., Canard, B., Seidah, N. G., & Decroly, E. (2020). The spike glycoprotein of the new coronavirus 2019-nCoV contains a furin-like cleavage site absent in CoV of the same clade. Antiviral Res, 176, 104742. doi:10.1016/j.antiviral.2020.104742

Decroly, E., Debarnot, C., Ferron, F., Bouvet, M., Coutard, B., Imbert, I., … Canard, B. (2011). Crystal structure and functional analysis of the SARS-coronavirus RNA cap 2’-O-methyltransferase nsp10/nsp16 complex. PLoS Pathog, 7(5), e1002059. doi:10.1371/journal.ppat.1002059

DeDiego, M. L., Alvarez, E., Almazan, F., Rejas, M. T., Lamirande, E., Roberts, A., … Enjuanes, L. (2007). A severe acute respiratory syndrome coronavirus that lacks the E gene is attenuated in vitro and in vivo. J Virol, 81(4), 1701–1713. doi:10.1128/JVI.01467-06

DeLano, W. L. (2002). The PyMOL molecular graphics system. Retrieved from http://www.pymol.org

Denison, M. R., Graham, R. L., Donaldson, E. F., Eckerle, L. D., & Baric, R. S. (2011). Coronaviruses. RNA Biology,, 8(2), 270–279. doi:10.4161/rna.8.2.15013

DiMaio, F., Leaver-Fay, A., Bradley, P., Baker, D., & Andre, I. (2011). Modeling symmetric macromolecular structures in Rosetta3. PLoS ONE, 6(6), e20450. doi:10.1371/journal.pone.0020450

Eckerle, L. D., Becker, M. M., Halpin, R. A., Li, K., Venter, E., Lu, X., … Denison, M. R. (2010). Infidelity of SARS-CoV Nsp14-exonuclease mutant virus replication is revealed by complete genome sequencing. PLoS Pathog, 6(5), e1000896. doi:10.1371/journal.ppat.1000896

Elbe, S., & Buckland-Merrett, G. (2017). Data, disease and diplomacy: GISAID’s innovative contribution to global health. Glob Chall, 1(1), 33–46. doi:10.1002/gch2.1018

Faure, G., & Koonin, E. V. (2015). Universal distribution of mutational effects on protein stability, uncoupling of protein robustness from sequence evolution and distinct evolutionary modes of prokaryotic and eukaryotic proteins. Phys Biol, 12(3), 035001. doi:10.1088/1478-3975/12/3/035001

Frick, D. N., Virdi, R. S., Vuksanovic, N., Dahal, N., & Silvaggi, N. R. (2020). Molecular Basis for ADP-Ribose Binding to the Mac1 Domain of SARS-CoV-2 nsp3. Biochemistry, 59(28), 2608–2615. doi:10.1021/acs.biochem.0c00309

Gao, X., Qin, B., Chen, P., Zhu, K., Hou, P., Wojdyla, J. A., … Cui, S. (2020). Crystal structure of SARS-CoV-2 papain-like protease. Acta Pharm Sin B. doi:10.1016/j.apsb.2020.08.014

Gao, Y., Yan, L., Huang, Y., Liu, F., Zhao, Y., Cao, L., … Rao, Z. (2020). Structure of the RNA-dependent RNA polymerase from COVID-19 virus. Science, 368(6492), 779–782. doi:10.1126/science.abb7498

Goodsell, D. S., Autin, L., & Olson, A. J. (2019). Illustrate: Software for Biomolecular Illustration. Structure, 27, 1716–1720. doi:10.1016/j.str.2019.08.011

Goodsell, D. S., Zardecki, C., Di Costanzo, L., Duarte, J. M., Hudson, B. P., Persikova, I., … Burley, S. K. (2020). RCSB Protein Data Bank: Enabling biomedical research and drug discovery. Protein Sci, 29, 52–65. doi:10.1002/pro.3730

Gordon, D. E., Jang, G. M., Bouhaddou, M., Xu, J., Obernier, K., White, K. M., … Krogan, N. J. (2020). A SARS-CoV-2 protein interaction map reveals targets for drug repurposing. Nature, 583(7816), 459–468. doi:10.1038/s41586-020-2286-9

Hadfield, J., Megill, C., Bell, S. M., Huddleston, J., Potter, B., Callender, C., … Neher, R. A. (2018). Nextstrain: real-time tracking of pathogen evolution. Bioinformatics, 34(23), 4121–4123. doi:10.1093/bioinformatics/bty407

Henderson, R., Edwards, R. J., Mansouri, K., Janowska, K., Stalls, V., Gobeil, S. M. C., … Acharya, P. (2020). Controlling the SARS-CoV-2 spike glycoprotein conformation. Nat Struct Mol Biol, 27(10), 925–933. doi:10.1038/s41594-020-0479-4

Hillen, H. S., Kokic, G., Farnung, L., Dienemann, C., Tegunov, D., & Cramer, P. (2020). Structure of replicating SARS-CoV-2 polymerase. Nature, 584(7819), 154–156. doi:10.1038/s41586-020-2368-8

Hoffmann, M., Kleine-Weber, H., Schroeder, S., Kruger, N., Herrler, T., Erichsen, S., … Pohlmann, S. (2020). SARS-CoV-2 Cell Entry Depends on ACE2 and TMPRSS2 and Is Blocked by a Clinically Proven Protease Inhibitor. Cell, 181(2), 271–280 e278. doi:10.1016/j.cell.2020.02.052

Huang, C., Wei, P., Fan, K., Liu, Y., & Lai, L. (2004). 3C-like proteinase from SARS coronavirus catalyzes substrate hydrolysis by a general base mechanism. Biochemistry, 43(15), 4568–4574. doi:10.1021/bi036022q

Humphrey, W., Dalke, A., & Schulten, K. (1996). VMD: visual molecular dynamics. J Mol Graph, 14(1), 33–38. Retrieved from http://www.ncbi.nlm.nih.gov/entrez/query.fcgi?cmd=Retrieve&db=PubMed&dopt=Citation&list_uids=8744570

Jia, Z., Yan, L., Ren, Z., Wu, L., Wang, J., Guo, J., … Rao, Z. (2019). Delicate structural coordination of the Severe Acute Respiratory Syndrome coronavirus Nsp13 upon ATP hydrolysis. Nucleic Acids Res, 47(12), 6538–6550. doi:10.1093/nar/gkz409

Jin, Z., Du, X., Xu, Y., Deng, Y., Liu, M., Zhao, Y., … Yang, H. (2020). Structure of M(pro) from SARS-CoV-2 and discovery of its inhibitors. Nature, 582(7811), 289–293. doi:10.1038/s41586-020-2223-y

Johnson, B. A., Xie, X., Kalveram, B., Lokugamage, K. G., Muruato, A., Zou, J., … Menachery, V. D. (2020). Furin Cleavage Site Is Key to SARS-CoV-2 Pathogenesis. bioRxiv. doi:10.1101/2020.08.26.268854

Kang, S., Yang, M., Hong, Z., Zhang, L., Huang, Z., Chen, X., … Chen, S. (2020). Crystal structure of SARS-CoV-2 nucleocapsid protein RNA binding domain reveals potential unique drug targeting sites. Acta Pharm Sin B, 10(7), 1228–1238. doi:10.1016/j.apsb.2020.04.009

Kellogg, E. H., Leaver-Fay, A., & Baker, D. (2011). Role of conformational sampling in computing mutation-induced changes in protein structure and stability. Proteins, 79(3), 830–838. doi:10.1002/prot.22921

Kern, D. M., Sorum, B., Hoel, C. M., Sridharan, S., Remis, J. P., Toso, D. B., & Brohawn, S. G. (2020). Cryo-EM structure of the SARS-CoV-2 3a ion channel in lipid nanodiscs. bioRxiv. doi:10.1101/2020.06.17.156554

Khailany, R. A., Safdar, M., & Ozaslan, M. (2020). Genomic characterization of a novel SARS-CoV-2. Gene Rep, 19, 100682. doi:10.1016/j.genrep.2020.100682

Kim, D., Lee, J. Y., Yang, J. S., Kim, J. W., Kim, V. N., & Chang, H. (2020). The Architecture of SARS-CoV-2 Transcriptome. Cell, 181(4), 914–921 e910. doi:10.1016/j.cell.2020.04.011

Kim, Y., Jedrzejczak, R., Maltseva, N. I., Wilamowski, M., Endres, M., Godzik, A., … Joachimiak, A. (2020). Crystal structure of Nsp15 endoribonuclease NendoU from SARS-CoV-2. Protein Sci, 29(7), 1596–1605. doi:10.1002/pro.3873

Korber, B., Fischer, W. M., Gnanakaran, S., Yoon, H., Theiler, J., Abfalterer, W., … Montefiori, D. C. (2020). Tracking Changes in SARS-CoV-2 Spike: Evidence that D614G Increases Infectivity of the COVID-19 Virus. Cell, 182(4), 812–827 e819. doi:10.1016/j.cell.2020.06.043

Lan, J., Ge, J., Yu, J., Shan, S., Zhou, H., Fan, S., … Wang, X. (2020). Structure of the SARS-CoV-2 spike receptor-binding domain bound to the ACE2 receptor. Nature, 581(7807), 215–220. doi:10.1038/s41586-020-2180-5

Littler, D. R., Gully, B. S., Colson, R. N., & Rossjohn, J. (2020). Crystal Structure of the SARS-CoV-2 Non-structural Protein 9, Nsp9. iScience, 23(7) 101258. doi:10.1016/j.isci.2020.101258

Ma, Y., Wu, L., Shaw, N., Gao, Y., Wang, J., Sun, Y., … Rao, Z. (2015). Structural basis and functional analysis of the SARS coronavirus nsp14-nsp10 complex. Proc Natl Acad Sci U S A, 112(30), 9436–9441. doi:10.1073/pnas.1508686112

MacKerell Jr., A. D., Bashford, D., Bellott, M., Dunbrack, R. L., Evanseck, J. D., Field, M. J., … Karplus, M. (1998). All-atom empirical potential for molecular modeling and dynamics studies of proteins. J Phys Chem B, 102(18), 3586–3616. doi:10.1021/jp973084f

Melero, R., Sorzano, C. O. S., Foster, B., Vilas, J. L., Martinez, M., Marabini, R., … Carazo, J. M. (2020). Continuous flexibility analysis of SARS-CoV-2 Spike prefusion structures. bioRxiv. doi:10.1101/2020.07.08.191072

Miao, Z., Tidu, A., Eriani, G., & Martin, F. (2020). Secondary structure of the SARS-CoV-2 5’-UTR. RNA Biol, 1–10. doi:10.1080/15476286.2020.1814556

Michalska, K., Kim, Y., Jedrzejczak, R., Maltseva, N. I., Stols, L., Endres, M., and Joachimiak, A. (2020). The crystal structure of papain-like protease of SARS CoV-2. doi: 10.2210/pdb6W9C/pdb.

Minasov, G., Shuvalova, L., Rosas-Lemus, M., Kiryukhina, O., Brunzelle, J. S., Satchell, K. J. F., & Center for Structural Genomics of Infectious Diseases (CSGID). (2020). Crystal Structure of Nsp16-Nsp10 from SARS-CoV-2 in Complex with 7-methyl-GpppA and S-Adenosylmethionine. doi: 10.2210/pdb6WVN/pdb.

Neuman, B. W., Kiss, G., Kunding, A. H., Bhella, D., Baksh, M. F., Connelly, S., … Buchmeier, M. J. (2011). A structural analysis of M protein in coronavirus assembly and morphology. J Struct Biol, 174(1), 11–22. doi:10.1016/j.jsb.2010.11.021

Newman, J. A., Yosaatmadja, Y., Douangamath, A., Arrowsmith, C.H., von Delft, F., Edwards, A., Bountra, C., Gileadi, O. (2020). Crystal structure of the SARS-CoV-2 helicase at 1.94 Angstrom resolution. 10.2210/pdb6ZSL/pdb.

Nisthal, A., Wang, C. Y., Ary, M. L., & Mayo, S. L. (2019). Protein stability engineering insights revealed by domain-wide comprehensive mutagenesis. Proc Natl Acad Sci U S A, 116(33), 16367–16377. doi:10.1073/pnas.1903888116

Phillips, J. C., Hardy, D. J., Maia, J. D. C., Stone, J. E., Ribeiro, J. V., Bernardi, R. C., … Tajkhorshid, E. (2020). Scalable molecular dynamics on CPU and GPU architectures with NAMD. J Chem Phys, 153(4), 044130. doi:10.1063/5.0014475

Pollack, A. (2003). Company Says It Mapped Part of SARS Virus. The New York Times, July 30, 2003, C2.

Privalov, P. L., & Gill, S. J. (1988). Stability of protein structure and hydrophobic interaction. Adv Protein Chem, 39, 191–234. doi:10.1016/s0065-3233(08)60377-0

Protein Data Bank. (1971). Crystallography: Protein Data Bank. Nature (London), New Biol., 233(42), 223–223. doi:10.1038/newbio233223b0

Ratia, K., Saikatendu, K. S., Santarsiero, B. D., Barretto, N., Baker, S. C., Stevens, R. C., & Mesecar, A. D. (2006). Severe acute respiratory syndrome coronavirus papain-like protease: structure of a viral deubiquitinating enzyme. Proc Natl Acad Sci U S A, 103(15), 5717–5722. doi:10.1073/pnas.0510851103

Razban, R. M., & Shakhnovich, E. I. (2020). Effects of Single Mutations on Protein Stability Are Gaussian Distributed. Biophys J, 118(12), 2872–2878. doi:10.1016/j.bpj.2020.04.027

Rogstam, A., Nyblom, M., Christensen, S., Sele, C., Talibov, V. O., Lindvall, T., … Kozielski, F. (2020). Crystal Structure of Non-Structural Protein 10 from Severe Acute Respiratory Syndrome Coronavirus-2. Int J Mol Sci, 21(19). doi:10.3390/ijms21197375

Rosas-Lemus, M., Minasov, G., Shuvalova, L., Inniss, N. L., Kiryukhina, O., Wiersum, G., … Satchell, K. J. F. (2020). The crystal structure of nsp10-nsp16 heterodimer from SARS-CoV-2 in complex with S-adenosylmethionine. bioRxiv, 2020.2004.2017.047498. doi:10.1101/2020.04.17.047498

Ruch, T. R., & Machamer, C. E. (2012). The coronavirus E protein: assembly and beyond. Viruses, 4(3), 363–382. doi:10.3390/v4030363

Rut, W., Lv, Z., Zmudzinski, M., Patchett, S., Nayak, D., Snipas, S. J., … Olsen, S. K. (2020). Activity profiling and crystal structures of inhibitor-bound SARS-CoV-2 papain-like protease: A framework for anti-COVID-19 drug design. Sci Adv, 6(42). doi:10.1126/sciadv.abd4596

Schoeman, D., & Fielding, B. C. (2019). Coronavirus envelope protein: current knowledge. Virol J, 16(1), 69. doi:10.1186/s12985-019-1182-0

Sehnal, D., Rose, A., Koca, J., Burley, S., & Velankar, S. (2018). Mol‪: Towards a Common Library and Tools for Web Molecular Graphics. MolVa: Workshop on Molecular Graphics and Visual Analysis of Molecular Data 2018. doi:10.2312/molva.20181103

Serrano, P., Johnson, M. A., Almeida, M. S., Horst, R., Herrmann, T., Joseph, J. S., … Wuthrich, K. (2007). Nuclear magnetic resonance structure of the N-terminal domain of nonstructural protein 3 from the severe acute respiratory syndrome coronavirus. J Virol, 81(21), 12049–12060. doi:10.1128/JVI.00969-07

Serrano, P., Johnson, M. A., Chatterjee, A., Neuman, B. W., Joseph, J. S., Buchmeier, M. J., … Wuthrich, K. (2009). Nuclear magnetic resonance structure of the nucleic acid-binding domain of severe acute respiratory syndrome coronavirus nonstructural protein 3. J Virol, 83(24), 12998–13008. doi:10.1128/JVI.01253-09

Shannon, A., Le, N. T., Selisko, B., Eydoux, C., Alvarez, K., Guillemot, J. C., … Canard, B. (2020). Remdesivir and SARS-CoV-2: Structural requirements at both nsp12 RdRp and nsp14 Exonuclease active-sites. Antiviral Res, 178, 104793. doi:10.1016/j.antiviral.2020.104793

Shin, D., Mukherjee, R., Grewe, D., Bojkova, D., Baek, K., Bhattacharya, A., … Dikic, I. (2020). Papain-like protease regulates SARS-CoV-2 viral spread and innate immunity. Nature. doi:10.1038/s41586-020-2601-5

Shiryaev, S. A., Chernov, A. V., Golubkov, V. S., Thomsen, E. R., Chudin, E., Chee, M. S., … Cieplak, P. (2013). High-resolution analysis and functional mapping of cleavage sites and substrate proteins of furin in the human proteome. PLoS ONE, 8(1), e54290. doi:10.1371/journal.pone.0054290

Shu, Y., & McCauley, J. (2017). GISAID: Global initiative on sharing all influenza data - from vision to reality. Euro Surveill, 22(13). doi:10.2807/1560-7917.ES.2017.22.13.30494

Simmonds, P., Bukh, J., Combet, C., Deleage, G., Enomoto, N., Feinstone, S., … Widell, A. (2005). Consensus proposals for a unified system of nomenclature of hepatitis C virus genotypes. Hepatology, 42(4), 962–973. doi:10.1002/hep.20819

Siu, Y. L., Teoh, K. T., Lo, J., Chan, C. M., Kien, F., Escriou, N., … Nal, B. (2008). The M, E, and N structural proteins of the severe acute respiratory syndrome coronavirus are required for efficient assembly, trafficking, and release of virus-like particles. J Virol, 82(22), 11318–11330. doi:10.1128/JVI.01052-08

Surya, W., Li, Y., & Torres, J. (2018). Structural model of the SARS coronavirus E channel in LMPG micelles. Biochim Biophys Acta Biomembr, 1860(6), 1309–1317. doi:10.1016/j.bbamem.2018.02.017

Tan, J., Vonrhein, C., Smart, O. S., Bricogne, G., Bollati, M., Kusov, Y., … Hilgenfeld, R. (2009). The SARS-unique domain (SUD) of SARS coronavirus contains two macrodomains that bind G-quadruplexes. PLoS Pathog, 5(5), e1000428. doi:10.1371/journal.ppat.1000428

Tang, T., Bidon, M., Jaimes, J. A., Whittaker, G. R., & Daniel, S. (2020). Coronavirus membrane fusion mechanism offers a potential target for antiviral development. Antiviral Res, 178, 104792. doi:10.1016/j.antiviral.2020.104792

Tokuriki, N., Stricher, F., Schymkowitz, J., Serrano, L., & Tawfik, D. S. (2007). The stability effects of protein mutations appear to be universally distributed. J Mol Biol, 369(5), 1318–1332. doi:10.1016/j.jmb.2007.03.069

Tseng, Y. T., Chang, C. H., Wang, S. M., Huang, K. J., & Wang, C. T. (2013). Identifying SARS-CoV membrane protein amino acid residues linked to virus-like particle assembly. PLoS ONE, 8(5), e64013. doi:10.1371/journal.pone.0064013

Verdia-Baguena, C., Nieto-Torres, J. L., Alcaraz, A., DeDiego, M. L., Torres, J., Aguilella, V. M., & Enjuanes, L. (2012). Coronavirus E protein forms ion channels with functionally and structurally-involved membrane lipids. Virology, 432(2), 485–494. doi:10.1016/j.virol.2012.07.005

Voss, D., Pfefferle, S., Drosten, C., Stevermann, L., Traggiai, E., Lanzavecchia, A., & Becker, S. (2009). Studies on membrane topology, N-glycosylation and functionality of SARS-CoV membrane protein. Virol J, 6, 79. doi:10.1186/1743-422X-6-79

Walls, A. C., Park, Y. J., Tortorici, M. A., Wall, A., McGuire, A. T., & Veesler, D. (2020). Structure, Function, and Antigenicity of the SARS-CoV-2 Spike Glycoprotein. Cell, 181(2), 281–292 e286. doi:10.1016/j.cell.2020.02.058

Wang, Q., Wu, J., Wang, H., Gao, Y., Liu, Q., Mu, A., … Rao, Z. (2020). Structural Basis for RNA Replication by the SARS-CoV-2 Polymerase. Cell, 182(2), 417–428 e413. doi:10.1016/j.cell.2020.05.034

Wang, Q., Zhang, Y., Wu, L., Niu, S., Song, C., Zhang, Z., … Qi, J. (2020). Structural and Functional Basis of SARS-CoV-2 Entry by Using Human ACE2. Cell, 181(4), 894–904 e899. doi:10.1016/j.cell.2020.03.045

Wang, R., Hozumi, Y., Yin, C., & Wei, G. W. (2020). Decoding SARS-CoV-2 Transmission and Evolution and Ramifications for COVID-19 Diagnosis, Vaccine, and Medicine. J Chem Inf Model. doi:10.1021/acs.jcim.0c00501

Watanabe, Y., Allen, J. D., Wrapp, D., McLellan, J. S., & Crispin, M. (2020). Site-specific glycan analysis of the SARS-CoV-2 spike. Science, 369(6501), 330–333. doi:10.1126/science.abb9983

Waterhouse, A., Bertoni, M., Bienert, S., Studer, G., Tauriello, G., Gumienny, R., … Schwede, T. (2018). SWISS-MODEL: homology modelling of protein structures and complexes. Nucleic Acids Res, 46(W1), W296–W303. doi:10.1093/nar/gky427

Westbrook, J. D., Soskind, R., Hudson, B. P., & Burley, S. K. (2020). Impact of Protein Data Bank on Anti-neoplastic Approvals. Drug Discov Today, 25, 837–850 doi:10.1016/j.drudis.2020.02.002

Wrapp, D., Wang, N., Corbett, K. S., Goldsmith, J. A., Hsieh, C. L., Abiona, O., … McLellan, J. S. (2020). Cryo-EM structure of the 2019-nCoV spike in the prefusion conformation. Science, 367(6483), 1260–1263. doi:10.1126/science.abb2507

Wu, F., Zhao, S., Yu, B., Chen, Y. M., Wang, W., Song, Z. G., … Zhang, Y. Z. (2020). A new coronavirus associated with human respiratory disease in China. Nature, 579(7798), 265–269. doi:10.1038/s41586-020-2008-3

wwPDB consortium. (2019). Protein Data Bank: the single global archive for 3D macromolecular structure data. Nucleic Acids Res, 47(D1), D520–D528. doi:10.1093/nar/gky949

Xia, S., Liu, M., Wang, C., Xu, W., Lan, Q., Feng, S., … Lu, L. (2020). Inhibition of SARS-CoV-2 (previously 2019-nCoV) infection by a highly potent pan-coronavirus fusion inhibitor targeting its spike protein that harbors a high capacity to mediate membrane fusion. Cell Res, 30(4), 343–355. doi:10.1038/s41422-020-0305-x

Xing, Y., Li, X., Gao, X., & Dong, Q. (2020). Natural Polymorphisms Are Present in the Furin Cleavage Site of the SARS-CoV-2 Spike Glycoprotein. Front Genet, 11, 783. doi:10.3389/fgene.2020.00783

Yan, R., Zhang, Y., Li, Y., Xia, L., Guo, Y., & Zhou, Q. (2020). Structural basis for the recognition of SARS-CoV-2 by full-length human ACE2. Science, 367(6485), 1444–1448. doi:10.1126/science.abb2762

Yin, W., Mao, C., Luan, X., Shen, D.-D., Shen, Q., Su, H., … Xu, H. E. (2020). Structural basis for inhibition of the RNA-dependent RNA polymerase from SARS-CoV-2 by remdesivir. Science, 368(6498), 1499–1504. doi:10.1126/science.abc1560

Yurkovetskiy, L., Wang, X., Pascal, K. E., Tomkins-Tinch, C., Nyalile, T. P., Wang, Y., … Luban, J. (2020). Structural and Functional Analysis of the D614G SARS-CoV-2 Spike Protein Variant. Cell, 183(3), 739–751 e738. doi:10.1016/j.cell.2020.09.032

Zhang, L., Jackson, C. B., Mou, H., Ojha, A., Rangarajan, E. S., Izard, T., … Choe, H. (2020). The D614G mutation in the SARS-CoV-2 spike protein reduces S1 shedding and increases infectivity. bioRxiv. doi:10.1101/2020.06.12.148726

Zinzula, L., Basquin, J., Bohn, S., Beck, F., Klumpe, S., Pfeifer, G., … Baumeister, W. (2020). High-resolution structure and biophysical characterization of the nucleocapsid phosphoprotein dimerization domain from the Covid-19 severe acute respiratory syndrome coronavirus 2. Biochem Biophys Res Commun. doi:10.1016/j.bbrc.2020.09.131

