## Supplementary descriptions, figures, tables, and models for "Evolution of the SARS-CoV-2 proteome in three dimensions (3D) during the first six months of the COVID-19 pandemic"

Table of Contents

1. Individual Commentaries for Remaining 16 SARS-CoV-2 Study Proteins, including nsp1; nsp2; nsp3 Overview and nsp3 components: nsp3a, nsp3b, nsp3c, nsp3e, and UNK; nsp4; nsp6; nsp9; nsp15; Orf3a; Orf6; Orf7a; Orf7b; and Orf8.

2. Legends for Supplementary Figures for 29 SARS-CoV-2 Study Proteins.

3. Legends for Supplementary Tables for 29 SARS-CoV-2 Study Proteins.

4. Description of Computed Structural Models for Unique Sequence Variants for 29 SARS-CoV-2 Study Proteins.

5. Supplementary Materials References

### 1. Individual Commentaries for Remaining 16 SARS-CoV-2 Study Proteins

Non-structural protein 1 (nsp1): nsp1 is a 180-residue protein expressed at the N-terminus of polyproteins 1a and 1ab, from which it is excised by PLPro. During coronavirus infections, nsp1 blocks host cell protein translation by binding to the mRNA entry tunnel of the 40S ribosomal subunit, stalling host mRNA translation and inducing degradation of host mRNAs (Kamitani, Huang, Narayanan, Lokugamage, & Makino, 2009; Lokugamage, Narayanan, Huang, & Makino, 2012). nsp1 interactions with a conserved segment of the 5′ untranslated region (UTR) of viral genome are thought to prevent shutdown of viral protein expression through an as yet uncharacterized mechanism(s) (Huang et al., 2011). At the time of writing, there were no experimental structures of full-length SARS-CoV-2 nsp1 available in the PDB. Our evolutionary analysis was carried out with a homology model computed using an NMR structure of the ~85% identical SARS-CoV-1nsp1 protein (PDB ID 2HSX (Almeida, Johnson, Herrmann, Geralt, & Wuthrich, 2007)), which provided coverage of residues 13-127 of SARS-CoV-2 nsp1. This segment of nsp1 is folded into a single, compact $\alpha/\beta$ domain (Supplementary Figure nsp1). A conserved Arginine residue (R124) contributes to host RNA degradation in SARS-CoV-1 (Tanaka, Kamitani, DeDiego, Enjuanes, & Matsuura, 2012) and may serve the same function in SARS-CoV-2. Experimental structure information was also available for nsp1 C-terminal residues 148-180 observed in a 3DEM structure of SARS-CoV-2 nsp1 bound to the 40S ribosomal subunit (PDB ID 6ZLW (Thoms et al., 2020)). Residues 148-180 of nsp1 form a two $\alpha$-helix hairpin that plugs the mRNA entry tunnel, interacting with both ribosomal proteins (*i.e.*, uS5 and uS3) and the ribosomal RNA (*i.e.*, rRNA helix h18). Within this C-terminal segment, K164 and H165 are essential for ribosome binding (Jauregui, Savalia, Lowry, Farrell, & Wathelet, 2013). The globular portion of nsp1 was not well resolved in the 3DEM electric potential map.

Overall substitution trends for nsp1 and energetics analysis results are summarized in Tables 1 and 2. D75E (conservative; surface) was the most common substitution (GISAID dataset count=332). D75E was also found in the context of 4 double mutant USVs. R124C (non-conservative; boundary), which was the third most common substitution (GISAID dataset count=35), is of interest because it is thought to play a role in host cell mRNA degradation (Tanaka et al., 2012). Substitutions were observed in 53% (17/32) of the residues comprising the C-terminal ribosome binding segment (residues 148-180). USVs affecting the C-terminal region included L149F, L149F/H/R, G150C/D, D152G, Q158H, E159G/K, I95V/N160K, N162Y, T163I, K164I, S166G, S167I, G168V, V169A, E172K/Q, R175C/G/H, E176G, and G179R. [N.B.: Structural information was not available for L149 in PDB ID 6ZME.] The most frequently substituted of the C-terminal residues were L149 and R175, each occurring in 3 USVs. Of the C-terminal residues that underwent substitution, two interact directly with ribosomal proteins: D152G (binding to uS3 residue 116) and Q158H (binding to uS2 residue 147). Three of the substituted residues interact directly with the 18S ribosomal RNA: K164 (binding to *C624, G625, A629, and U630; italics denote RNA bases*), S167 (binding to *G600*), and R175 (binding to *A605* and *G606*).

Supplementary Figure nsp1. Ribbon/ball-and-stick figure representation of the computed structural model of nsp1 (N- and C-termini labeled). C-terminal residues 148-180 responsible for binding to the 40S ribosomal subunit are colored orange. The most frequently substituted residues D75 and R124 are color coded as follows: C-green, N-blue, O-red). [N.B.: The conformation of residues 148-180 shown differs from that observed when the protein binds the ribosome.]

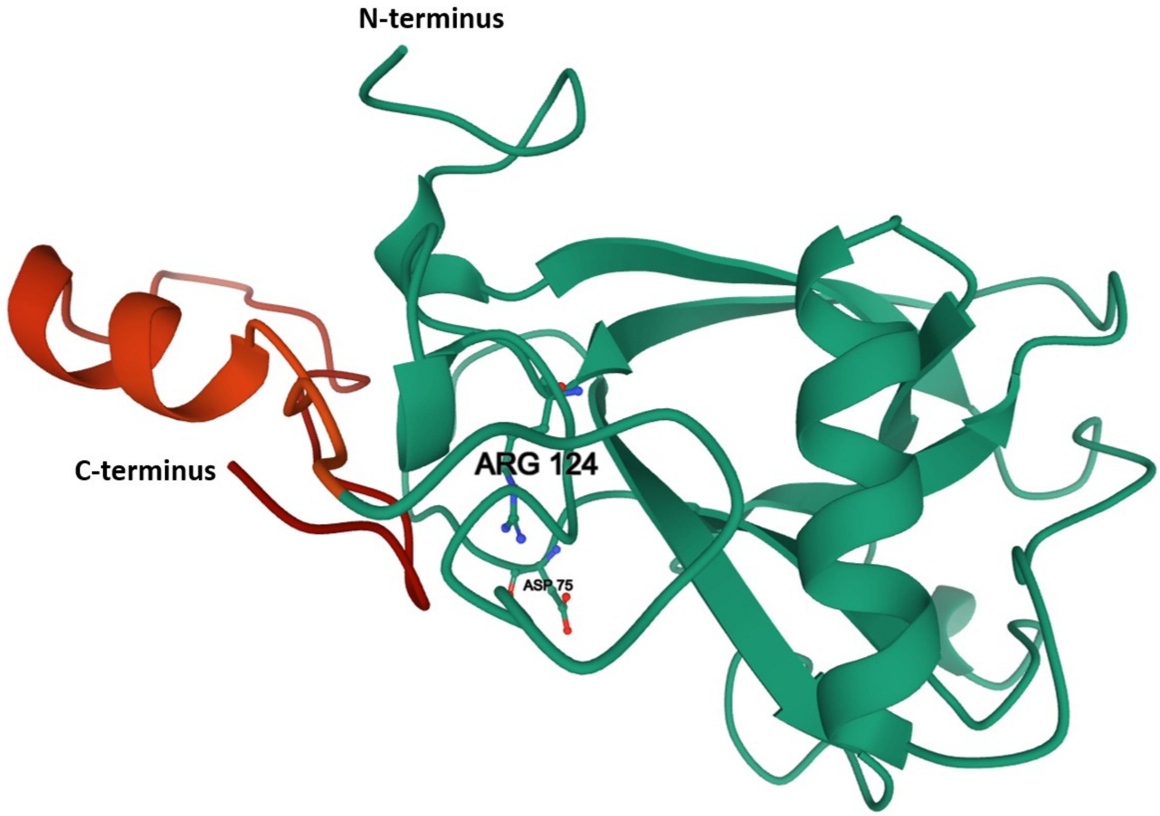

Non-structural protein 2 (nsp2): nsp2 is a 638-residue protein that is translated within pp1ab, from which it is excised by PLPro. At present, the function of SARS-CoV-2 nsp2 is not known. Recent bioinformatics analyses have predicted that the protein possesses four membrane spanning $\alpha$-helical segments (Angeletti et al., 2020). Deletion of nsp2 from SARS-CoV-1 and murine hepatitis virus (MHV) revealed that it is not essential for viral replication but instead plays a role in sustaining infection. (Graham, Sims, Brockway, Baric, & Denison, 2005). The nsp2 structural model used for analyzing evolution in 3D was computed by the David Baker Laboratory during a CASP competition (CASP-C1901 Stage 2) (Supplementary Figure nsp2).

Overall substitution trends for nsp2 and energetics analysis results are summarized in Tables 1 and 2. T85I (non-conservative, boundary) is the most common substitution, observed 7,763 times in the GISAID dataset and in 205 USVs. The dual substitution F10L/T85I also occurred frequently (GISAID dataset count=150). The F10L substitution did not appear in a single-mutant USV, but instead only appeared in concert with T85I in 8 USVs. I559V (conservative, boundary) and P585S (non-conservative, boundary) appeared in concert in 39 USVs (GISAID dataset count=1,472).

Supplementary Figure nsp2. Ribbon representation of the computed structural model for nsp2 rainbow colored from N- (blue) to C-terminus (red).

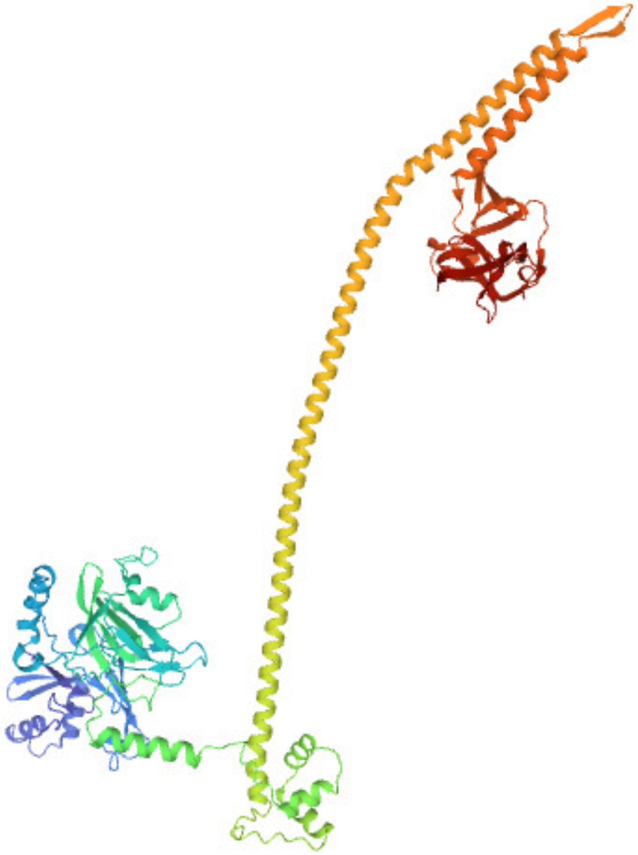

Non-structural protein 3 (nsp3) Overview: nsp3 is a large multidomain protein (Lei, Kusov, & Hilgenfeld, 2018)) that is excised from polyproteins 1a and ab by PLPro. Eight nsp3 domains are conserved among all coronaviruses and appear to support similar biological/biochemical functions (Serrano et al., 2007). To simplify our work on nsp3, the polypeptide chain was subdivided into 6 segments, identified in order of occurrence as nsp3a (residues 819-1024), nsp3b (1025-1230), nsp3c (1231-1562), PLPro (1563-1905), nsp3e (1906-2077), and UNK (2078-2763). Excluding UNK, the other five segments of nsp3 are found in the cytoplasm. UNK possesses two transmembrane regions that insert the polypeptide chain into the double membrane vesicle structure and return the C-terminus of nsp3 to the cytoplasm. The results of our evolutionary analyses carried out separately for nsp3a, nsp3b, nsp3c, nsp3e, and UNK are summarized below (see main text for discussion of PLPro).

Non-structural protein 3 (nsp3a): nsp3a, the N-terminal portion of nsp3, is 206 residues in length. It is responsible for RNA binding and processing and is conserved across all coronaviruses (Serrano et al., 2007). Nsp3a includes the ubiquitin-like domain 1 (Ubl1) and the Glu-rich acidic region. SARS-CoV-1 nsp3a binds to single-stranded RNA AUA sequences, which appear frequently in the 5' untranslated regions of the viral genome. The $\beta$1 and $\alpha$1 regions of nsp3a have positively charged surfaces that are highly conserved among coronaviruses strongly indicative of RNA binding function (Serrano et al., 2007). SARS-CoV-2 nsp3a is also highly similar to its Mouse Hepatitis Virus (MHV) homolog, which binds to the nucleocapsid protein (N-protein), indicating that this domain is also important during viral infection. Acidic residues within the $\alpha$2 helix of MHV nsp3a (PDB ID 2M0A (Keane & Giedroc, 2013)); corresponding to residues 868-882 of SARS-CoV-2 nsp3a) interact with the serine and arginine rich region (SR-rich region) of the N-protein (Serrano et al., 2007). The structure of SARS-CoV-2 nsp3a has not been elucidated experimentally. The structural model computed using the SARS-CoV-1 homolog (sequence identity: ~79%) NMR structure (PDB ID 2IDY (Serrano et al., 2007)) was used for our evolutionary analyses (Supplementary Figure nsp3a).

Overall substitution trends for nsp3a and energetics analysis results are summarized in Tables 1 and 2. A876T (non-conservative, boundary) is the most common USV (GISAID dataset count=532). The following basic residues have the potential for RNA interactions: K822, K837, R848, K851, K856, K881, K915, and K958. Of those, only three underwent non-conservative substitutions: R848S, K851T, and K881M (while conservative K-R exchanges occurred at sites 822, 837, 848, 851, 865, and 858). One or more of the unchanged/conserved residues may participate in an essential interaction with RNA. Within the $\alpha$2 helix (residues 868-882), thought to interact with the basic SR-rich region of the N-protein, only one acidic residue E870D underwent substitution that was conservative (Supplementary Figure nsp3a).

Supplementary Figure nsp3a. Ribbon representation of the computed structural model for nsp3a with labeled C- and N-termini and α2 helix denoted within a dashed box.

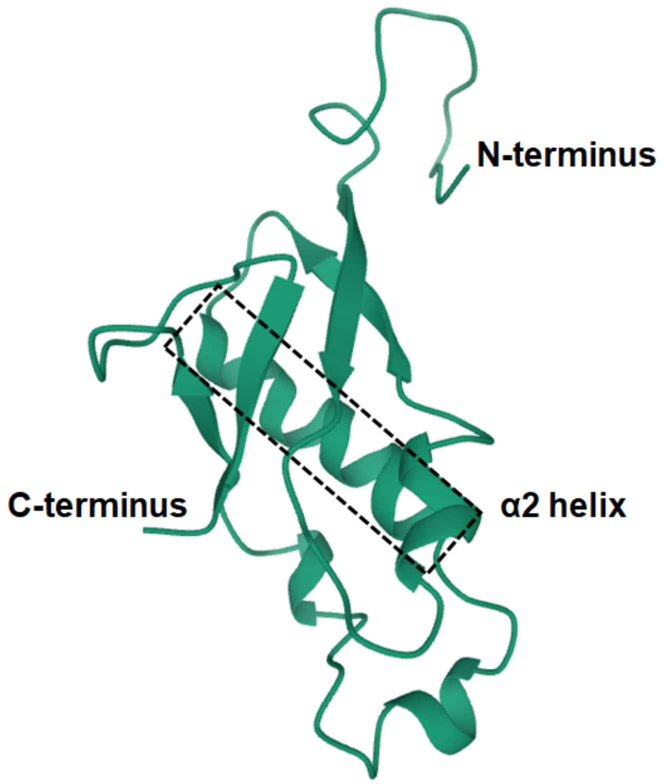

Non-structural protein 3 nsp3b: nsp3b, part of nsp3, is 206 residues in length (also known as “X”, Mac1, or Macro X domain, reviewed in (Lei et al., 2018)). Viral macrodomains are known to be key players in the viral-host arms race to control cell signaling in the innate immune response. They counteract host ADP-ribosyltransferase-mediated immune signaling by hydrolyzing ADP-ribose or poly(ADP-ribose) from host and viral proteins. However, the role played by nsp3b in coronaviral RNA replication is not well understood. For SARS-CoV-1, nsp3b is not essential for RNA replication in cell culture (Fehr et al., 2016); however, the domain appears to be involved in counteracting the host immune response in animal studies of coronavirus infections (Grunewald et al., 2019). Mutations in the catalytic machinery of MHV macrodomain result in a largely non-pathogenic virus (Fehr et al., 2016), implying that the hydrolytic activity of the domain may be critical -- making it an attractive target for drug discovery. Indeed, high throughput crystallographic screening efforts aimed at finding inhibitors of the SARS-CoV-2 macrodomain have recently been reported (Schuller et al., 2020). The experimental structure of SARS CoV-2 nsp3b (PDB ID 6WEY (Frick, Virdi, Vuksanovic, Dahal, & Silvaggi, 2020)) was used for our evolutionary analyses (Supplementary Figure nsp3b). Ligand-binding residues were identified using the experimental structure of the SARS-CoV-2 nsp3b-ADP-ribose complex (PDB ID 6WOJ (Alhammad et al., 2020)).

Overall substitution trends for nsp3b and energetics analysis results are summarized in Tables 1 and 2. D1036E (conservative, surface) is the most common substitution (GISAID dataset count=182). 12 residues are responsible for binding ADP-ribose, including D1044, I1045, N1062, H1067, G1069, V1071, A1072, P1147, S1150, G1152, I1153, and F1154. Nine of these residues underwent a total of 12 substitutions: I1045S/V, N1062S, H1067Y, G1069E/V, V1071I, P1147S, G1152S, I1153T/V, and F1154V.

Supplementary Figure nsp3b. Ribbon/stick figure representation of the experimental structure of nsp3b (PDB ID 6WEY (Frick et al., 2020)) with labeled N- and C- termini (left). Inset (right) shows the ADP-ribose binding pocket with selected residues highlighted (PDB ID 6WOJ (Alhammad et al., 2020)). Atomic stick figure color coding: C-green, O-red, N-blue, P-orange.

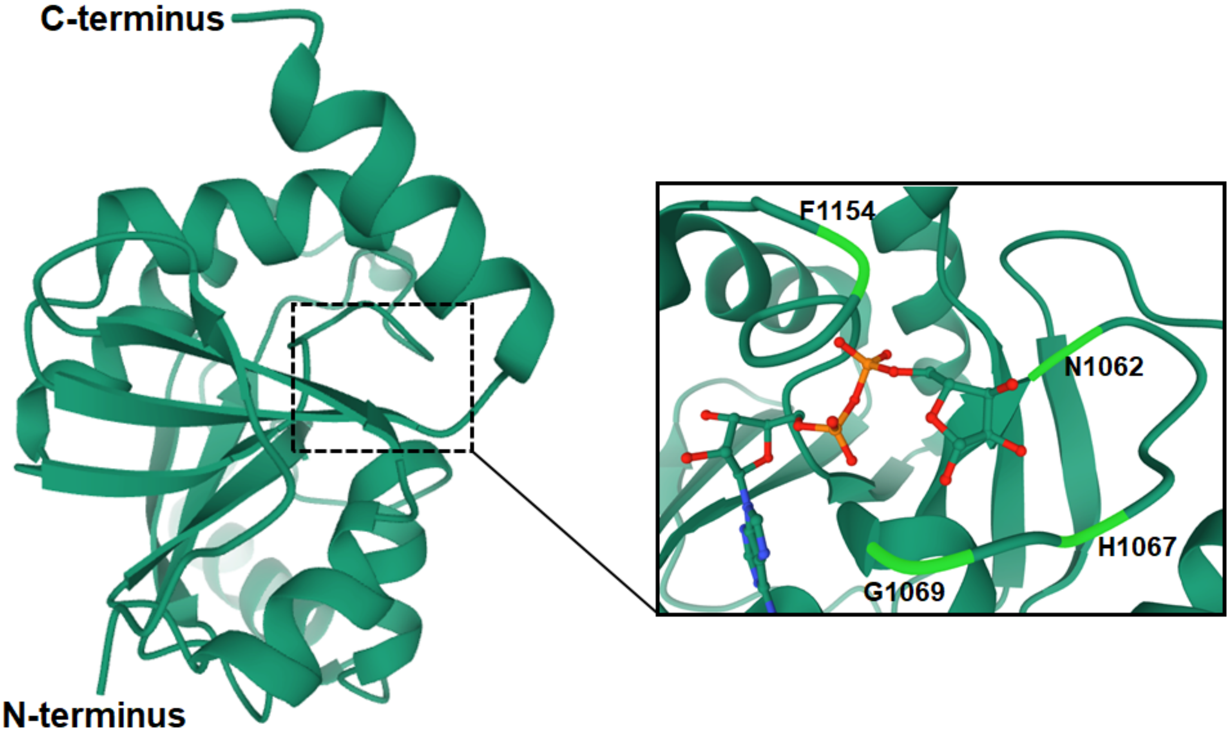

Non-structural protein 3 (nsp3c): nsp3c, part of nsp3, is 332 residues in length (reviewed in (Lei et al., 2018)). It contains the so-called SARS-unique domain or SUD (Snijder et al., 2003), which occurs only in SARS-CoV-2, SARS-CoV, and highly-related viruses found in bats. Our structural knowledge of nsp3c is currently restricted to the so-called "SUD(core)" of SARS-CoV-1 nsp3 (~76% identical to the corresponding segment of SARS-CoV-2 nsp3), which consists of two subdomains (SUD-N and SUD-M) that forms a homodimer (PDB ID 2W2G (Tan et al., 2009)). The SUD(core) is found in the cytosol. It binds to oligonucleotide G-quadruplexes, which suggests that it interacts with host-cell RNAs containing extended G tracts (*e.g.*, those found in the 3'-untranslated regions of mRNAs encoding proteins involved in signal transduction and apoptosis). In SARS-CoV-1, mutations of lysine residues on the surface showed reduction or abrogation of G-quadruplex binding (Tan et al., 2009). The nsp3c structural model used for analyzing evolution in 3D was computed by the David Baker Laboratory during a CASP competition (CASP-C1904 Stage 2, Supplementary Figure nsp3c).

Overall substitution trends for nsp3c and energetics analysis results are summarized in Tables 1 and 2. T1246I (non-conservative, core) is the most common substitution (GISAID dataset count=495). T1246I is found in 7 USVs, including 5 double-substitutions and 1 triple-substitution. Lysine substitutions are of interest because of their potential impact on G-quadruplex binding. A total of 18 USVs with substitutions involving Lysine residues were observed, including K1231R, K1233E, K1247N, K1280N, K1281E, K1305N, K1315Q, K1319E, K1343R, K1347R, K1348N, K1410R, K1428E/R, K1512R, K1529R, K1533E, and K1557N. Two sites of substitutions mapping the homodimer interface were observed including G1307D/S/V and A1420S/T. While four of these substitutions have moderate effects on stability ($\Delta\Delta G^{App}$ ~-0.1 to +3.9), G1307V may produce more pronounced perturbation with predicted $\Delta\Delta G^{App}$~+16.3 REU. Finally, Proline substitutions L1249P and S1400P, occurring with an $\alpha$-helix, were predicted to be highly destabilizing ($\Delta\Delta G^{App}$ >20 REU), indicating that a backbone rearrangement may occur due to the substitution. Both of these substitutions were observed in multiple GISAID dataset sequences, and the latter was present in a double-substitution S1400P/T1444I.

Supplementary Figure nsp3c. Ribbon/stick figure representation of the computed structural model of nsp3c. Lysine sidechain atom color coding is as follows: invariant or conservative substituted residue-C purple, N-blue; non-conservative substituted residue-C purple, N-blue.

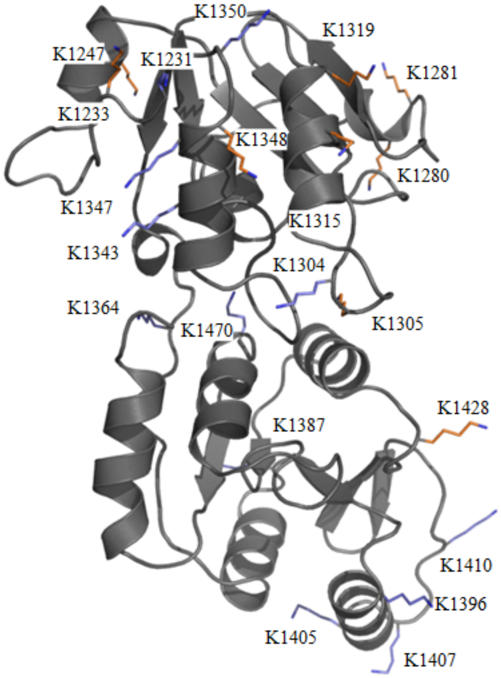

Non-structural protein 3 (nsp3e): nsp3e, part of nsp3, is 172 residues in length. It encompasses the nucleic-acid binding (NAB) and betacoronavirus-specific marker (βSM) domains. At the time of writing, there were no experimental structures of the SARS-CoV2 NAB or βSM domains in the PDB. The amino acid sequence to SARS-CoV-2 NAB domain is ~81% identical to that of the SARS-CoV-1 NAB domain, for which a PDB structure is available (PDB ID 2K87 (Serrano et al., 2009)). SARS-CoV-1 NAB is a compact α/β domain in which more than a dozen residues were shown using NMR spectroscopy to interact with single-stranded RNAs possessing GGG repeats. Sequence alignment of the two NABs permitted identification of four positively-charged residues in SARS-CoV-2 nsp3e that may be involved in RNA binding (K1980, K1981, K2000, R2011). Evolution of nsp3e in 3D was analyzed using a structural model computed by the David Baker Laboratory for CASP-C1904 Stage 2 (Supplementary Figure nsp3e).

Overall substitution trends for nsp3e and energetics analysis results are summarized in Tables 1 and 2. T2016K (non-conservative, surface) is the most common substitution (GISAID dataset count=452). Only one of the four positively-charged residues implicated in RNA binding underwent substitution (K2000R), which was conservative. Other amino acid changes involving positively-charged residues include both conservative (R1953K, K1973R, K1988R, K2004R, K2059R, and K2067R) and non-conservative substitutions (K1929T, K2029E/N, K2045Q, and K2059N) (Supplementary Figure nsp3e).

Supplementary Figure nsp3e. Ribbon/stick figure representation of the computed structural model of nsp3c. Lysine and Arginine sidechain atom color coding is as follows: invariant or conservative substituted residue-C purple, N-blue; non-conservative substituted residue-C purple, N-blue.

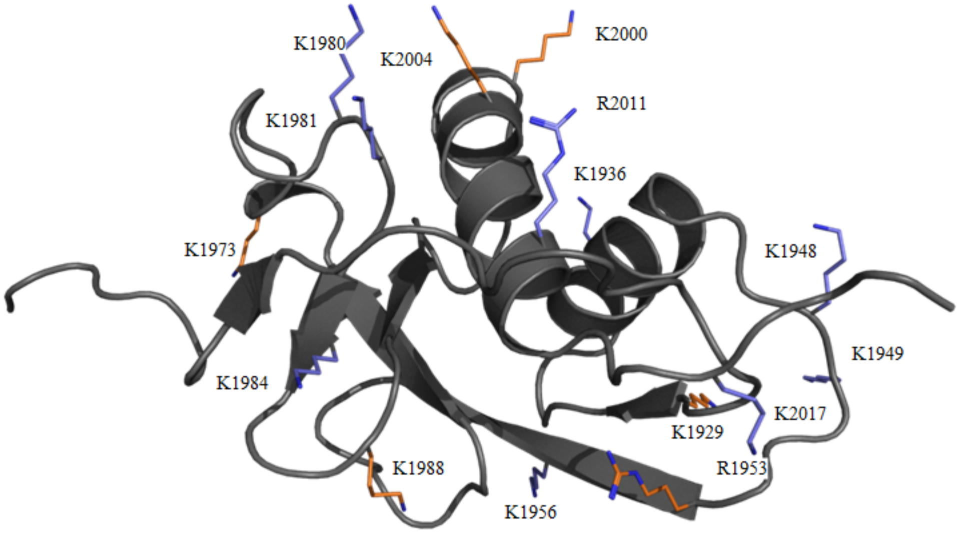

Nonstructural protein 3 Unknown (UNK): UNK, the C-terminal portion of nsp3, is 686 residues in length. It is ~80% identical to SARS-CoV-1 UNK, from which the functional description that follows was inferred. Binding of UNK to nsp4 is critical for formation of double-membrane vesicles (DMVs) derived from the endoplasmic reticulum (ER) in coronavirus-infected cells (Lei et al., 2018). It is 80% sequence identical to the corresponding segment of SARS-CoV-1 nsp3 (cite DOI: 10.1126/science.1085952). UNK contains two transmembrane segments (TM1, TM2), three soluble domains (3Ecto, Y1, CoV-Y), and one additional hydrophobic region (AH1). The order of these domains in the polypeptide chain is TM1—3Ecto—TM2—AH1—Y1—CoV-Y. UNK is believed to pass through the endoplasmic reticulum (ER) membrane twice, positioning the 3Ecto region on the lumenal side of the ER (Lei et al., 2018). The transmembrane segments plus the 3Ecto domain are critical for the nsp3-nsp4 cleavage site to be processed by PLPro. The 3Ecto domain undergoes N-glycosylation (Kanjanahaluethai, Chen, Jukneliene, & Baker, 2007). N-linked glycans often serve as recognition points for other molecules, which is notable because the 3Ecto domain interacts with nsp4 during rearrangements of the ER membrane in coronavirus infected cells. The Y1 and CoV-Y domains are located within the cytosol. Although functional information is somewhat limited, it is known that nsp3 binds less efficiently to nsp4 in the absence of the Y1 and CoV-Y domains ER (Lei et al., 2018). The UNK structural model used for analyzing evolution in 3D was computed by the David Baker Laboratory during a CASP competition (CASP-C1904 Stage 2, Supplementary Figure UNK). [N.B.: This structural model contains part of the βSM domain, which is sometimes considered to be part of nsp3e.]

Overall substitution trends for UNK and energetics analysis results are summarized in Tables 1 and 2. S2242F (non-conservative, surface) is the most common substitution (GISAID dataset count=128). The most intriguing amino acid change observed within UNK was N2272D. N2272 is one of two glycosylation sites occurring in the 3Ecto domain (Supplementary Figure UNK). Mutation to Aspartate would preclude glycosylation, possibly affecting the interaction between nsp3 and nsp4.

Supplementary Figure UNK. Ribbon/stick figure representation of the structural model for UNK. Color coding: βSM-purple, TM1-blue, 3Ecto domain-dark green, TM2-light green, AH1-orange, and the Y1 and CoV-Y domains-red. N2272, the site of glycosylation, is labeled and shown as a dark green stick figure.

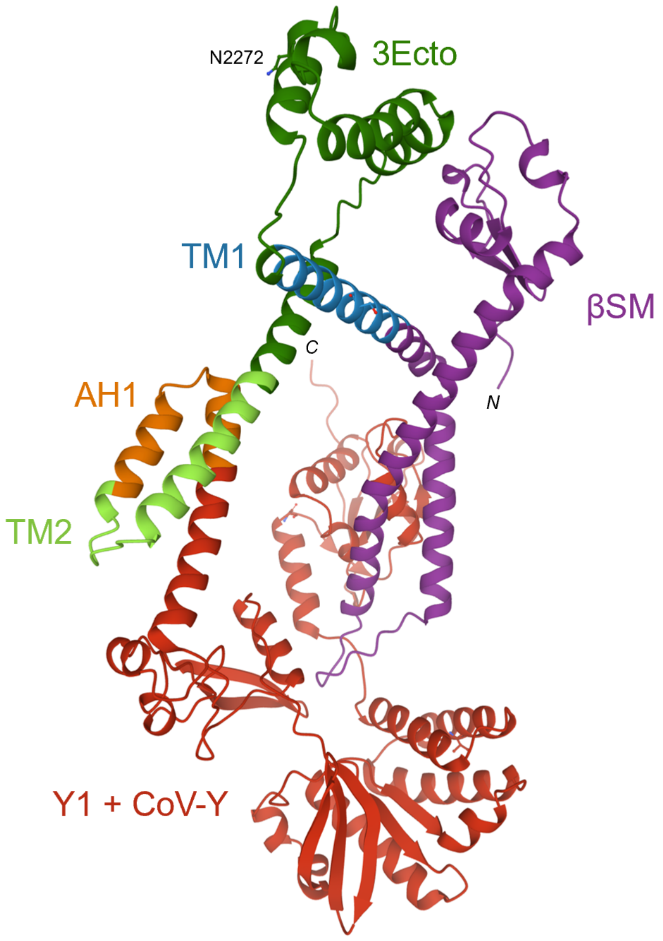

Non-structural protein 4 (nsp4): nsp4, 500 residues in length, is expressed as part of polyproteins 1a and 1ab and excised therefrom by nsp5. It is ~80% sequence identical to SARS-CoV-1 nsp4 (Yoshimoto, 2020), from which the functional description that follows was inferred. Nsp4 is an essential component of the replication-transcription complex that is responsible, together with nsp3 and nsp6, for inducing DMV and convoluted membrane formation in infected cells (Angelini, Akhlaghpour, Neuman, & Buchmeier, 2013). nsp4 is localized to the ER membrane where it is recruited into the replication complex in infected cells (Xu et al., 2009). It is predicted to contain four trans-membrane helices (Domains 1 and 3) with its N- and C-termini occurring the cytosol. Nsp4 interacts with nsp3 to induce membrane rearrangement. The nsp3 binding has been mapped to the C-terminal portion of nsp3 (UNK) and a lumenal segment of nsp4 occurring within Domain 2 (Sakai et al., 2017). This interaction is not sufficient for membrane rearrangement, suggesting that host proteins may be involved in this process (Sakai et al., 2017). The nsp4 structural model used for analyzing evolution in 3D was computed by the David Baker Laboratory during a CASP competition (CASP-C1902 Stage 2, Supplementary Figure nsp4).

Overall substitution trends for nsp4 and energetics analysis results are summarized in Tables 1 and 2. F308Y (conservative, boundary) is the most common substitution (GISAID dataset count=800). Two potentially important amino acid changes occurred together in luminal Domain 2 at residues 120 and 121 (H120Q, F121C; Supplementary Figure nsp4). These two conserved nsp4 residues are involved in inducing membrane rearrangements. The SARS-CoV-1 H120N/F121L double substitution displayed defects in membrane remodeling activity (Sakai et al., 2017). Although the mechanism of DMV formation is unknown, it is possible that the H120Q and F121C substitutions impact protein function of nsp4 during SARS-CoV-2 infection.

Supplementary Figure nsp4. Ribbon/ball-and-stick figure representation of the computed structural model for nsp4 color coded as follows: N-terminal transmembrane Domain 1-purple, luminal Domain 2-light blue, additional transmembrane regions Domain 3-orange, and C-terminal Domain 4-red. Four residues (L18, H120, F121, F308) are shown with Atom color coding as follows: C-blue or purple, N-blue, O-red). N- and C-termini are labeled.

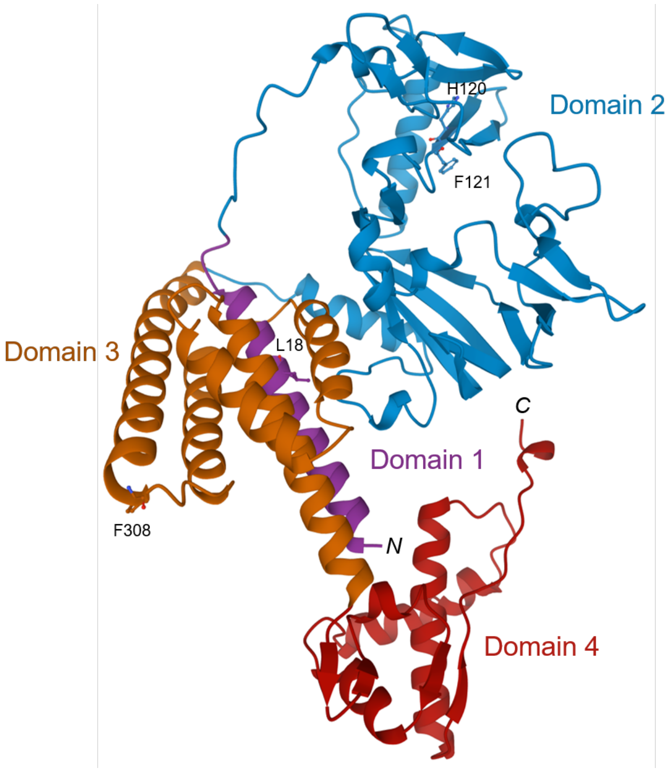

Non-structural protein 6 (nsp6): nsp6, a 290 amino acid multi-pass membrane protein, is expressed as part of polyproteins 1a and 1ab and excised therefrom by nsp5. It is ~88% sequence identical to SARS-CoV-1 nsp6, from which the functional description that follows was inferred. Nsp6 is thought to be responsible for restricting autophagosome expansion within the cell (Angelini et al., 2013). It also interferes with autophagosome delivery of viral factors to lysosomes for destruction. As shown for other coronaviruses (*e.g.*, SARS-CoV-1 and MHV), nsp6 gives rise to autophagosomes with smaller diameters (Cottam, Whelband, & Wileman, 2014). Restriction of autophagosome expansion may allow for further infection by limiting lysosomal degradation of viral components. Little structural information is available for any coronaviral nsp6 homologs. SARS-CoV-2 nsp6 is predicted to have 7 transmembrane helices with both termini occurring in the cytoplasm (Oostra et al., 2008). The computed structural model of nsp6 used to analyze evolution in 3D was generated by the David Baker Laboratory during the CASP-C1903 Stage 2 competition (Supplementary Figure nsp6).

Overall substitution trends for nsp6 and energetics analysis results are summarized in Tables 1 and 2. L37F (conservative, boundary) is the most common substitution (GISAID dataset count=5,375). L37F is also observed in 69 out 81 double substitution USVs. One notable USV is the sole occurrence of a double point substitution (K285N/A287S) in which the altered residues are separated by a single amino acid. Neither K285N nor A287S were observed in single occurrence USVs. A not dissimilar case is the USV with sole occurrence of three adjacent substitutions (S106A/G107C/F108C) None of S106A, G107C, or F108C substitutions were observed in single occurrence USVs. One or both of these cases could represent sequencing artifacts. An understanding of the potential impact of observed substitutions awaits further structural and functional characterization of nsp6.

Supplementary Figure nsp6. Ribbon representation of the computed structural model of nsp6, with N- and C- termini labeled.

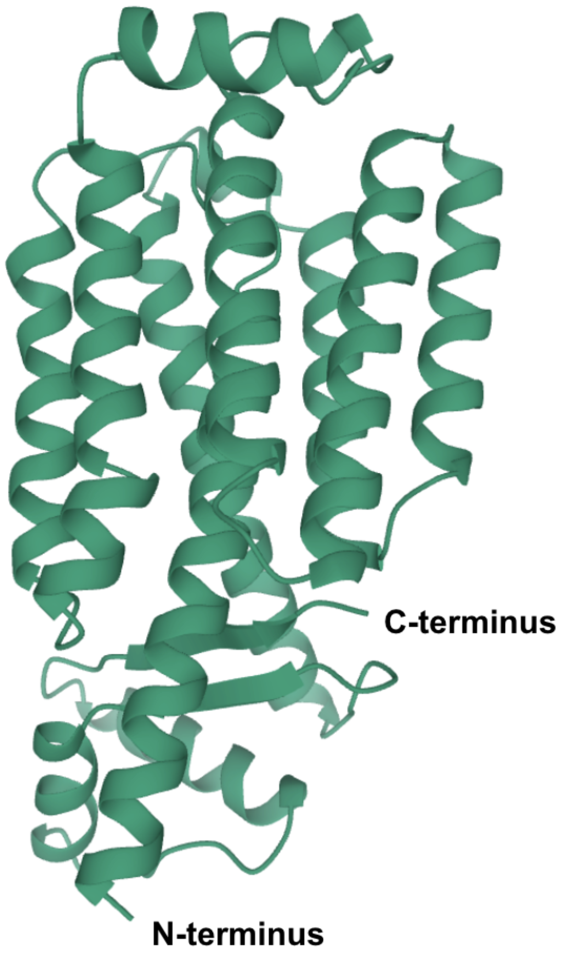

Non-structural protein 9 (nsp9): nsp9, 113 residues in length, is expressed as part of polyproteins 1a and 1ab and excised therefrom by nsp5. It is ~97% sequence identical to SARS-CoV-1 nsp9, which essential for viral replication (Frieman et al., 2012). The X-ray first structure of SARS-CoV-2 nsp9 (PDB ID 6W9Q (Littler, Gully, Colson, & Rossjohn, 2020)) revealed an $\alpha/\beta$ structure (Supplementary Figure nsp9) highly similar to that of its SARS-CoV-1 homolog (r.m.s.d.~0.6Å with PDB ID 1UW7 (Sutton et al., 2004)). Coronavirus nsp9 proteins are structurally similar to Greek key motif Ob-fold proteins known to bind nucleic acids (Theobald, Mitton-Fry, & Wuttke, 2003). Like its coronavirus homologs of known structure, SARS-CoV-2 nsp9 forms a symmetric homodimer stabilized by parallel approximation of two $\alpha$-helices presenting conserved GxxxG motifs to one another (100-GMVLG-104). Thirteen positively charged residues occurring on the molecular surface and in the boundary layer include R10, K36, R39, K52, R55, K58, R74, K81, K84, K86, K92, R99, and R111 (no Lysine or Arginine residues occur in the protein core). The experimental structure of SARS CoV-2 nsp9 (PDB ID 6W9Q (Littler et al., 2020)) was used for our evolutionary analyses. (Supplementary Figure nsp9)

Overall substitution trends for nsp9 and energetics analysis results are summarized in Tables 1 and 2. T77I (non-conservative, surface) is the most common substitution (GISAID dataset count=68). One of the two Glycine residues at the heart of the homodimer interface, G100, did not undergo amino acid changes during the pandemic. G104, on the other hand, was substituted with an Arginine residue (G104R) in a single USV with no other sequence changes that was found only once in the GISAID dataset (Supplementary Figure nsp9 A). The energetic consequences of this change are highly destabilizing ($\Delta\Delta G^{App}$=+20.4 REU). Experimental characterization of G104R substituted nsp9 could shed light on the plasticity of the nsp9 dimer interface and/or the importance of homodimer formation. Alternatively, it may reveal that the observed G104R substitution is a sequencing artifact.

Supplementary Figure nsp9 B highlights the locations of all positively charged amino acids occurring in the symmetric homodimer, which may be of importance for RNA binding. Five invariant Arginine and Lysine residues (purple) include R10, K84, K86, R99, and R111. The remaining eight positively charged amino acids underwent one or more changes (orange). Most of these substitutions were conservative: R36K, R55K, K58R, K81R, and K92R. Only three basic residues exhibited non-conservative changes: K36M, K36N, K52Q, and R74G. Visual inspection of the homodimer structure revealed a groove flanked by two Lysine (K81, K81', where ' denotes second protomer) and four Arginine (R10, R10',

R111, R111’) amino acids, all of which were either invariant (R10, R111) or underwent conservative substitution (K81R) during the pandemic. Preservation of six positively charged residues flanking the groove on the surface of the homodimer provides important clues as to the location of the RNA binding site, which could be verified experimentally.

Supplementary Figure nsp9. Ribbon/stick figure representations of the experimental structure of the nsp9 homodimer (PDB ID 6W9Q (Littler et al., 2020)) with individual protomers colored magenta and grey. (A) Dimerization interface viewed perpendicular to the two-fold axis of symmetry showing the locations of invariant G100 (purple) and substituted G104 (orange). (B) Putative RNA-binding groove flanked by six basic residues (R10, K81, and R111) viewed along the two-fold axis of symmetry. Atom color coding: C-purple(invariant), orange(substituted); N-blue; O-red.

A

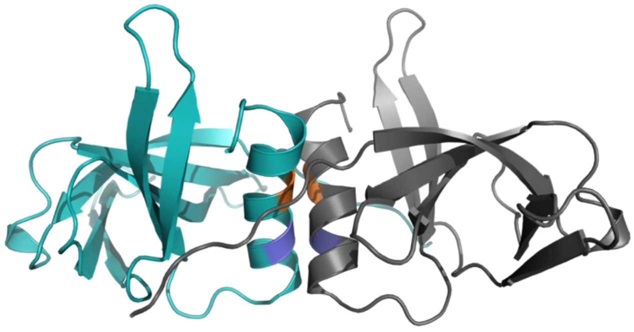

B

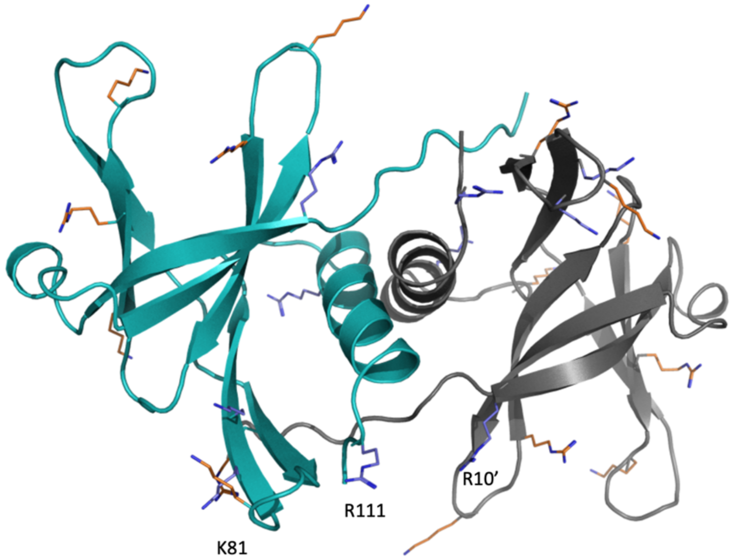

Non-structural protein 15 (nsp15): nsp15, 346 residues in length, is expressed as part of polyproteins 1a and 1ab and excised therefrom by nsp5. It is ~88% sequence identical to SARS-CoV-1 nsp15 (Y. Kim et al., 2020), from which much of the functional description below was derived. Nsp15 possesses a nidoviral uridylate-specific endoribonuclease (NendoU) catalytic site at its C-terminus (Y. Kim et al., 2020). The NendoU domain of nsp15 belongs to the EndoU family of RNA processing enzymes. While nsp15 is not essential for viral replication, conservation of NendoU activity amongst coronaviruses suggests that it must be biologically significant. Nsp15 assembles into a homohexamer made up of a dimer of trimers (Supplementary Figure nsp15 A). Active site residues include: H235, H250, K290, T341, Y343, and S294 (Supplementary Figure nsp15 B). H235, H250, and K290 form a catalytic triad within the active site. Y343, S294, and T341 are thought to participate in uracil binding (Bhardwaj et al., 2008). Studies of SARS-CoV-1 nsp15 concluded that residues 265, 267, 269-272, 278, 280-289, 291, and 292 participate in intersubunit interactions, interacting with residues 10-15, 17, 34, 36, 41-43, 62, 64, 89, 163-166, and 168-173 on adjacent protomers (Joseph et al., 2007). There is evidence that nsp15 might be responsible for degrading viral RNA to conceal it from host cells. Current PDB holdings include 15 different coronavirus nsp15 structures, including homologs from SARS-CoV-2 (7), MERS-CoV (1), Human Coronavirus 229E (2), 3 SARS-CoV-1 (3), and HMV 2. Two experimental structures of nsp15 (PDB IDs 6WLC and 6WXC (Youngchang Kim et al., 2020)) were used to analyze its evolution in 3D (Supplementary Figure nsp15).

Overall substitution trends for nsp15 and energetics analysis results are summarized in Tables 1 and 2. V321L (conservative, boundary) is the most common substitution (GISAID dataset count=216). Strikingly, every nsp15 active site residue underwent amino acid changes with H250 being the sole exception (Supplementary Figure nsp15 B). The most intriguing of the active site substitutions involved residue 235, part of the catalytic triad (H235Y; non-conservative, boundary). This change was observed 8 times, suggesting that it does not abrogate catalytic activity. K290N (non-conservative, boundary), another substitution to a catalytic residue, was observed 4 times and appears to remodel a hydrogen bond network in the active site. Other active site changes include H338Y, T341A/I, and Y343C. Among the nsp15 USVs, 74 included substitutions within 48 sites at protein-protein interfaces. Position 283 was substituted in five USVs, and D283G (non-conservative, boundary) was observed 13 times. A173L (conservative, boundary) was observed 110 times and was the most frequently occurring substitution. Experimental characterization of the active site substitutions could reveal interesting aspects of nsp15 function.

Supplementary Figure nsp15. (A) Space filling representation of the nsp15 hexamer with protomers colored in various shades of blue. (B) Ribbon/ball-and-stick figure representation of the enzyme active site showing H235, H250, K290, T341, Y343, and S294. Atom color coding: C-green, N-blue, O-red.

A

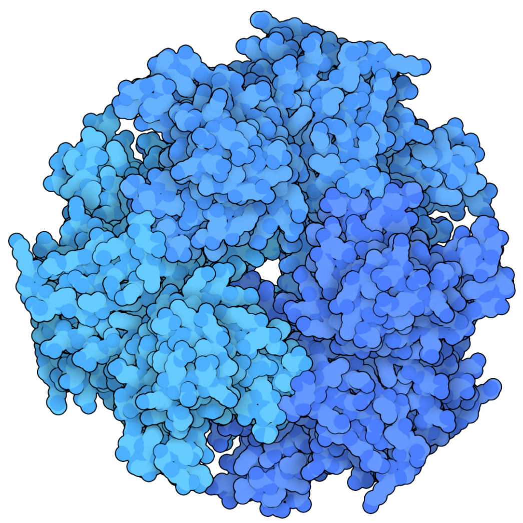

B

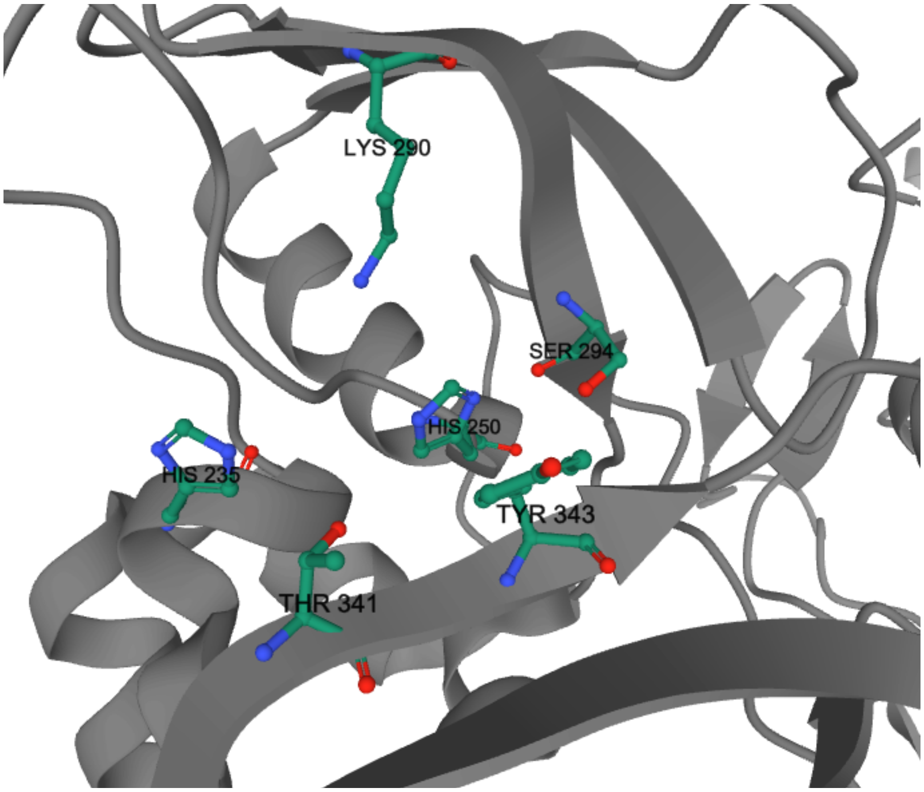

Open reading frame 3a (Orf3a): Orf3a, 275 residues in length expressed by subgenomic RNA 3, is an accessory protein that plays multiple roles during viral infection. SARS-CoV-2 Orf3a is ~79% identical in sequence to SARS-CoV-1 Orf3a, from which much of the functional description below is inferred (Issa, Merhi, Panossian, Salloum, & Tokajian, 2020). Orf3a interacts with the M-protein, the E-protein, and the S-protein to help form virus-like particles, playing a structural role in infection (Padhan et al., 2007). Orf3a interacts with the S-protein through disulfide bonds to assist in S-protein trafficking within the endoplasmic reticulum-Golgi intermediate complex. Finally, Orf3a functions as an ion channel, which has been implicated in viral particle formation and host cells apoptosis (Hachim et al., 2020; Kern et al., 2020). It has been found that membrane association of SARS-CoV-2 Orf3a but not SARS-CoV-1 Orf3a is necessary for apoptosis, leading to a weaker pro-apoptotic activity of the SARS-CoV-2 homolog (Ren et al., 2020). This property of SARS-CoV-2 may result in reduced apoptosis-mediated antiviral defense in infected cells. Reduced immune response may underpin the fact that many individuals affected by the COVID-19 pandemic are only mildly symptomatic (possibly even asymptomatic) in the initial stages of infection, thereby allowing the virus to spread more widely.

Orf3a consists of five functionally distinct segments, including an N-terminal domain, a C-terminal domain, and three transmembrane segments (Kern et al., 2020). The first 15 N-terminal residues serve as a signal sequence. The three transmembrane $\alpha$-helices contain pore forming residues. The C-terminal domain, comprising ~50% of the polypeptide chain, contains three sequence motifs (Cysteine-rich, Tyrosine-sorting, Glutamate-X-Aspartate or EXD diacidic motifs), all of which contribute to subcellular localization within the Golgi complex. A 3DEM structure of Orf3a (PDB ID 6XDC (Kern et al., 2020)) revealed a symmetric homodimer (Supplementary Figure Orf3a). Segments of the polypeptide chain unresolved in the 3DEM structure include residues 1-39, 175-180, and 239-275. At the time of writing, there were no other coronavirus Orf3a structures and no structurally similar proteins in the PDB. Pore-forming residues identified in PDB ID 6XDC are as follows: F43, L46, I47, V50, L53, A54, Q57, S60, K61, H78, C81, L85, V88. The experimental structure of Orf3a (PDB ID 6XDC (Kern et al., 2020)) was used to analyze its evolution in 3D (Supplementary Figure Orf3a).

Overall substitution trends for Orf3a and energetics analysis results are summarized in Tables 1 and 2. Orf3a was the most prolifically substituted SARS-CoV-2 protein in the GISAID dataset (2.46 USVs/protein residue). Q57H (non-conservative, boundary) is the most common substitution (GISAID dataset count=10,593), and G251V is the second most common (GISAID dataset count=3,867). Q57H and G251V substitutions also occur together in some USVs. The Q57H substitution has no significant impact on protein function (Kern et al., 2020). The impact of the G251V substitution has not been assessed experimentally. A total of 58 USVs contain substitutions map to the transmembrane segments (Supplementary Figure Orf3a). Mutations of residues lining the pore are as follows: F43I/L/S/Y, L46F, I47V, V50A/I, L53F/H, A54S/T/V, Q57H/R, S60F, K61N, H78R/Y, C81F/Y, L85F, and V88A/L. Cysteines occurring in the Cysteine-rich motif are thought to contribute to homodimerization (Issa et al., 2020). Observed Cysteine substitutions included C81F/Y, C13S, C148F/Y, C153Y, and C200Y, of which residues 81, 148, and 153 were observed to undergo substitution in more than one USV. Potentially important changes observed in this region involve W131, which was observed in 13 USVs, substituted for C, L, R, or S.

Supplementary Figure Orf3a. Ribbon/ball-and-stick figure representations of the experimental structure of the Orf3a homodimer (PDB ID 6XDC (Kern et al., 2020)) with protomers colored orange and green. (A) Orthogonal views showing frequently observed substitutions. (B) View showing pore-lining residues. Atom color coding: C-green or orange, N-blue, O-red, S-yellow.

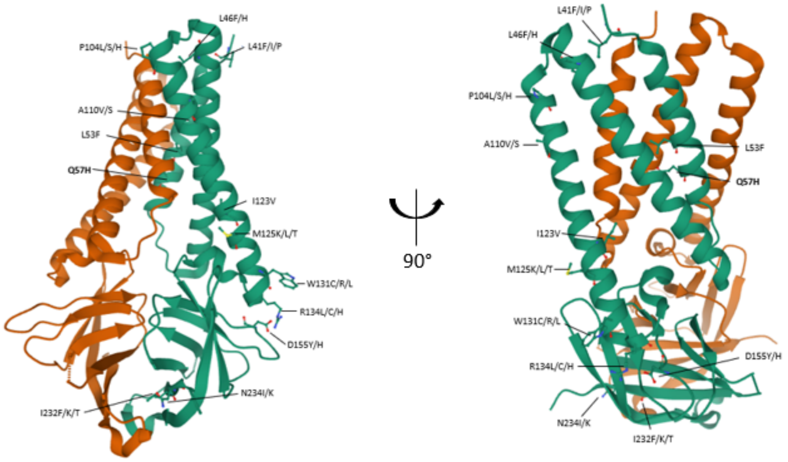


B

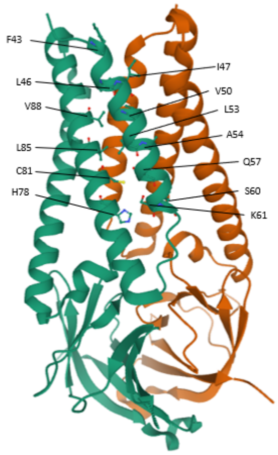

Open reading frame 6 (Orf6): Orf6, 61 residues in length, is expressed by subgenomic RNA 6. SARS-CoV-2 Orf6 is ~69% sequence identical to SARS-CoV-1 Orf6 (Cagliani, Forni, Clerici, & Sironi, 2020), from which much of the functional description below is inferred. Orf6 acts as an interferon antagonist that prevents interferon production and signaling (particularly for β interferon), aiding SARS-CoV-2 in evading the host immune response. In SARS-CoV-1 and SARS-CoV-2 Orf6 potently suppress primary interferon production and interferon signaling (Miorin et al., 2020; Narayanan, Huang, & Makino, 2008; Yuen et al., 2020). The computed structural model of Orf6 used to analyze evolution in 3D was generated by the David Baker Laboratory during the CASP-C1907 Stage 1 competition (Supplementary Figure Orf6).

Overall substitution trends for Orf6 and energetics analysis results are summarized in Tables 1 and 2 (Supplementary Figure Orf6). I33T was the most common observed substitution (GISAID dataset count=177). The most intriguing amino changes observed map to C-terminal residues (D53G/Y, E54D, E55D/Q, Q56H/R, P57L, M58I/L/T) which play a key role in Orf6 binding to the Nup98-Rae1 complex at the nuclear pore to disrupt nuclear import leading to impaired interferon signaling (Miorin et al., 2020). Finally, substitutions to or from Proline residues (S50P, P57L, both observed alone in singleton USVs) within the Orf6 predicted α-helix structural model may perturb structure and/or function.

Supplementary Figure Orf6. Ribbon representation of the computed structural model for Orf6 showing labeled N- and C-termini.

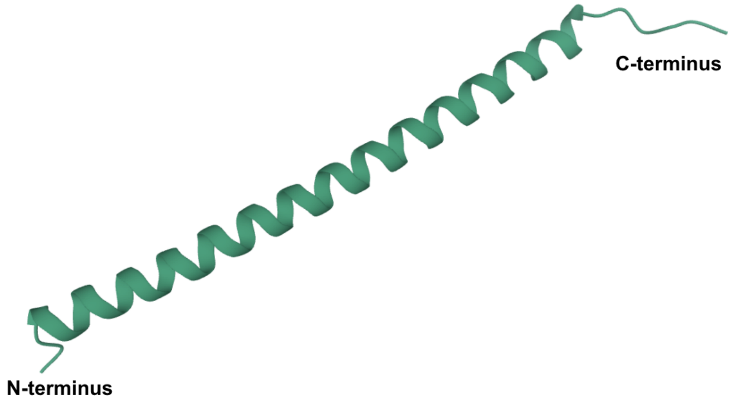

Open reading frame 7a (Orf7a): Orf7a, 121 residues in length expressed by subgenomic RNA 7, is a type I transmembrane protein that localizes to Golgi bodies and the endoplasmic reticulum (C. A. Nelson, Pekosz, Lee, Diamond, & Fremont, 2005). SARS-CoV-2 Orf7a ~85% sequence identical to SARS-CoV-1 Orf7a, from which much of the functional description below was inferred. Although the function of Orf7a in the SARS-CoV-2 infection cycle is not yet established, we know that its SARS-CoV-1 homolog is responsible for binding BST-2 (tetherin), a host restriction factor that prevents new viruses from leaving the cell. Orf7a disrupts N-linked glycosylation of BST-2, resulting in increased SARS-CoV-1 virion release (Yuan et al., 2006). In addition, SARS-CoV-1 Orf7a transfected cells undergo cell cycle arrest at the G0/G1 checkpoint. Cell-cycle arrest appears to be mediated through the cyclin D3/pRb pathway, possibly due to inhibition of host cell mRNA translation. Arrested cell cycle progression may allow time for SARS-CoV-1 to replicate at higher rates, with increased nucleotide pools, and promote viral pathogenicity (Yuan et al., 2006). Interactions between SARS-CoV-1 Orf7a and human $Ap_4A$-hydrolase has documented. $Ap_4A$-hydrolase is thought to be in DNA replication and repair, RNA processing and apoptosis. Coronavirus Orf7a homologs may processes relating to cell cycle progression and apoptosis (Vasilenko, Moshynskyy, & Zakhartchouk, 2010). The experimental structure of SARS-CoV-2 Orf7a (PDB ID 6W37 (C.A. Nelson, Minasov, Shuvalova, & Fremont, 2020)) was used to analyze its evolution in 3D (Supplementary Figure Orf7a).

Overall substitution trends for Orf7a and energetics analysis results are summarized in Tables 1 and 2 (Supplementary Figure Orf7a). S81L (non-conservative, boundary), the most frequently observed substitution (GISAID dataset count=130), occurs in between the β-sheet bundle and stalk α-helix adjacent to the transmembrane domain. G26D/R (non-conserved, boundary), located on the loop between the first and second beta sheet bundle, may affect loop flexibility due to the larger bulk of arginine. V104F (conservative, boundary), located on the C-terminal stalk α-helix, is another variant of interest, as it appears as both a single and a double mutant substitution (V104F/T120I). Currently, little is known regarding the binding interface with BST-2, limiting further insights into the structural and functional significance of these substitutions. While we did not identify any insertions or deletions in the GISAID dataset, there has been a report of an observed 27-amino acid deletion of a portion of Orf7a (putative signal peptide and first two beta strands) in a sample isolated in Arizona, USA (Holland et al., 2020). The impact of this deletion on Orf7a and the resulting virus properties remains to be investigated.

Supplementary Figure Orf7a. Ribbon representation of the experimental structure of Orf7a (PDB ID 6W37 (C.A. Nelson et al., 2020)) with N- and C-termini labeled.

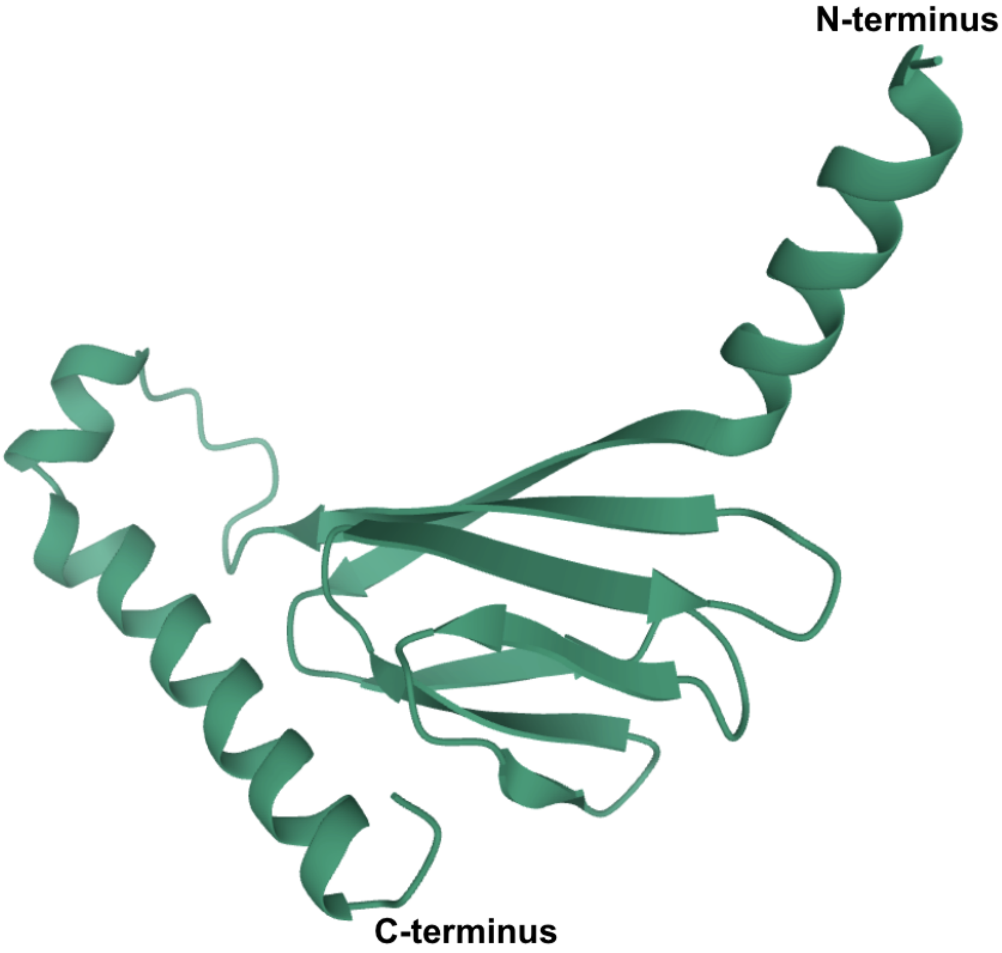

Open Reading Frame 7b (Orf7b): Orf7b, 43 residues in length expressed by subgenomic RNA 8, consists of three segments: N-terminal (residues 1-8), C-terminal (residues 31-43), and the transmembrane domain (TMD, residues 9-30). SARS-CoV-2 Orf7b is ~80% sequence identical to SARS-CoV-1 Orf7b, from which much of the functional description below has been inferred. There are no homologs of Orf7b in more distantly related coronaviruses (*e.g.*, MERS, MHV) (Schaecher, Mackenzie, & Pekosz, 2007). The TMD localizes in the Golgi membrane (Schaecher, Diamond, & Pekosz, 2008). The N-terminal and C-terminal segments occur in the Golgi lumen and the cytosol, respectively, and do not contribute to subcellular localization. Orf7b is classified as an accessory protein but is also thought to play a structural role in SARS-CoV-1 virions. It is believed to be incorporated into SARS-CoV-1particles due to the proximity of the protein to the viral budding site of the Golgi complex (Schaecher et al., 2007). The computed structural model of Orf7b used for analyzing evolution in 3D was generated by the David Baker Laboratory during a CASP competition (CASP-C1910 Stage 1) (Supplementary Figure Orf7b).

Overall substitution trends for Orf7b and energetics analysis results are summarized in Tables 1 and 2 (Supplementary Figure Orf7b). The most commonly observed substitution was C41F (GISAID dataset count=74). Nearly half of all substitutions 44% (25/57 variants) occurred within the TMD. USVs of potential interest include L18P, L25P, and a double substitution (L32H/E33P), all of which result in the introduction of a helix-breaking Proline residue within or close to the $\alpha$-helical TMD (residues 9-30). There are seven Orf7b residues, located in the TMD, that are thought to play a role in Golgi complex retention (residues 13-15, 19-22). A total of 9 USVs include substitutions at these positions. Residues 14, 21, and 22 were unchanged, and in particular, 14 and 21 are positioned on the safe face of the $\alpha$-helicx. Experimental characterization of these substitutions could help further elucidate the role(s) played by Orf7b during viral infection.

Supplementary Figure Orf7b. Ribbon representation of the computed structural model of Orf7b with labeled N- and C-termini, showing the site of the most frequently changed residue C41 (Atom color coding: C-green, N-blue, O-red, and S-yellow).

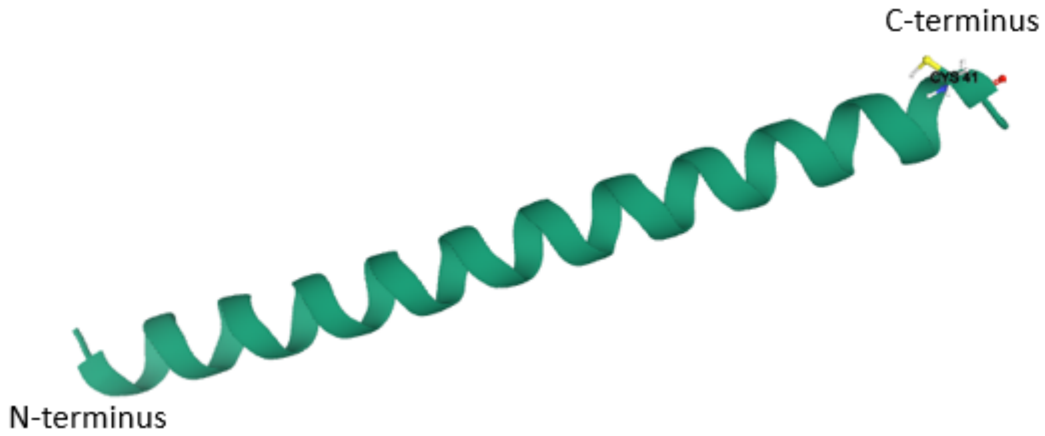

Open reading frame 8 (Orf8): Orf8, 121 residues in length expressed by subgenomic RNA 8, includes an N-terminal signal sequence that directs it to the ER (Mohammad et al., 2020). While the function of Orf8 is not yet established, it was hypothesized that it binds major histocompatibility-I (MHC-I) molecules, targeting them for lysosomal degradation (Mohammad et al., 2020). Reduction of MHC-I density on the cell surface is thought to reduce cytotoxic T lymphocyte surveillance and killing of infected cells (Zhang et al., 2020). Unlike most SARS-CoV-2 study proteins, Orf8 is <20% identical in sequence to SARS-CoV-1 and other related coronavirus homologs, making it the most divergent of the coronavirus proteins (Flower et al., 2020). Orf8 also appears to be secreted from the ER. Antibodies to Orf8 constitute serological markers of SARS-CoV-2 infection (Hachim et al., 2020). The computed structural model of Orf8 used to analyze its evolution in 3D was generated by the David Baker Laboratory during the CASP-C1908 Stage 2 competition (Supplementary Figure Orf8).

Overall substitution trends for Orf8 and energetics analysis results are summarized in Tables 1 and 2. L84S (non-conservative, surface) is the most common substitution, observed 3,847 times. L84S was also observed in 30 multi-point mutations. A number of cysteine residues that form intra-molecular disulfide bonds are found to be substituted, including C25F, C37F, C83F, and C102F. These substitutions do not occur concurrently in USVs with substitutions gaining cysteine. The variants may also have altered structural and/or folding properties and may be worthy of further investigation. While no deletions were observed in the dataset we collected, Orf8 deletions appearing in patients in Singapore have been reported (Su et al., 2020), indicating that Orf8 function may not be critical for viral replication and transmission.

Orf8 was modeled by the David Baker laboratory as a monomer with an IgG-like fold (Supplementary Figure Orf8). A dimeric crystal structure of Orf8 was reported after our analyses were concluded (PDB ID 7JTL (Flower et al., 2020)). As both monomeric and dimeric species are present during size exclusion chromatography (Flower et al., 2020), it is unclear if the dimeric species observed in the crystal structure is a crystallization artifact. Comparison of the computed structural model of the monomer and the dimeric X-ray crystal structure reveals that while the overall folds are similar, β-strand pairings do differ. The dimer includes two interfaces: a covalent disulfide-linked dimer is formed *via* residue C20 found in SARS-CoV-2 (but not SARS-CoV-1), and a separate non-covalent interface is formed by another SARS-CoV-2-specific sequence (73-YIDI-76). However, the possibility of dimeric linkage at C20 suggests the observed C20F substitution merits further investigation, as it would abolish the disulfide bond and potentially cause further structural perturbation(s). C20F was observed in two USVs. Substitutions were identified for all residues occurring within the second protein-protein interface, including Y73H, I74F, D75G/N/V, and I76F/V.

Supplementary Figure Orf8. Ribbon/stick figure representation of the computed structural model of Orf8 with labeled N- and C-termini, showing R48, V49, and L84–the most commonly substituted residue (Atom color coding: C-green, N-blue, O-red, and S-yellow).

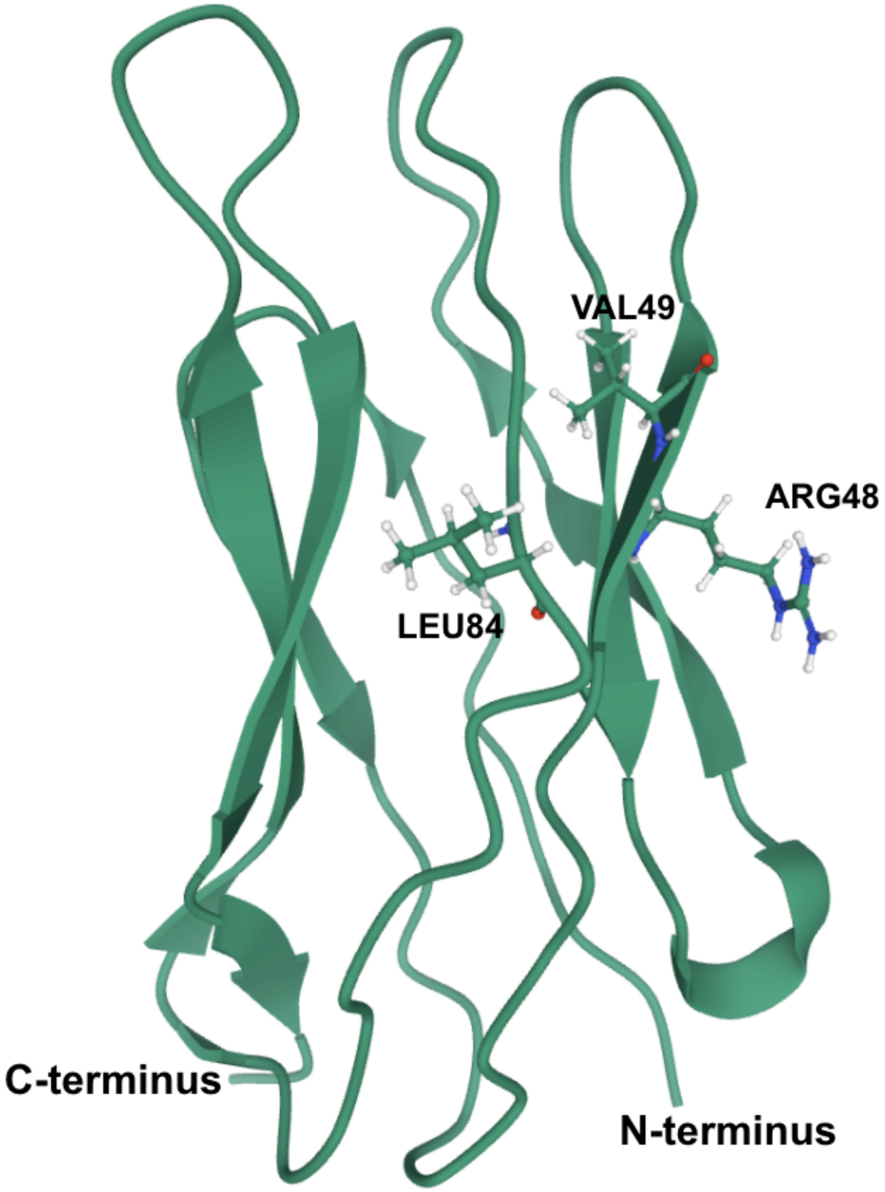

2. Legends for Supplementary Figures for 29 SARS-CoV-2 Study Proteins

**Separate analysis of protein changes was performed for each study protein and complex. Description below applies to all figures.**

A: Grey scale representation of observed frequencies for all USV substitutions of Native Residue (i.e., amino acid type in the reference protein sequence) changing to Substituted Residue for a given protein/complex. Red boxes enclose conservative substitutions for hydrophobic, uncharged polar, positively charged, and negatively charged amino acids, respectively in order from upper left to lower right. Cysteine, Glycine and Proline are excluded from these groupings.

B-D: Normalized Frequency histograms for $\Delta\Delta G^{App}$ calculated for all USVs for a given protein/complex. These were calculated using three methods, which we refer to as hard-hard (B), soft-hard (C), and soft-soft (D), based on the scoring functions used for sidechain rotamer optimization and gradient-based energy minimization respectively (see methods). All energy values described in the text were obtained using the soft-hard method. Overlay of energy histogram with fitted bi-Gaussian curve (solid red line) and fitted single Gaussian curves for subsets of USVs with surface (green), boundary layer (yellow), or core (blue) substitutions. USVs with multiple substitutions were included in single Gaussian fitting when all substitutions mapped to the same region of the study protein. The data used for fitting includes the energies of all unique protein models produced by a given method, excluding extreme outliers with energy values greater than four standard deviations away from the central mean.

E-G: USV Count histograms indicate the number of USVs among the full set for a given protein in which each site included a substitution. Sites are separated by burial layer. Substitutions at sites that are absent from the available crystal structures are excluded from the histograms. In most cases, only a single protein is analyzed, and only panel E is included. In the case of complexes, a separate histogram is provided for each protein in the complex: for methyltransferase Nsp10-Nsp16, E is Nsp10 and F is Nsp16; for RDRP Nsp12-Nsp7-Nsp8, E is Nsp7, F is Nsp8, and G is Nsp12.

### 3. Legends for Supplementary Tables for 29 SARS-CoV-2 Study Proteins

**Table USVs**: All identified USVs for a protein/complex. Columns are:

- date: Date of first collection of a strain with the USV reported to GISAID
- gisaid_count: The number of sequences in the GISAID database that include the USV
- id: The GISAID strain identification for the first collected instance of the USV
- location: The country in which the first strain including the USV was collected
- substitutions: All substitutions in the USV, in the form [chain]_[sequence][site][substitution], with multiple substitutions separated by semicolons
- is_in_PDB: whether a substitution is present in the PDB model used to generate the USV structure, with multiple substitutions separated by semicolons
- multiple: whether more than one amino acid substitution is present in the USV
- conservative: whether a substitution is conservative, with multiple substitutions separated by semicolons
- layer: Identification of the burial layer (surface, boundary, or core) of a substitution in the reference structure, with multiple substitutions separated by semicolons and substitutions absent from the PDB excluded
- sh_rmsd: The RMSD of the USV to the reference structure when modeled using the soft-hard method
- sh_ddG: The $\Delta\Delta G^{App}$ of the USV when modeled using the soft-hard method
- hh_rmsd: The RMSD of the USV to the reference structure when modeled using the hard-hard method
- hh_ddG: The $\Delta\Delta G^{App}$ of the USV when modeled using the hard-hard method
- ss_rmsd: The RMSD of the USV to the reference structure when modeled using the soft-soft method
- ss_ddG: The $\Delta\Delta G^{App}$ of the USV when modeled using the soft-soft method

**Table Substitutions**: All substitutions identified for a protein/complex

- chain: The chain identifier of the protein in the PDB file in which the substitution is present
- site: The residue number at which the substitution is present
- reference: The one-letter amino acid name of the residue in the reference sequence
- mutant: The one-letter amino acid name of the residue in a USV
- conservative: Indication of whether a substitution is conservative
- in_pdb: whether the substitution site is present in the PDB model used to generate the USV structure
- layer: Identification of the burial layer (surface, boundary, or core) of a substitution in the reference structure
- date: date: Date of first collection of a strain with the substitution reported to GISAID
- location: The country in which the first strain including the substitution was collected
- gisaid_count: The number of sequences in the GISAID database including the substitution
- usv_count: The number of identified USVs including the substitution
- ddG: The soft-hard $\Delta\Delta G^{App}$ of the USV that includes only the substitution, left empty if no single-substitution USV was identified with the substitution
- single: Indication of whether the substitution was present in a single-substitution USV
- multiple: Indication of whether the substitution was present in a USV with multiple substitutions
- associates: List of all other substitutions that were identified in a USV that included the substitution
- strains: List of all USV-representative GISAID strains that included the substitution, with the single-substitution USV strain listed first if one was available

**Table Gaussian Fit Statistics**: Fitted models for the energies of all USVs either together (ALL) or by study protein.

- fit: The number of Gaussian curves in the fitted energy model
- protein: The protein/complex name
- method: The modeling method used to calculate energy values
- layer: The subset burial layer (surface, boundary, or core) of USVs for which the energy model was fitted, excluding all USVs with substitutions not in that layer
- $\mu_1$: Mean of the first Gaussian in the fitted model
- $\sigma_1$: Variance of the first Gaussian in the fitted model
- $wt_1$: Weight of the first Gaussian in the fitted model
- $\mu_2$: Mean of the second Gaussian in the fitted model
- $\sigma_2$: Variance of the second Gaussian in the fitted model
- $wt_2$: Weight of the second Gaussian in the fitted model
- $R^2$: R-squared value indicating the goodness of fit

### 4. Description of Computed Structural Models for Unique Sequence Variants for 29 SARS-CoV-2 Study Proteins.

**USV Computed Structural Models.** Computed structural models for all amino acid substituted USVs. We are providing the structural models of all study proteins modeled using the soft-hard modeling method (see Methods). Structural models are named according to the GISAID strain identification of the first strain in which the USV was identified, followed by an underscore-separated list of substitutions in the form [chain]_[sequence][site][substitution]. Atomic coordinates for each computed structural model are provided in the legacy Protein Data Bank format used by most molecular graphics software tools (see https://www.wwpdb.org/documentation/file-format-content/format33/v3.3.html for detailed description).

### 5. Supplementary Materials References


Alhammad, Y. M. O., Kashipathy, M. M., Roy, A., Gagne, J. P., McDonald, P., Gao, P., . . . Fehr, A. R. (2020). The SARS-CoV-2 conserved macrodomain is a mono-ADP-ribosylhydrolase. *J Virol*. doi:10.1128/JVI.01969-20

Almeida, M. S., Johnson, M. A., Herrmann, T., Geralt, M., & Wuthrich, K. (2007). Novel beta-barrel fold in the nuclear magnetic resonance structure of the replicase nonstructural protein 1 from the severe acute respiratory syndrome coronavirus. *J Virol, 81*(7), 3151-3161. doi:10.1128/JVI.01939-06

Angeletti, S., Benvenuto, D., Bianchi, M., Giovanetti, M., Pascarella, S., & Ciccozzi, M. (2020). COVID-2019: The role of the nsp2 and nsp3 in its pathogenesis. *J Med Virol, 92*(6), 584-588. doi:10.1002/jmv.25719

Angelini, M. M., Akhlaghpour, M., Neuman, B. W., & Buchmeier, M. J. (2013). Severe acute respiratory syndrome coronavirus nonstructural proteins 3, 4, and 6 induce double-membrane vesicles. *MBio, 4*(4). doi:10.1128/mBio.00524-13

Bhardwaj, K., Palaninathan, S., Alcantara, J. M., Yi, L. L., Guarino, L., Sacchettini, J. C., & Kao, C. C. (2008). Structural and functional analyses of the severe acute respiratory syndrome coronavirus endoribonuclease Nsp15. *J Biol Chem, 283*(6), 3655-3664. doi:10.1074/jbc.M708375200

Cagliani, R., Forni, D., Clerici, M., & Sironi, M. (2020). Coding potential and sequence conservation of SARS-CoV-2 and related animal viruses. *Infect Genet Evol, 83*, 104353. doi:10.1016/j.meegid.2020.104353

Cottam, E. M., Whelband, M. C., & Wileman, T. (2014). Coronavirus NSP6 restricts autophagosome expansion. *Autophagy, 10*(8), 1426-1441. doi:10.4161/auto.29309

Fehr, A. R., Channappanavar, R., Jankevicius, G., Fett, C., Zhao, J., Athmer, J., . . . Perlman, S. (2016). The Conserved Coronavirus Macrodomain Promotes Virulence and Suppresses the Innate Immune Response during Severe Acute Respiratory Syndrome Coronavirus Infection. *MBio, 7*(6). doi:10.1128/mBio.01721-16

Flower, T. G., Buffalo, C. Z., Hooy, R. M., Allaire, M., Ren, X., & Hurley, J. H. (2020). Structure of SARS-CoV-2 ORF8, a rapidly evolving coronavirus protein implicated in immune evasion. *bioRxiv*. doi:10.1101/2020.08.27.270637

Frick, D. N., Virdi, R. S., Vuksanovic, N., Dahal, N., & Silvaggi, N. R. (2020). Molecular Basis for ADP-Ribose Binding to the Mac1 Domain of SARS-CoV-2 nsp3. *Biochemistry, 59*(28), 2608-2615. doi:10.1021/acs.biochem.0c00309

Frieman, M., Yount, B., Agnihothram, S., Page, C., Donaldson, E., Roberts, A., . . . Baric, R. S. (2012). Molecular determinants of severe acute respiratory syndrome coronavirus pathogenesis and virulence in young and aged mouse models of human disease. *J Virol, 86*(2), 884-897. doi:10.1128/JVI.05957-11

Graham, R. L., Sims, A. C., Brockway, S. M., Baric, R. S., & Denison, M. R. (2005). The nsp2 replicase proteins of murine hepatitis virus and severe acute respiratory syndrome coronavirus are dispensable for viral replication. *J Virol, 79*(21), 13399-13411. doi:10.1128/JVI.79.21.13399-13411.2005

Grunewald, M. E., Chen, Y., Kuny, C., Maejima, T., Lease, R., Ferraris, D., . . . Fehr, A. R. (2019). The coronavirus macrodomain is required to prevent PARP-mediated inhibition of virus replication and enhancement of IFN expression. *PLoS Pathog, 15*(5), e1007756. doi:10.1371/journal.ppat.1007756

Hachim, A., Kavian, N., Cohen, C. A., Chin, A. W. H., Chu, D. K. W., Mok, C. K. P., . . . Valkenburg, S. A. (2020). ORF8 and ORF3b antibodies are accurate serological markers of early and late SARS-CoV-2 infection. *Nat Immunol, 21*(10), 1293-1301. doi:10.1038/s41590-020-0773-7

Holland, L. A., Kaelin, E. A., Maqsood, R., Estifanos, B., Wu, L. I., Varsani, A., . . . Lim, E. S. (2020). An 81-Nucleotide Deletion in SARS-CoV-2 ORF7a Identified from Sentinel Surveillance in Arizona (January to March 2020). *J Virol, 94*(14). doi:10.1128/JVI.00711-20

Huang, C., Lokugamage, K. G., Rozovics, J. M., Narayanan, K., Semler, B. L., & Makino, S. (2011). SARS coronavirus nsp1 protein induces template-dependent endonucleolytic cleavage of mRNAs: viral mRNAs are resistant to nsp1-induced RNA cleavage. *PLoS Pathog, 7*(12), e1002433. doi:10.1371/journal.ppat.1002433

Issa, E., Merhi, G., Panossian, B., Salloum, T., & Tokajian, S. (2020). SARS-CoV-2 and ORF3a: Nonsynonymous Mutations, Functional Domains, and Viral Pathogenesis. *mSystems, 5*(3). doi:10.1128/mSystems.00266-20

Jauregui, A. R., Savalia, D., Lowry, V. K., Farrell, C. M., & Wathelet, M. G. (2013). Identification of residues of SARS-CoV nsp1 that differentially affect inhibition of gene expression and antiviral signaling. *PLoS ONE, 8*(4), e62416. doi:10.1371/journal.pone.0062416

Joseph, J. S., Saikatendu, K. S., Subramanian, V., Neuman, B. W., Buchmeier, M. J., Stevens, R. C., & Kuhn, P. (2007). Crystal structure of a monomeric form of severe acute respiratory syndrome coronavirus endonuclease nsp15 suggests a role for hexamerization as an allosteric switch. *J Virol, 81*(12), 6700-6708. doi:10.1128/JVI.02817-06

Kamitani, W., Huang, C., Narayanan, K., Lokugamage, K. G., & Makino, S. (2009). A two-pronged strategy to suppress host protein synthesis by SARS coronavirus Nsp1 protein. *Nat Struct Mol Biol, 16*(11), 1134-1140. doi:10.1038/nsmb.1680

Kanjanahaluethai, A., Chen, Z., Jukneliene, D., & Baker, S. C. (2007). Membrane topology of murine coronavirus replicase nonstructural protein 3. *Virology, 361*(2), 391-401. doi:10.1016/j.virol.2006.12.009

Keane, S. C., & Giedroc, D. P. (2013). Solution structure of mouse hepatitis virus (MHV) nsp3a and determinants of the interaction with MHV nucleocapsid (N) protein. *J Virol, 87*(6), 3502-3515. doi:10.1128/JVI.03112-12

Kern, D. M., Sorum, B., Hoel, C. M., Sridharan, S., Remis, J. P., Toso, D. B., & Brohawn, S. G. (2020). Cryo-EM structure of the SARS-CoV-2 3a ion channel in lipid nanodiscs. *bioRxiv*. doi:10.1101/2020.06.17.156554

Kim, Y., Jedrzejczak, R., Maltseva, N. I., Wilamowski, M., Endres, M., Godzik, A., . . . Joachimiak, A. (2020). Crystal structure of Nsp15 endoribonuclease NendoU from SARS-CoV-2. *Protein Sci, 29*(7), 1596-1605. doi:10.1002/pro.3873

Kim, Y., Wower, J., Maltseva, N., Chang, C., Jedrzejczak, R., Wilamowski, M., . . . Joachimiak, A. (2020). Tipiracil binds to uridine site and inhibits Nsp15 endoribonuclease NendoU from SARS-CoV-2. *bioRxiv*, 2020.2006.2026.173872. doi:10.1101/2020.06.26.173872

Lei, J., Kusov, Y., & Hilgenfeld, R. (2018). Nsp3 of coronaviruses: Structures and functions of a large multi-domain protein. *Antiviral Res, 149*, 58-74. doi:10.1016/j.antiviral.2017.11.001

Littler, D. R., Gully, B. S., Colson, R. N., & Rossjohn, J. (2020). Crystal Structure of the SARS-CoV-2 Non-structural Protein 9, Nsp9. *iScience, 23*(7), 101258. doi:10.1016/j.isci.2020.101258

Lokugamage, K. G., Narayanan, K., Huang, C., & Makino, S. (2012). Severe acute respiratory syndrome coronavirus protein nsp1 is a novel eukaryotic translation inhibitor that represses multiple steps of translation initiation. *J Virol, 86*(24), 13598-13608. doi:10.1128/JVI.01958-12

Miorin, L., Kehrer, T., Sanchez-Aparicio, M. T., Zhang, K., Cohen, P., Patel, R. S., . . . Garcia-Sastre, A. (2020). SARS-CoV-2 Orf6 hijacks Nup98 to block STAT nuclear import and antagonize interferon signaling. *Proc Natl Acad Sci U S A, 117*(45), 28344-28354. doi:10.1073/pnas.2016650117

Mohammad, S., Bouchama, A., Mohammad Alharbi, B., Rashid, M., Saleem Khatlani, T., Gaber, N. S., & Malik, S. S. (2020). SARS-CoV-2 ORF8 and SARS-CoV ORF8ab: Genomic Divergence and Functional Convergence. *Pathogens, 9*(9). doi:10.3390/pathogens9090677

Narayanan, K., Huang, C., & Makino, S. (2008). SARS coronavirus accessory proteins. *Virus Res, 133*(1), 113-121. doi:10.1016/j.virusres.2007.10.009
Nelson, C. A., Minasov, G., Shuvalova, L., & Fremont, D. H. (2020). Structure of the SARS-CoV-2 ORF7a encoded accessory protein. *10.2210/pdb6W37/pdb*.
Nelson, C. A., Pekosz, A., Lee, C. A., Diamond, M. S., & Fremont, D. H. (2005). Structure and intracellular targeting of the SARS-coronavirus Orf7a accessory protein. *Structure, 13*(1), 75-85. doi:10.1016/j.str.2004.10.010
Oostra, M., Hagemeijer, M. C., van Gent, M., Bekker, C. P., te Lintelo, E. G., Rottier, P. J., & de Haan, C. A. (2008). Topology and membrane anchoring of the coronavirus replication complex: not all hydrophobic domains of nsp3 and nsp6 are membrane spanning. *J Virol, 82*(24), 12392-12405. doi:10.1128/JVI.01219-08
Padhan, K., Tanwar, C., Hussain, A., Hui, P. Y., Lee, M. Y., Cheung, C. Y., . . . Jameel, S. (2007). Severe acute respiratory syndrome coronavirus Orf3a protein interacts with caveolin. *J Gen Virol, 88*(Pt 11), 3067-3077. doi:10.1099/vir.0.82856-0
Ren, Y., Shu, T., Wu, D., Mu, J., Wang, C., Huang, M., . . . Zhou, X. (2020). The ORF3a protein of SARS-CoV-2 induces apoptosis in cells. *Cell Mol Immunol, 17*(8), 881-883. doi:10.1038/s41423-020-0485-9
Sakai, Y., Kawachi, K., Terada, Y., Omori, H., Matsuura, Y., & Kamitani, W. (2017). Two-amino acids change in the nsp4 of SARS coronavirus abolishes viral replication. *Virology, 510*, 165-174. doi:10.1016/j.virol.2017.07.019
Schaecher, S. R., Diamond, M. S., & Pekosz, A. (2008). The transmembrane domain of the severe acute respiratory syndrome coronavirus ORF7b protein is necessary and sufficient for its retention in the Golgi complex. *J Virol, 82*(19), 9477-9491. doi:10.1128/JVI.00784-08
Schaecher, S. R., Mackenzie, J. M., & Pekosz, A. (2007). The ORF7b protein of severe acute respiratory syndrome coronavirus (SARS-CoV) is expressed in virus-infected cells and incorporated into SARS-CoV particles. *J Virol, 81*(2), 718-731. doi:10.1128/JVI.01691-06
Schuller, M., Correy, G. J., Gahbauer, S., Fearon, D., Wu, T., Díaz, R. E., . . . Ahel, I. (2020). Fragment Binding to the Nsp3 Macrodomain of SARS-CoV-2 Identified Through Crystallographic Screening and Computational Docking. *bioRxiv*, 2020.2011.2024.393405. doi:10.1101/2020.11.24.393405
Serrano, P., Johnson, M. A., Almeida, M. S., Horst, R., Herrmann, T., Joseph, J. S., . . . Wuthrich, K. (2007). Nuclear magnetic resonance structure of the

N-terminal domain of nonstructural protein 3 from the severe acute respiratory syndrome coronavirus. *J Virol, 81*(21), 12049-12060. doi:10.1128/JVI.00969-07

Serrano, P., Johnson, M. A., Chatterjee, A., Neuman, B. W., Joseph, J. S., Buchmeier, M. J., . . . Wuthrich, K. (2009). Nuclear magnetic resonance structure of the nucleic acid-binding domain of severe acute respiratory syndrome coronavirus nonstructural protein 3. *J Virol, 83*(24), 12998-13008. doi:10.1128/JVI.01253-09

Snijder, E. J., Bredenbeek, P. J., Dobbe, J. C., Thiel, V., Ziebuhr, J., Poon, L. L., . . . Gorbalenya, A. E. (2003). Unique and conserved features of genome and proteome of SARS-coronavirus, an early split-off from the coronavirus group 2 lineage. *J Mol Biol, 331*(5), 991-1004. doi:10.1016/s0022-2836(03)00865-9

Su, Y. C. F., Anderson, D. E., Young, B. E., Linster, M., Zhu, F., Jayakumar, J., . . . Smith, G. J. D. (2020). Discovery and Genomic Characterization of a 382-Nucleotide Deletion in ORF7b and ORF8 during the Early Evolution of SARS-CoV-2. *MBio, 11*(4). doi:10.1128/mBio.01610-20

Sutton, G., Fry, E., Carter, L., Sainsbury, S., Walter, T., Nettleship, J., . . . Stuart, D. I. (2004). The nsp9 replicase protein of SARS-coronavirus, structure and functional insights. *Structure, 12*(2), 341-353. doi:10.1016/j.str.2004.01.016

Tan, J., Vonrhein, C., Smart, O. S., Bricogne, G., Bollati, M., Kusov, Y., . . . Hilgenfeld, R. (2009). The SARS-unique domain (SUD) of SARS coronavirus contains two macrodomains that bind G-quadruplexes. *PLoS Pathog, 5*(5), e1000428. doi:10.1371/journal.ppat.1000428

Tanaka, T., Kamitani, W., DeDiego, M. L., Enjuanes, L., & Matsuura, Y. (2012). Severe acute respiratory syndrome coronavirus nsp1 facilitates efficient propagation in cells through a specific translational shutoff of host mRNA. *J Virol, 86*(20), 11128-11137. doi:10.1128/JVI.01700-12

Theobald, D. L., Mitton-Fry, R. M., & Wuttke, D. S. (2003). Nucleic acid recognition by OB-fold proteins. *Annu Rev Biophys Biomol Struct, 32*, 115-133. doi:10.1146/annurev.biophys.32.110601.142506

Thoms, M., Buschauer, R., Ameismeier, M., Koepke, L., Denk, T., Hirschenberger, M., . . . Beckmann, R. (2020). Structural basis for translational shutdown and immune evasion by the Nsp1 protein of SARS-CoV-2. *Science, 369*(6508), 1249-1255. doi:10.1126/science.abc8665

Vasilenko, N., Moshynskyy, I., & Zakhartchouk, A. (2010). SARS coronavirus protein 7a interacts with human Ap4A-hydrolase. *Virol J, 7*, 31. doi:10.1186/1743-422X-7-31

Xu, X., Lou, Z., Ma, Y., Chen, X., Yang, Z., Tong, X., . . . Rao, Z. (2009). Crystal structure of the C-terminal cytoplasmic domain of non-structural protein 4 from mouse hepatitis virus A59. *PLoS ONE, 4*(7), e6217. doi:10.1371/journal.pone.0006217

Yoshimoto, F. K. (2020). The Proteins of Severe Acute Respiratory Syndrome Coronavirus-2 (SARS CoV-2 or n-COV19), the Cause of COVID-19. *Protein J, 39*(3), 198-216. doi:10.1007/s10930-020-09901-4

Yuan, X., Wu, J., Shan, Y., Yao, Z., Dong, B., Chen, B., . . . Cong, Y. (2006). SARS coronavirus 7a protein blocks cell cycle progression at G0/G1 phase via the cyclin D3/pRb pathway. *Virology, 346*(1), 74-85. doi:10.1016/j.virol.2005.10.015

Yuen, C. K., Lam, J. Y., Wong, W. M., Mak, L. F., Wang, X., Chu, H., . . . Kok, K. H. (2020). SARS-CoV-2 nsp13, nsp14, nsp15 and orf6 function as potent interferon antagonists. *Emerg Microbes Infect, 9*(1), 1418-1428. doi:10.1080/22221751.2020.1780953

Zhang, Y., Zhang, J., Chen, Y., Luo, B., Yuan, Y., Huang, F., . . . Zhang, H. (2020). The ORF8 Protein of SARS-CoV-2 Mediates Immune Evasion through Potently Downregulating MHC-I. *bioRxiv*, 2020.2005.2024.111823. doi:10.1101/2020.05.24.111823
